# Extended amygdala function is governed by threat probability, not uncertainty

**DOI:** 10.64898/2026.09.28.754981

**Authors:** Mesude Okhan, Kalina N. Kalinova, Kathryn A. DeYoung, Paige R. Didier, Zachary S. Scott, Dominique B. Ewing, Jordan Hassani, Jennifer V. Cunneen, Elisa Liu, Julia K. Earp, Gaby B. Castro, Jonathan A. Oler, Patrick H. Roseboom, Connor W. Christensen, Carly M. Drzewiecki, Jairo Chavez, Erie D. Boorman, Jason F. Smith, Ned H. Kalin, Alexander J. Shackman, Andrew S. Fox

## Abstract

Anxiety is a universal experience that arises as we anticipate negative events that may or may not happen. When extreme, anxiety is a major cause of distress, suffering, and disability-adjusted life years, with nearly one in four individuals experiencing clinically relevant anxiety in their lifetime^1–3^. Elucidating the biological basis of anxiety has the potential to help people understand, cope with, and ultimately receive improved treatments for their anxiety. The canonical neuroscientific model of anxiety posits that distinct regions of the extended amygdala (EA) elicit distress that is conditional on threat uncertainty^4^. Here, we refute this model using a precise computational framework to disambiguate specific dimensional facets of threat uncertainty in a racially diverse sample of 104 individuals. We show that self-report, psychophysiological, and neural correlates of threat anticipation are much more strongly associated with threat probability than uncertainty *per se*. Importantly, EA regions, including the central nucleus of the amygdala (Ce) and bed nucleus of the stria terminalis (BST), were most reliably associated with threat probability—contrary to the canonical model. Re-examination of the pioneering experiments that motivated the canonical model reveal a previously unappreciated confound between probability and uncertainty, suggesting the novel hypothesis that EA regions differ in their sensitivity to low-probability threats. Our fMRI data support this, showing that BST is more sensitive to low-probability threats than Ce. Moreover, lesion experiments in rhesus monkeys (*Macaca mulatta*) demonstrate that BST causally contributes to defensive behavior in low-probability threat contexts. Together, these data reconcile previous empirical inconsistencies, provide a fresh theoretical perspective on the neurobiology of anxiety, and underscore the theoretical importance of a formal *lingua franca* for coordinated cross-species emotion research.

## Introduction

Threat certainty plays a central role in contemporary theories of anxiety^4–6^. It is widely believed that "certain" and "uncertain" threats elicit distinct forms of distress that arise from strictly segregated circuits within the extended amygdala (EA), which encompasses the central nucleus of the amygdala (Ce) and bed nucleus of the stria terminalis (BST)^4,7,8^. This canonical model posits that the Ce triggers phasic responses to threats that are certain, acute, or imminent; whereas the BST triggers sustained responses to threats that are uncertain, distal, or diffuse^8,9^. This model has had an enormous impact on basic, translational, and clinical research, and is embedded within the U.S. National Institute of Mental Health’s influential Research Domain Criteria (RDoC) framework^6,8,9^.

The canonical model originated from pioneering work demonstrating that targeted manipulations of Ce and BST have doubly dissociable effects on defensive behavior. Despite their many neuroanatomical similarities (e.g. cell-types and connectivity), manipulations of Ce and BST dissociably affected defensive behavior to threats often characterized as "certain" or "uncertain", respectively. For example, early work by Davis, Walker, and colleagues demonstrated that Ce inactivation selectively decreases "fear-potentiated" startle during "acute" Pavlovian threat cues, whereas BST inactivation selectively blocks "light-potentiated" startle to the "diffuse" threat signaled by a brightly illuminated chamber in nocturnal rodents^10^. Subsequent work demonstrated Ce-BST dissociations in other paradigms, including contextual conditioning at short-vs-long intervals^11,12^, cued-vs-uncued shocks^4^, and forward-vs-backward threat conditioning^13^. Moreover, some human neuroimaging research has revealed parallel differences in Ce-BST fMRI activation during certain-vs-uncertain threat anticipation^e.g.14,15^. Together, these results have been interpreted as strong evidence for the canonical model.

Nevertheless, an expanding body of research has failed to support the canonical model’s predictions in relation to "certain" and "uncertain" threat processing, suggesting that the canonical model is either incomplete or incorrect^16–19^. In rodents, manipulations of Ce and BST can each alter responses to both imminent and diffuse threats^20,21^. In human fMRI studies, certain and uncertain threats are associated with a highly overlapping neural circuit, including the Ce and BST^22,23^. However, the impact of these findings has been limited, leading to a theoretical impasse exacerbated by inconsistent and amorphous definitions of uncertainty and a reliance on oversimplified categorical manipulations^24,25^. Computational approaches provide a path to overcome these barriers and develop viable alternative models that can account for these seemingly disparate observations.

Here, we used a computational framework to disentangle specific dimensional facets of threat uncertainty from threat likelihood ("Probability") in 104 racially diverse participants (see Methods and data exclusions). By leveraging behavioral-economic theory, we defined two mathematically distinct facets of threat "uncertainty": outcome uncertainty ("Risk") and probability uncertainty ("Ambiguity"). "Risk" quantifies the variance in potential outcomes (aversive or benign; i.e. *Var[p_Threat_]*). For example, a cue signaling an aversive outcome half the time is *riskier* than one that always leads to an aversive outcome— certain threat cues are not risky, they are just bad. "Ambiguity" quantifies the extent to which threat probability is partially unknowable. Ambiguous threats encompass a range of potential probabilities (*range[p_Threat_]*), making it impossible to know if an aversive outcome is likely or not. Combined with an affective-neuroscience approach, these precise definitions allowed us to separate the consequences of threat Probability, Risk, and Ambiguity on subjective distress, objective psychophysiological arousal, and brain function. Contrary to the canonical model, self-reported distress, skin conductance responses (SCR), and the BOLD fMRI signal in threat-sensitive brain regions (including Ce & BST) most closely track threat Probability, rather than Risk or Ambiguity. Instead, these data support a revised model of EA function, which we evaluated across human neuroimaging and non-human primate lesion data. This model reconciles previous inconsistencies, provides fresh insights into the neuroanatomy of fear and anxiety, and underscores the utility of theory-driven computational approaches for emotion research.

## Results

We developed a randomized event-related fMRI task, the Maryland-Davis Ambiguous and Risky Threat (M-DART) paradigm, to dissociate distinct facets of uncertainty. All trials began with the presentation of a cue and terminated with a compound aversive or benign outcome, consisting of an image, audio clip, and electrical stimulation (**Fig. 1a**). Cues indicated threat likelihood (0-100% in 10% increments), allowing us to model parametric variation in threat Probability (i.e. *likelihood* of aversive outcome) and Risk (i.e. *variability* in aversive outcome)(**Fig. 1c**). On 50/136 trials, the cue is partially obscured by a mask (10-90% in 10% increments), rendering threat probability partially unknowable and enabling us to model variation in threat Ambiguity (i.e. *range of potential* probabilities) (**Fig. 1d**). On 104/136 of inter-trial intervals, subjects were prompted to rate the intensity of distress experienced during the preceding cue. SCR and fMRI were continuously collected throughout.

**Fig. 1.**
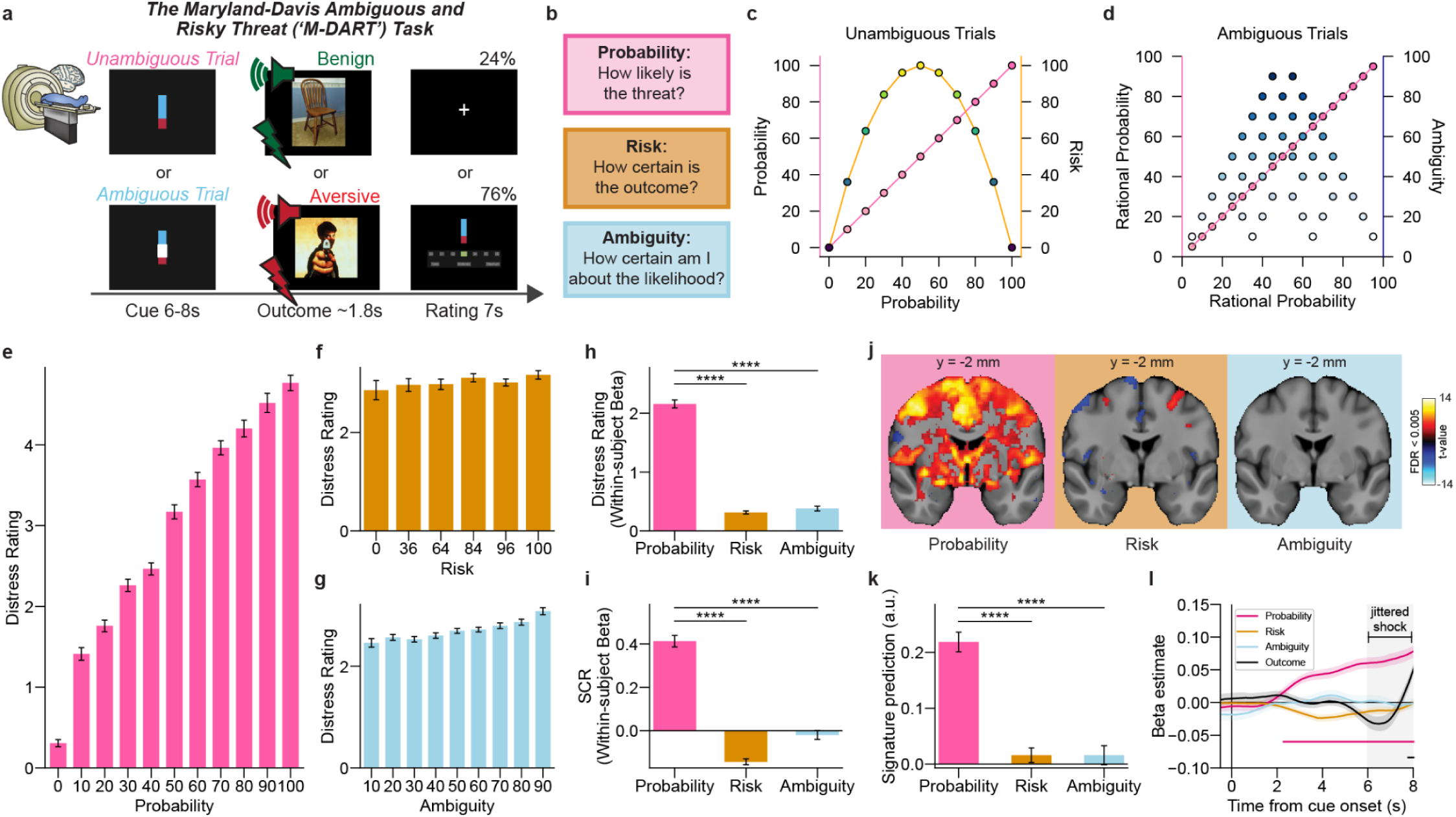
Probability of threat predicts self-reported distress, skin conductance, and brain activation better than Risk and Ambiguity. a, Schematic of the Maryland-Davis Ambiguous and Risky Threat (M-DART) task. Unambiguous and Ambiguous trials consisted of a visual cue signaling threat probability (6–8 s), a benign or aversive outcome (∼1.8 s), and a distress rating (7 s) on 76% of trials. b, Three task dimensions dissociated in the M-DART paradigm. Probability captures threat likelihood; Risk and Ambiguity represent two mathematically and conceptually distinct facets of threat uncertainty. c-d, Demonstration of independent, orthogonalized cues for Risk (c) and Ambiguity (d) trials. On Risk trials (no ambiguity), probability increases linearly (pink) while Risk follows an inverted-U peaking at 50% (orange). On Ambiguity trials, rational Probability varies linearly (pink) while Ambiguity varies independently (blue) (see Method for details). e–g, Distress ratings across threat Probability (e), Risk (f), and Ambiguity (g) levels (N = 104). Distress increased monotonically with threat Probability, Risk, and Ambiguity (linear-mixed-effects model; Probability: z = 144.09, Risk: z = 21.73, Ambiguity: z = 14.83, all p < 0.0001). h,i, Within-subject beta estimates for distress ratings (h; N = 104) and skin conductance response (SCR; i; N = 102) as a function of Probability, Risk, and Ambiguity. Probability significantly predicted both distress ratings and SCR (linear mixed-effects model; distress: z = 44.69; SCR: z = 19.82; both p < 0.0001). Probability also elicited significantly larger effects than Risk and Ambiguity in both measures (distress: Risk, z = −27.03; Ambiguity, z = −26.06; SCR: Risk, z = −18.89; Ambiguity, z = −14.68; all p < 0.0001). j, Whole-brain activation for Probability, Risk, and Ambiguity contrasts (q < 0.005, FDR-corrected; N=101). Coronal slice at MNI y = -2 mm. Color scale, t-values. k, The prediction for the negative affect signature^26^ as a function of Probability, Risk, and Ambiguity (N = 101). Signature prediction was significantly greater for Probability than for Risk or Ambiguity (linear mixed-effects model, Probability as reference level: Risk z = −8.96, Ambiguity z = −8.95, both p < 0.0001). l, Mean timecourse of beta estimates for negative affect signature^26^ predictions as a function of Probability, Risk, Ambiguity, and Outcome (shock vs. no shock), averaged across N = 101 subjects. By simultaneously estimating the effect of Probability, Risk, Ambiguity, and Outcome (aversive or benign) at each time-point, we found only Probability to be significantly impacted during the cue period. Specifically, Probability-driven modulation of the negative affect signature during cue anticipation was dissociable from responses to aversive outcomes. Horizontal lines above the x-axis mark timepoints at which the beta estimate was significantly greater than zero (two-sided one-sample t-tests, FDR q < 0.05 across predictors and timepoints); line colors match the corresponding predictors. Time is aligned to cue onset (0 s) and sampled every 100 ms (81 timepoints per line, 0–8 s window); shaded interval (6–8 s) denotes the jittered outcome period. Confidence envelopes, SEM. Error bars, SEM. All statistical tests were two-sided. ****p < 0.0001. a.u., arbitrary units.

As a precursor to hypothesis testing, we confirmed that the threat manipulation had the intended impact on subjective distress, objective psychophysiological arousal, and brain function. Relative to "Certain-Safety" (0% probability; 0% ambiguity), "Certain-Threat" cues (i.e. 100% probability; 0% ambiguity) were associated with significantly increased self-reported distress and SCR (distress: z=130.35; SCR: z=24.96, both p<0.0001; **Supplementary Fig. 1a-b;** full statistics for all tests are provided in **Supplementary Table 1**). Certain-Threat cues also recruited a distributed network of brain regions previously implicated in fear and anxiety, including Ce and BST (q<0.005 FDR-corrected; **Supplementary Fig. 1c & 2; Supplementary Table 2**). Finally, established whole-brain multi-voxel signatures confirmed that Certain-Threat cues were associated with higher levels of signature-predicted negative affect^26^ and anticipatory anxiety that were not accounted for by uncertainty^27^ (**Supplementary Fig. 1d-e; Supplementary Fig. 3; Supplementary Table 3**). Thus, the threat manipulation worked as intended.

Next, we used trial-by-trial parametrically modulated regressors to model the unique contribution of threat Probability, Risk, and Ambiguity on self-reported distress, SCR, and BOLD signals (see Methods). Results demonstrated that both distress and SCR significantly covaried with threat Probability (distress: z = 44.69; SCR: z = 19.82; both p < 0.0001; **Fig. 1h,i**). Moreover, the effect of Probability was markedly stronger than either Risk or Ambiguity (distress: Probability-vs-Risk, z=−27.03; Probability-vs-Ambiguity, z=−26.06; SCR: Probability-vs-Risk, z=−18.89; Probability-vs-Ambiguity, z=−14.68; all p<0.0001). A similar pattern was evident in whole-brain voxelwise analyses, where activation magnitude strongly covaried with threat Probability in regions previously associated with fear and anxiety, including Ce, BST, hypothalamus, periaqueductal gray (PAG), midcingulate cortex (MCC), and anterior insula (AI) (q<0.005, FDR-corrected; **Fig. 1j**; **Supplementary Fig. 4; Supplementary Table 4**). Qualitatively, the Probability effect was much stronger than those of Risk or Ambiguity, which were associated with comparatively sparse activation (**Fig. 1j**; **Supplementary Note**, **Supplementary Fig. 4**; **Supplementary Table 4-6**). To statistically compare the global effects of threat Probability, Risk, and Ambiguity, we again leveraged the whole-brain neural signatures. Signature-predicted negative affect and anticipatory anxiety were much more strongly associated with variation in threat Probability relative to Risk and Ambiguity (negative affect: Risk, z=-8.96; Ambiguity, z=−8.95; **Fig. 1k**; anticipatory anxiety: Risk, z=−13.06; Ambiguity, z=−14.40; **Supplementary Fig. 1f**; all p<0.0001). Critically, moment-by-moment regression analyses demonstrated that the impact of Probability on the anticipatory BOLD signal during the cue cannot be attributed to an artifact of aversive stimulation (**Fig. 1l**; see Methods). Collectively, these data show that the major contributor to anticipatory distress, psychophysiological arousal, and brain activation is the likelihood of an aversive outcome, *not* outcome uncertainty or unknown probabilities.

To test the canonical model, we extracted the estimated effects of Probability, Risk and Ambiguity from Ce and BST anatomical ROIs. Both EA regions were significantly modulated by threat Probability (Ce: z=7.00; BST: z=6.22; both p<0.0001), not Risk or Ambiguity (Risk: Ce, z=-1.13, p_adj_=0.287; BST, z=1.64, p_adj_=0.169; Ambiguity: Ce, z=-1.30, p_adj_=0.270; BST, z=0.40, p_adj_=0.686). The Probability effect was significantly stronger than either Risk or Ambiguity in both regions (Ce-Risk-vs-Probability: z=−5.91; Ce-Ambiguity-vs-Probability: z=−6.03; BST-Risk-vs-Probability: z=-3.33; BST-Ambiguity-vs-Probability: z=-4.23, all p<0.0001, **Fig. 2a; Supplementary Fig. 5**). There was no significant Region (Ce/BST) × Facet (Probability/Risk or Ambiguity) interaction (Region × Probability-vs-Risk: z=−1.82, p=0.068; Region × Probability-vs-Ambiguity: z=−1.28, p=0.202). Together, these results fail to support the canonical model of EA function, which posits that Ce exclusively mediates responses to certain threats, whereas BST exclusively mediates responses to uncertain threats (**Fig. 2a**).

**Fig. 2.**
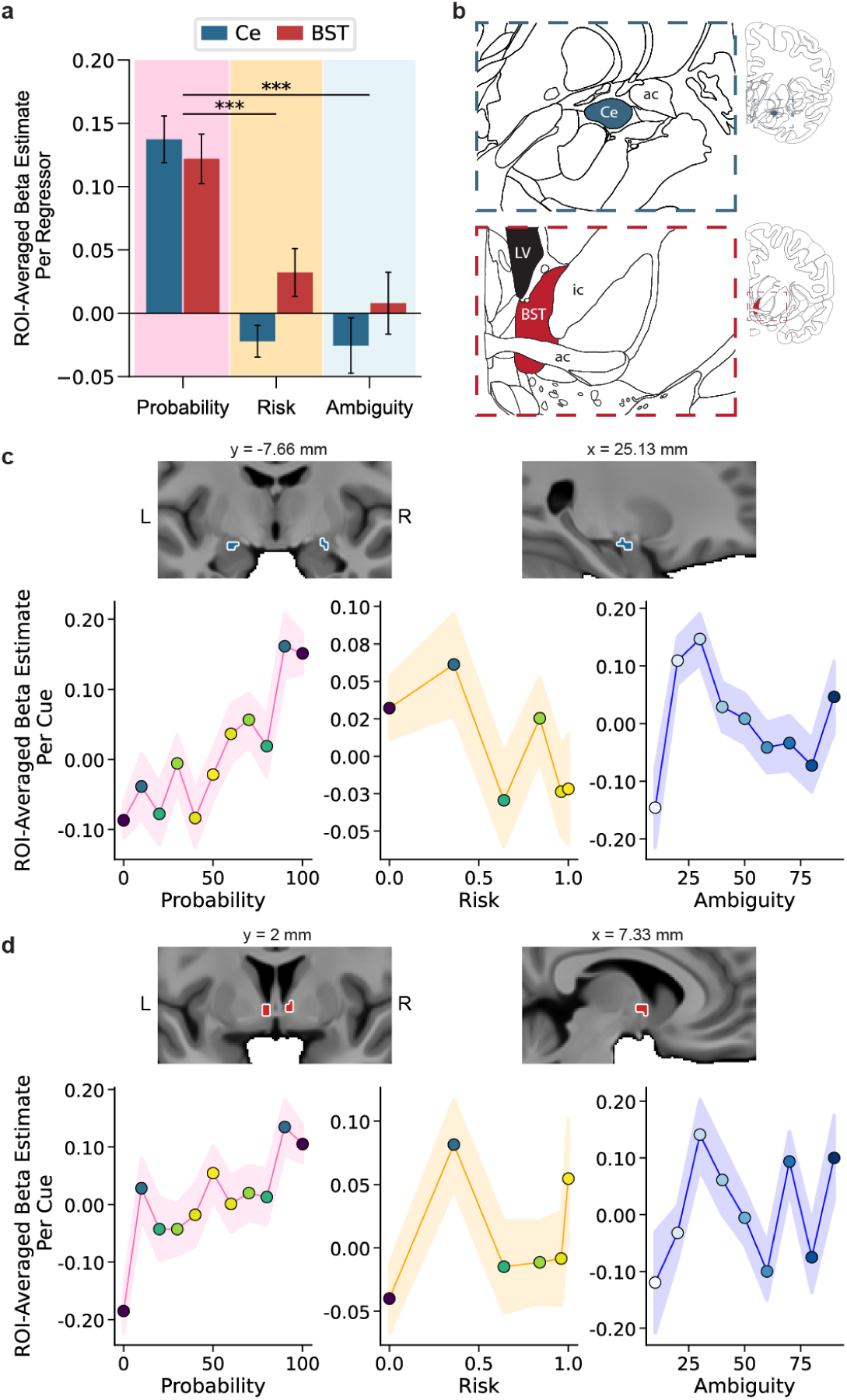
Ce and BST are modulated by Probability, not Risk or Ambiguity. a, Mean ROI beta estimates for Ce and BST across Probability, Risk, and Ambiguity facets (N = 101). Both regions demonstrated a significant effect of Probability (linear mixed-effects contrasts; Ce: z=7.00; BST: z=6.22; both p_adj_<0.0001), not Risk (linear mixed-effects contrasts; Ce; z=-1.13, p_adj_=0.287; BST; z=1.64, p_adj_=0.169) or Ambiguity (linear mixed-effects contrasts; Ce, z=-1.30, p_adj_=0.270; BST, z=0.40, p_adj_=0.686). Both regions showed greater responses to Probability than Risk or Ambiguity (linear mixed-effects model; Risk vs. Probability: Ce: z = −5.91, BST: z = −3.33; Ambiguity vs. Probability: Ce: z = −6.03, BST: z = −4.23; all p < 0.001), with no significant interaction (Region × Probability-vs-Risk: z = −1.82, p = 0.068; Region × Probability-vs-Ambiguity: z = −1.28, p = 0.202). **b,** Anatomical location of Ce (top) and BST (bottom) in the Allen Brain Atlas. ac, anterior commissure; LV, lateral ventricle; ic, internal capsule. **c-d,** (c) Ce ROI displayed on average structural scan (top; MNI y = −7.66 mm and x = 25.13 mm) and visualization of cue-specific mean beta estimates as a function of Probability, Risk, and Ambiguity (bottom; N = 101). (d) The same as c for BST ROI (MNI y = 2 mm and x = 7.33 mm). Error bars and confidence envelopes (a, c, d), SEM. All statistical tests were two-sided. ***p < 0.001

These results stand in contrast to prior work that relied on categorical comparisons of uncertain-threat anticipation to certain-safety, often operationalized as <60% threat-probability vs. safety^e.g.28–30^. Surprisingly, we were able to replicate prior reports of greater BST-vs-Ce reactivity when directly contrasting 50% threat-probability vs. safety (z=−2.95, p=0.003, **Fig. 3a-b; Supplementary Fig. 6; Supplementary Table 7**). In summary, our parametric results challenge a core tenet of the canonical model (e.g. **Fig. 2a**), showing that uncertainty is not the primary facet of threat distinguishing EA regions. Yet paradoxically we were able to replicate neuroimaging results taken as supportive of this model. Together, these results beg the question: *If not uncertainty, then what is driving Ce-BST dissociations?*

**Fig. 3.**
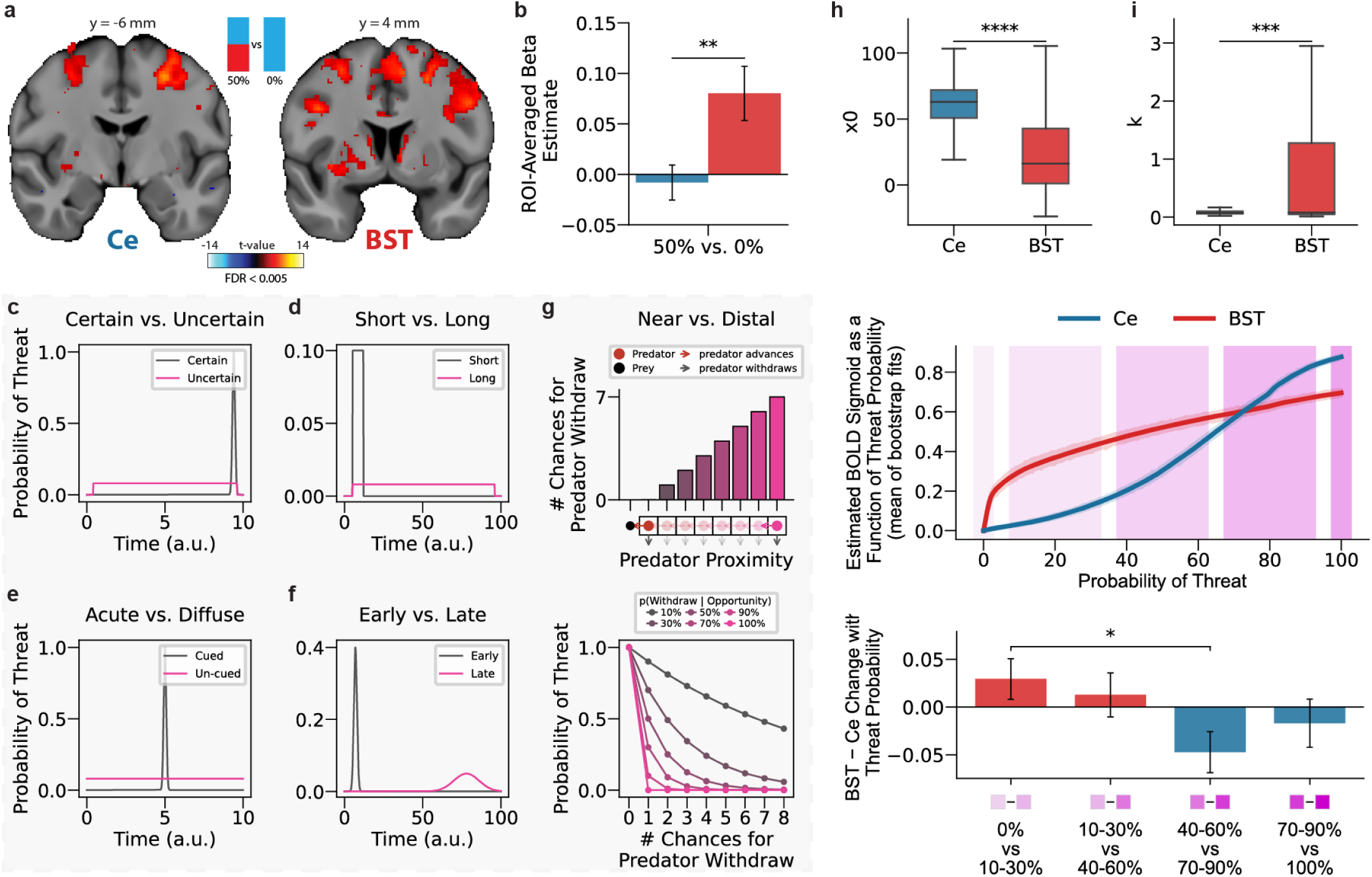
**BST preferentially responds to low-probability threats**. **a**, Whole-brain activation for the 50% vs. 0% threat probability contrast (q < 0.005, FDR-corrected; N = 101) demonstrates a main effect for BST, but not Ce. Inset (top center) depicts the contrasted cue types. Coronal slices at MNI y = -6 mm and y = 4 mm were selected to highlight the Ce (left) and BST (right) ROIs, respectively. Color scale, t-values. **b**, Replication of prior reports of "uncertainty" on Ce-BST dissociations using ROI-averaged beta estimates for the 50% vs. 0% contrast (N = 101). BST showed a significantly greater response to 50% vs. 0% threat probability than Ce (linear mixed-effects model; z = −2.95, p = 0.003). **c–f**, Theoretical illustrations of threat probability across dimensions previously used to dissociate Ce and BST: (**c**) *certain vs. uncertain*: manipulations of temporal uncertainty require that the subject not know when a threat will occur, resulting in a low probability of threat at any given moment, as compared to when subjects know precisely when the negative event will occur (e.g. 10% at each second in a 10-second countdown vs. 100% at the cued time); (**d**) *short vs. long duration*: manipulations of sustained threats often distribute the probability of threat over a longer period of time, which result in the probability of threat at any given moment being lower than if it was distributed over a short period of time (e.g. 1% for 100 seconds vs. 10% for 10 seconds), (**e**) *acute (cued) vs. diffuse (un-cued)*: diffuse threats are characterized by stimuli and contexts with a low-probability of predicting a threat, e.g. testing a nocturnal animal with the light on, reflects a context where the likelihood of threat is present, but low (i.e. <10%); and, (**f**) *early vs. late*: Manipulations of threats that occur early vs. late during a sustained cue might also be different in the perceived probability, as the accuracy of elapsed time becomes less accurate as time passes. As a result, the probability that a negative event will happen at a specific moment may be less certain as more time has elapsed. In each case, the pink condition — regardless of its specific label across paradigms — corresponds to lower threat probability at any given moment. Time axes, arbitrary units. **g**, Theoretical illustration of *near vs. distal threat*. *Top*: An illustrative bar-graph showing that as the distance between the predator and prey gets larger, the predator has more chances to withdraw. The x-axis is predator distance, shown as prey (black dot) on the left, and predator’s (red-light pink dots) of various distances from left (closest) to right (furthest). The bars show the number of chances the predator has to withdraw. *Bottom*: Depiction of how the number of chances for the predator to withdraw impacts the probability of threat. Each curve shows the probability of threat to potential prey, as a function of the number of opportunities a predator has to withdraw. Different lines indicate different probabilities of predator withdraw given they have the opportunity (i.e. p(Withdraw | Opportunity)), with values from 10% to 100% (dark gray to pink). Imminence and many other factors can provide more opportunities for predator withdrawal, reducing effective threat probability. For example, a distal predator would have multiple opportunities to withdraw, but the probability of threat to its potential prey will also depend on the likelihood of the predator withdrawing when it has the chance opportunity (i.e. 1-p(attack)). This provides a conceptual reformulation of distal vs. near threats as a function of threat probability. **h**,**i**, Bootstrapped estimates of sigmoid midpoint (*x_0_*; **h**) and slope (*k*; **i**) for Ce and BST across 1,000 bootstrap iterations (half-sample resampling with replacement, n = 50 drawn from N = 101 subjects). Box plots show the quartiles of parameter estimates across bootstrap permutations. BST exhibited a significantly lower midpoint and steeper slope than Ce (Mann-Whitney U:*x_0_*, U = 193,918, p < 0.0001; *k*, U = 543,695, p = 0.0007), indicating that BST activation rises at lower threat probabilities and transitions more sharply than Ce. **j**, Visualization of mean bootstrapped logistic BOLD sigmoid curve fits for Ce and BST as a function of threat probability. Shaded envelopes show the 99.9% CI of the mean curve across bootstrap iterations (precision of the average; variability across fits is shown in **h,i**); background shading delineates the probability bins compared in **k**. **k**, Change in BST–Ce estimated BOLD in adjacent probability bins, when the number of trials in each bin is matched. Results support the bootstrapping analysis, showing greater BST–Ce change in low-probability bins. A one-way ANOVA on the BST–Ce difference scores showed a marginal main effect of probability-bin transition (F(3,400) = 2.61, p = 0.051). Planned pairwise comparisons of the lowest-probability transition (0% vs. 10–30%) against the other three transitions (Holm-corrected) showed a significant difference from the 70–90% vs. 40–60% transition (z = 2.59, p_adj_ = 0.029), but not from the 40–60% vs. 10–30% (z = 0.56, p_adj_ = 0.574) or 100% vs. 70–90% (z = 1.57, p_adj_ = 0.234) transitions. Error bars (b, k), SEM. All statistical tests were two-sided. *p < 0.05, **p < 0.01, ***p < 0.001, ****p < 0.0001. a.u., arbitrary units.

To address this paradox, we re-examined the foundational Ce-BST dissociation research that motivated the canonical model, searching for statistical confounds that might better account for effects heretofore attributed to threat "uncertainty." In both human and animal studies, manipulations of threat uncertainty (e.g. temporally certain vs. uncertain, acute vs. diffuse, etc.) were nearly always confounded with threat *probability* (**Fig. 3c-g**). For instance, during exposure to "diffuse" threats (e.g. "light-potentiated startle"), the likelihood of threat at any given moment is lower compared to an "acute" shock-predicting cue. This insight motivates the hypothesis that a core EA function is to track threat likelihood, and that BST may be more sensitive to low-probability threats than Ce.

To test this, we modeled Ce and BST BOLD activation using a logistic function of threat probability. Leveraging a between-subjects bootstrapped model, results revealed BST fits to be significantly steeper and left-shifted compared to Ce (**Fig. 3h-j**; Mann-Whitney U: *x_0_*, U = 193,918, p < 0.0001; *k*, U = 543,695, p = 0.0007). Thus, although Ce and BST both track threat probability, BST is more sensitive to low-probability threats, whereas Ce predominantly responds to moderate/high-probability threats. This conclusion was also evident when performing targeted comparisons of Ce-vs-BST change in activation between adjacent probability "bins" (F(3,400)=2.61, p=0.051; **Fig. 3k**; see Method for details). Taken with the probability-uncertainty confounds in prior work, these results form the basis for a refined model relating EA function to anticipated threats. Consistent with a voluminous literature, our model suggests that Ce drives behavioral responses to high-probability threats^31^ (though also see:^32,33^). In contrast, our model predicts that BST is particularly important for driving behavioral responses to low-probability threats, not uncertain ones.

Because we cannot perform targeted manipulations of human BST, we used a variant of the human intruder paradigm (HIP-Persistence) to test the hypothesis that BST is necessary for mounting defensive responses to low probability threat contexts in another primate species, rhesus monkeys. Rhesus monkeys are an ideal species to begin the process of reverse translation because of their evolutionary closeness to humans, which manifests as close homologies in neurobiology and socio-emotional behavior. HIP-Persistence includes two distinct "uncertain" threat contexts: one where a human intruder presents their profile while avoiding eye-contact (no eye-contact; NEC) and a second encompassing the period after the intruder leaves (alone; ALN-following-NEC)^34^. Both contexts are "uncertain," insofar as it is unclear if an intruder will approach, return, or attack; and both elicit freezing, a defensive behavior rarely evoked by a certain threat (e.g. a staring intruder)^34–36^. This mirrors real-world scenarios where individuals show signs of fear and anxiety when a predator is present but has not detected the prey, and persistent trepidation after the predator departs. Despite their similarities, the two contexts differ in threat probability, which is moderate/high during NEC and much lower in ALN-following-NEC. Past work demonstrates that Ce lesions decrease freezing during NEC^36^. Here, we tested competing predictions about the impact of BST lesions across threat contexts. Because NEC and ALN-following-NEC are both uncertain, the canonical model predicts decreased freezing across contexts. In contrast, the threat-probability model predicts a selective decrease during the low-probability ALN-following-NEC context. To adjudicate these accounts, we performed MRI-guided ibotenic-acid lesions of the BST-region and assessed freezing during 30-min exposures to each context (**Fig. 4a-d**; see methods). Results revealed a significant lesion by context interaction (z=2.15, p=0.031): lesioned animals froze significantly less than unoperated controls in the ALN-following-NEC context (z=- 3.14, p=0.002), but groups did not differ in the NEC context (z=-0.10, p=0.920; **Fig. 4e**). The reduction in ALN-following-NEC freezing remained significant when the lesion group was restricted to the animals with bilateral BST overlap (z = −6.16, p < 0.0001; **Fig. 4e,f**; also see Methods). No significant lesion by condition interaction was present for locomotion, demonstrating that gross motor deficits are insufficient to account for the observed findings (z=1.37, p=0.171; also see **Supplementary Fig. 7**). To clarify the larger significance of BST damage, we assessed freezing during a naturalistic encounter with an unfamiliar same-sex conspecific (**Fig. 4h**). Because prior work has interpreted anxiety while interacting with novel peers as resulting from the intrinsic uncertainty^37^, we aimed to test the hypothesis that our BST-lesions, which alter response to low-probability but not uncertain threats, would be sufficient to reduce defensive behaviors during novel social interactions. Results demonstrated that lesioned animals froze less than controls during a single-exposure social interaction (z=-4.14, p <0.0001; **Fig. 4i).** Together, these data fail to support the canonical model, show that BST is crucial for mounting adaptive emotional responses to low-probability threats, but not higher-probability threats which are known to depend on Ce, and hint at the significance of this low-probability-threat system for shyness and social anxiety.

**Fig. 4.**
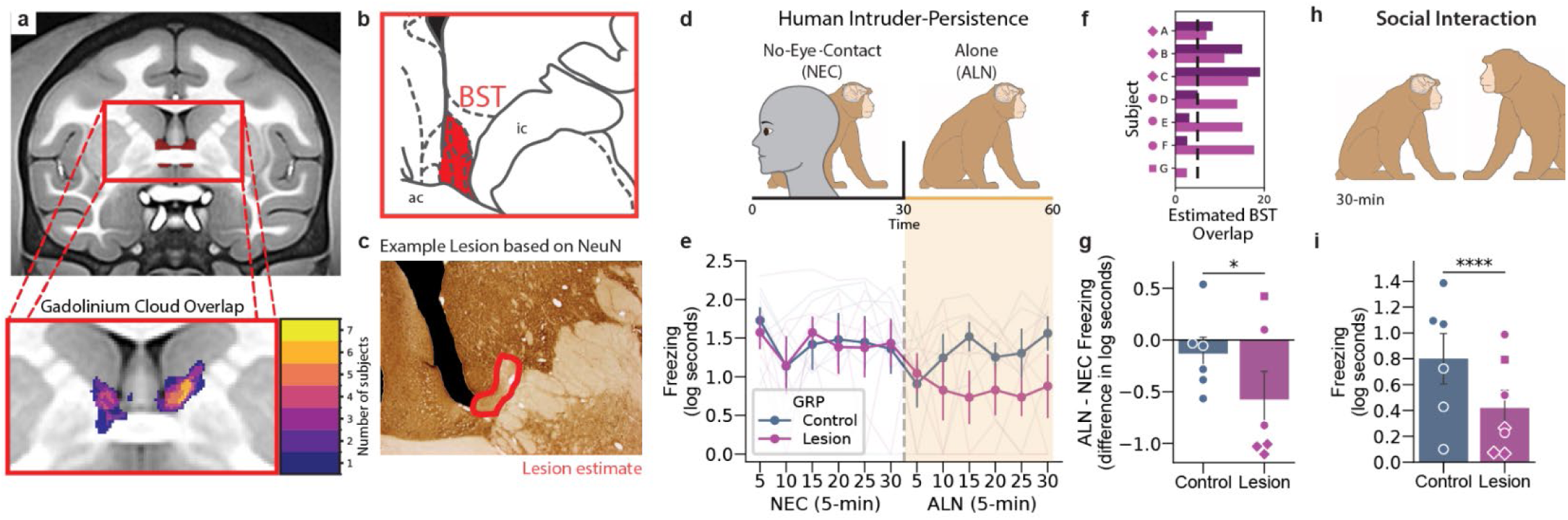
BST lesions in NHPs selectively reduce freezing during low, but not high, threat probability contexts. **a**, BST ROI in NHP NMT-template space (top), zoomed to show the voxelwise overlap of gadolinium-visualized infusion clouds across the experimental group (N = 7; BST-Lesion Group), colored by the number of subjects with coverage at each voxel (bottom). **b**, Anatomical location of BST in the Paxinos Rhesus Monkey Brain atlas. ac, anterior commissure; ic, internal capsule. **c**, Example ibotenic acid-induced BST lesion, with the lesion boundary (red) estimated from loss of NeuN immunostaining. **d**, Schematic of the HIP-Persistence paradigm, depicting the 30-min No-Eye-Contact (NEC) period followed by the 30-min Alone (ALN) period. The animal is unrestrained and freely moves within the enclosure throughout. **e**, Mean freezing across 5-min bins during NEC and ALN portions of the HIP-Persistence paradigm in BST-lesioned (magenta, N = 6) and control (blue, N = 6) animals; thin lines show individual animals. Freezing differed between groups depending on condition (linear mixed-effects model; group × condition interaction: z = 2.15, p = 0.031). Simple effects derived from linear contrasts of the fitted model’s coefficients (Wald z-tests) showed that lesioned animals froze significantly less than controls in the ALN context (z = −3.14, p = 0.002), but not in the NEC context (z = −0.10, p = 0.920). **f**, Estimated BST damage for each subject, shown separately for each infusion. Diamond, bilateral overlap with BST; circle, unilateral overlap; square, no strong overlap. The dashed line indicates threshold for strong BST overlap. **g**, Average ALN−NEC difference in freezing during the HIP-Persistence paradigm in BST-lesioned (magenta) and control (blue) groups; individual animals shown with markers matching **f**. Analyses are based on the same linear mixed-effects model as Fig. 4e (group × condition interaction: z = 2.15, p = 0.031). Difference scores remained trend-level significant between groups across the full sample when using a linear model (t(5) = −2.29, p = 0.071), and this analysis remained significant when restricted to animals with the most bilateral BST overlap (linear model; t(2) = −5.74, p = 0.029). **h**, Schematic of the Social Interaction paradigm, in which the target animal freely interacts with an older, larger, dominant conspecific (stimulus animal) for 30 min. **i**, Mean freezing during the 30-min Social Interaction paradigm in BST-lesioned (magenta, N = 7) and control (blue, N = 6) animals; individual animals shown with markers matching **f**. Freezing differed significantly between groups across the full sample (linear mixed-effects model; group effect: z = −4.14, p < 0.0001), and this difference remained when restricted to animals with confirmed bilateral BST overlap (z = −4.35, p < 0.0001, N = 3), reflecting reduced freezing in BST-lesioned animals during direct interaction with a dominant conspecific. Error bars (e, g, i), SEM. All statistical tests were two-sided. *p < 0.05; **p < 0.01; ****p < 0.0001.

## Discussion

For the past 15 years, the canonical model has been the dominant perspective on the neurobiology of typical and pathological fear and anxiety and has motivated rodent, NHP, and human research focused on threat “uncertainty.” The present results support a refined perspective—one rooted in a precisely formulated computational framework—that can reconcile theoretically troubling inconsistencies in the evidentiary record. Across species and anxiety-related readouts—self-reported feelings, physiological arousal, brain function, and defensive behavior—we failed to find support for the canonical model’s prediction that uncertainty *per se* drives threat responses. In contrast to the canonical model’s central claim, Ce and BST were both modulated by threat Probability, *not* Risk or Ambiguity; begging the question: *if not uncertainty, then what?* Using a computational framework to reinterpret past work, we identified a systematic confound between threat probability and "uncertainty" manipulations. Based on this, we demonstrate that low-probability threats differentially modulate Ce and BST activation: low-probability threats were specifically associated with elevated BST BOLD. Finally, we leveraged an NHP model to demonstrate BST’s causal contribution to defensive responding in low-probability threat contexts. Together these results suggest a new model for how human brains process threats, with implications for both preclinical animal studies and human clinical research. The beauty of the canonical model stemmed from its conceptual unification and integration of seemingly disparate results. Building on this foundation, our alternative model addresses a crucial limitation of the canonical model, providing increased explanatory scope and predictive power across species and varied threat assays. Together, these findings reconcile prior empirical findings, ground the study of fear and anxiety in an ethological and computational framework, and provide concrete next steps for basic and clinical research.

### Reconciling prior empirical findings

Our new model provides insight into the specific facet of threat that drives anxiety-like behavior in a variety of Ce-BST dissociating paradigms previously unified under the banner of "uncertain" threats^4,38^. Whether focused on uncertain, diffuse, or distal threats, the "uncertain" cues and contexts were invariably low probability. To be concrete, uncertain cues are by definition less likely to culminate in harm (e.g. 50%) than certain ones (100%) or less likely at any given moment (e.g. temporal uncertainty); diffuse threats are defined by stimuli that predict threat with a low probability (e.g. dark room); and distal threats necessarily provide opportunity for avoidance and alternative outcomes (e.g. predator withdraws), lowering the probability of harm (**Fig 3c-g**). Put differently, all low-probability threats are uncertain, but not all uncertain threats are low-probability. Consider two threats with a 10% and 90% probability of occurring. Both outcomes are uncertain (may or may not happen, with identical Risk), but only the 10% threat is low-probability. Unlike the canonical model, our new framework puts the emphasis squarely on low-probability threat.

### Grounding in an ethologically relevant framework

In principle, having distinct circuits for processing low-vs-high probability threats enhances the organism’s adaptive capacity. Our neuroimaging results show a gradient of anticipatory changes in brain activity as threat probability increases, which could enable adaptive calibration of defensive and appetitive behaviors to maximize fitness. This aligns with the ethological observations that motivated the influential Predator Imminence Continuum (PIC) model. In naturalistic settings, with meaningful avoidance-approach tradeoffs, the adaptive response to low-probability threat is to increase wariness and vigilance. As threat probability increases, it becomes adaptive to engage in more costly responses, including passive and active defensive behaviors (e.g. freezing or withdrawal). Here, we provide a computationally grounded, unified account for *why* Ce and BST are preferentially involved in driving responses to high-and low-imminence threats, respectively. In high-probability contexts, the Ce is sufficient to initiate defensive responding, but in low-probability contexts, where the Ce is relatively quiescent, the BST takes the reins. This model is derived from the current findings, as well as decades of experiments in humans, NHPs, and rodents, and has explanatory power that extends beyond the current paradigm. For example, although freezing to a 100% probability CS+ in a tone-shock conditioning paradigm is dependent on Ce^39^, freezing to the CS-can be BST-dependent^40^. Because the CS-is presented a limited number of times in the same context where shocks and cues are paired, our model predicts that it should be treated as a low-probability threat. Because our computational model is focused on the statistics of the environment, it is agnostic to the nature of the threat and can be deployed across a range of assays and species. Thus, this model provides a concrete framework for the kinds of coordinated cross-species experiments that will be necessary to fully elucidate the neurobiological bases of threat processing, from molecules to microcircuits.

### Grounding in a falsifiable computational framework

This approach illustrates how computational models can provide new insights into the nature and neurobiology of emotion. This has both specific and general implications for future research. A specific next step is to determine whether our findings are unique to threat probability or whether they are better explained by expected threat value (probability * intensity), or threat-response policies (Action | probability*intensity). From an adaptive-capacity standpoint, both alternatives are plausible. For example, an expected value account is plausible because a hungry animal should not forgo foraging in the face of a low-intensity threat, even if it is high-probability. In fact, although not formally tested, there are initial indications that otherwise similar threat-anticipation tasks can result in Ce-and-BST activation or BST-only activation depending on the intensity of the threat (e.g.^14,23^). More generally, the current approach provides a sturdy computational backbone for deepening our understanding of fear and anxiety. By anchoring the model on threat probability, it enables cross-fertilization with fields that have productively studied learning and anticipation in the context of sensation, action, language, and reward. For example, a productive line of work would be to incorporate temporal-difference anticipation functions, prediction errors, hazard rates, and more – all parameters that could shape how threat is anticipated. None of this is possible with the canonical model or other narrative accounts of emotion.

### Implications for cross-species basic science

Our findings point to the need for cross-species reverse translation, a prerequisite for identifying the specific mechanisms underlying our results, which include both similarities and differences in Ce and BST function. A distributional or rate code for threat-probability is plausible. From this perspective, individual Ce and BST neurons selectively fire in response to threats that cross a probability threshold, with a greater proportion of BST neurons tuned to low-probability threat. Testing this hypothesis is critical given the complex interrelationships between the BOLD signal, neuronal computations, and cellular physiology^41–44^. Ultimately, these insights will dovetail with ongoing research elucidating the contributions of specific cell-types and molecules that underlie threat-responding.

### Implications for clinical research

Anxiety is a universal component of human experience, a fundamental dimension of temperament and personality, and a core element of multiple psychiatric disorders. Anxious individuals are prone to pervasive distress and arousal, manifesting as exaggerated and contextually inappropriate responses in situations many would consider "safe." Historically, this is often framed as aberrant responses to "uncertain" threats^5^. Our model and NHP data motivate the hypothesis that this actually reflects heightened responses to low-probability threats. For example, when an anxious person struggles to engage with strangers at a party, this could reflect an exaggerated response to the unlikely possibility of scorn or social rejection. From this perspective, it is easy to see how most real-world anxiety is experienced in anticipation of objectively low-probability aversive events. This conceptual insight has the potential to guide research in clinical populations. We anticipate that alterations in the tuning of EA neurons to threat-probability promote the miscalibrated emotional responses that are the hallmark of anxiety disorders. Our prior work demonstrated that dispositionally threat-sensitive monkeys show increased BST metabolism in the ALN-following-NEC context^34^. Alongside studies in clinical populations^28,45–50^, this reinforces the hypothesis that exaggerated BST reactivity to low-probability-threats is a key contributor to extreme trait anxiety and anxiety disorders.

## Conclusion

Anxiety is a fundamental aspect of the human condition. Here, we provide two new insights: first, anticipatory anxiety is primarily driven by threat probability; second, likely and unlikely threats are differentially encoded across the extended amygdala, such that defensive responses to unlikely threats require BST. This provides a computational affective neuroscience framework for understanding emotion. Our new model overcomes nebulous definitions, reconciles decades of research, provides novel hypotheses, and highlights useful avenues for future experiments. Our model makes new predictions about how and why BST-focused interventions could be used to alleviate extreme anxiety and anxiety-related psychopathology^51^. This is important because existing treatments have limited effectiveness, durability, and tolerability. Our approach and model can guide the next generation of systems neuroscience and ultimately inform the development of new and better treatments.

## Materials & Methods

### Human Subject Data Collection and Preprocessing

#### Study Overview and Recruitment

As part of an on-going study focused on understanding the neurocomputational architecture of fear- and anxiety-related states, traits, and symptoms (R01-MH131264), we used the PROMIS Anxiety 8a to screen 2,238 adults residing in the Baltimore-Washington DC metropolitan region (61.5% female; 52.7% White, 12.4% Asian, 16.2% African American, 6.5% Hispanic, 12.2% Multiracial/Other; *M*=28.6 years, *SD*=7.1 years. *Range*=18-45 years)^52–54^. Using established norms, screening data were organized into four bins (bottom quartile, middle quartiles, 75-87.4 percentile, and 87.5-100 percentile), separately for males and females. Individuals who met preliminary inclusion criteria were independently and randomly recruited via email from each of the resulting eight sampling strata. Because of the parent project’s focus on individual differences, ∼10% of subjects were recruited from the bottom quartile, with the remainder evenly split between the remaining three strata. This enabled us to sample a broad range of fear and anxiety symptoms without gaps or discontinuities, consistent with recent recommendations^55,56^. Simulations show that this over-sampling (‘enrichment’) approach does not bias statistical tests to a degree that would compromise their validity^57^. All subjects were right-handed, English proficient, and owned a smartphone; had normal hearing and normal or corrected-to-normal color vision; and reported the absence of lifetime neurological problems, pervasive developmental disorders, MRI contraindications, and prior experience with noxious electrical stimulation. All subjects were free from a lifetime history of psychotic, bipolar, and obsessive-compulsive disorders; severe alcohol/substance misuse; active suicidality (past 3 months); and on-going psychotropic treatment (past 3 months) as determined by an experienced masters-level diagnostician using the Mini International Neuropsychiatric Interview for DSM-5^58^. Likewise, individuals reporting weekly or more frequent recreational use of stimulants, opioids, inhalants, sedatives, hypnotics, or anxiolytics were excluded. Subjects provided informed written consent and procedures were approved by the Institutional Review Board at the University of Maryland, College Park (Protocols #2052626 and #1917962). The study was not preregistered.

#### Participants

A total of 108 individuals were enrolled and scanned. Of these, 4 withdrew from the study due to excess distress or discomfort during the imaging session. This yielded a racially diverse sample of 104 subjects (55.3% female; 40.8% White, 23.3% Asian, 19.4% African American, 9.7% Hispanic, 6.8% Multiracial/Other; *M*=26.7 years, *SD*=7.7 years, *Range*=18-45 years). A total of 2-3 subjects were censored from fMRI or SCR analyses due to insufficient usable data, as detailed below.

#### M-DART fMRI task

The Maryland-Davis Ambiguous and Risky Threat (M-DART) is an fMRI-optimized, randomized, event-related, repeated-measures threat-anticipation paradigm (see **Fig. 1**). Trial order was counterbalanced across subjects and scans (136 trials; 4 scans; 4 orders). Subjects were completely informed about the task design and contingencies prior to scanning. Simulations were used to optimize the detection and deconvolution of task-related hemodynamic signals (variance inflation factors < 2)^59^. Stimulus presentation and ratings acquisition were controlled using Presentation software (version 19.0, Neurobehavioral Systems, Berkeley, CA).

Trials began with the presentation of a thermometer-like visual cue (6-8 s) signaling the likelihood of aversive stimulation and terminated with a compound aversive or benign outcome (1.8 s), including a naturalistic photograph, audio clip, and electrical stimulation. Cues indicated threat likelihood (0-100% in 10% increments), allowing us to model parametric variation in threat Probability (i.e. *likelihood* of aversive outcome) and Risk (i.e. *variance* in aversive outcome). On 36.8% of trials, the cue was partially obscured by a visual mask (10-90% in 10% increments), rendering threat probability partially unknowable and enabling us to model variation in threat Ambiguity (i.e. *range* of potential probabilities). On 76.5% of trials (pseudo-randomized), subjects were prompted to rate the intensity of distress experienced during the prior cue using a 0 (“none”) to 6 (“maximum”) scale and an MRI-compatible response pad (MRA, Washington, PA). All subjects showed adequate engagement with the distress ratings task (>93%). Skin conductance and fMRI were continuously collected throughout.

#### Procedures

Before scanning, subjects practiced an abbreviated version of the paradigm without electrical stimulation until they indicated and staff confirmed via verbal and written assessments that they understood the task. Benign and aversive stimulation levels were individually titrated. *Benign:* Subjects were asked whether they could “reliably detect” a 20 V stimulus and whether it was “at all unpleasant.” If the subject could not detect the stimulus, the voltage was increased by 4 V and the process repeated. If the subject indicated that the stimulus was unpleasant, the voltage was reduced by 4 V and the process repeated. The final level chosen served as the benign electrical stimulation during the imaging assessment (*M*=21.1, *SD*= 4.8). *Aversive:* Subjects received a 100 V stimulus and were asked whether it was “as unpleasant as you are willing to tolerate.” If the subject indicated that they were willing to tolerate more intense stimulation, the voltage was increased by 10 V and the process repeated. If the subject indicated that the stimulus was too intense, the voltage was reduced by 5 V and the process repeated. The final level chosen served as the aversive electrical stimulation during the imaging assessment (*M*=101.3, *SD*=20.0). Following each M-DART scan, we reassessed whether stimulation was sufficiently intense and recalibrated as necessary. In total, 49.5% of subjects adjusted the level of stimulation at least once during the imaging assessment.

#### Electrical stimuli

Electrical stimuli (100 ms; 2 ms pulses every 10 ms) were generated using an MRI-compatible constant-voltage stimulator system (STMEPM-MRI; Biopac Systems). Stimuli were delivered using MRI-compatible, disposable carbon electrodes (Biopac) attached to the fourth and fifth digits of the left hand.

#### Visual stimuli

Visual stimuli (1.8 s) were digitally back-projected (Powerlite Pro G5550, Epson America) onto a semi-opaque screen mounted at the head-end of the scanner bore and viewed using a mirror mounted on the head-coil. A total of 136 photographs (half aversive) were adapted from existing scientific databases (e.g. the International Affective Picture System^60^) and open-access online sources (see *Resource Sharing*).

#### Auditory stimuli

Auditory stimuli (0.80 s) were delivered using an amplifier (PA-1 Whirlwind) with inline noise-reducing filters and ear buds (S14; Sensimetrics) fitted with noise-reducing ear plugs (Hearing Components). A total of 136 auditory stimuli (half aversive) were adapted from open-access online sources (see *Resource Sharing*).

#### MRI Data Acquisition

Data was acquired using a Siemens Prisma 3T scanner (32-channel head-coil). During scanning, foam inserts were used to immobilize the subject’s head. To further mitigate motion artifacts, a strip of medical tape was positioned just above the forehead, providing tactile feedback^61^. Subjects were continuously monitored using an MRI-compatible eye-tracker (Eyelink 1000; SR Research, Ottawa, Ontario, Canada) and FIRMM^62^, enabling staff to provide feedback between scans. Eye-tracking data were not recorded. Short animated films were presented during structural scans to minimize artifacts and fatigue. Sagittal T1-weighted anatomical images were acquired using a magnetization prepared rapid acquisition gradient echo sequence (TR=2,400 ms; TE=2.07 ms; inversion time=1,060 ms; flip=8°; GRAPPA=2; slice thickness=0.8 mm; in-plane=0.8 × 0.8 mm; matrix=300 × 320; field-of-view=240 × 256). A multi-band sequence was used to collect oblique-axial echoplanar images (multiband acceleration=4; TR=1,250 ms; TE=34 ms; flip=75°; slice thickness=2.2 mm, number of slices=64; in-plane resolution=2.2 × 2.2 mm; matrix=96 × 96). Images were collected in the oblique-axial plane (approximately −20° relative to the AC-PC plane) to minimize potential susceptibility artifacts. No in-plane or partial-Fourier acceleration was used. This sequence is similar to that deployed in the HBCD study and affords high spatiotemporal resolution while sidestepping the limitations of aggressive acceleration^63^. For the M-DART task, four 468-volume EPI scans were acquired. The scanner automatically discarded 7 volumes prior to the first recorded volume. To enable fieldmap correction, two oblique-axial spin echo (SE) images were collected in opposing phase-encoding directions (rostral-to-caudal and caudal-to-rostral) at the same location and resolution as the functional volumes (i.e., co-planar; TR=7,800 ms; TE=73.4 ms). Following the last scan, subjects were removed from the scanner.

#### MRI data processing pipeline

Methods were optimized to minimize spatial normalization errors and other potential sources of noise. The general pipeline was adapted from that of Cornwell and colleagues^64^. Data was visually inspected before and after processing for quality assurance.

##### Anatomical Data Processing

T1- and T2-weighted images were inhomogeneity corrected using *N4*^65^ and denoised using *ANTS*^66^. The brain was then extracted using *BEaST*^67^ and brain-extracted and normalized reference brains from *IXI*^68^. Brain-extracted T1 images were normalized to a variant of the 1-mm T1-weighted MNI152 template modified to remove extracerebral tissue (non-linear 6th-generation symmetric average)^69^. Normalization was performed using a four-stage transformation model, including translation, rigid, affine, and the symmetric diffeomorphic normalization in ANTS (version 2.3.4; gradient step=.1 with 32 histogram bins for each linear transformation; for diffeomorphic normalization SyN was used with gradient step=0.15, 64 histogram bins, update-field variance 3.0, total-field variance 0, with Mattes mutual information [64 bins, regular sampling at 60% of voxels] over 500×500×500×500×500 iterations with a convergence threshold of 1e-8 over a window of 40, shrink factors of 8×6×4×2×1, and smoothing sigmas of 4×2×1×0.5×0 voxels)^66^. T2-weighted images were rigidly co-registered with the corresponding T1 prior to normalization. The brain extraction mask from the T1 was then applied. Tissue priors were unwarped to native space using the inverse diffeomorphic transformation^70^. Brain-extracted T1 and T2 images were segmented using native-space priors generated in *FAST* (version 6.0.4) for subsequent use in T1-EPI co-registration^71^.

##### Fieldmap Data Processing

SE images and *topup* were used to create fieldmaps. Fieldmaps were converted to radians, median-filtered, and smoothed (2-mm). The average of the distortion-corrected SE images was inhomogeneity corrected using *N4* and masked to remove extracerebral voxels using *3dSkullStrip* (*AFNI* version 23.1.10). The resulting mask was minimally eroded to further exclude extracerebral voxels.

##### Functional Data Processing

EPI files were de-spiked (*3dDespike*), slice-time corrected to the TR-center using *3dTshift*, and motion-corrected to the first volume and inhomogeneity corrected using *ANTS* (12-parameter affine). Transformations were saved in ITK-compatible format for subsequent processing^72^. The first volume was extracted for EPI-T1 co-registration. The reference EPI volume was simultaneously co-registered with the corresponding T1-weighted image in native space and corrected for geometric distortions using boundary-based registration^71^. This step incorporated the previously created fieldmap, undistorted SE, T1, white matter (WM) image, and masks. The spatial transformations necessary to transform each EPI volume from native space to the reference EPI, from the reference EPI to the T1, and from the T1 to the template were concatenated and applied to the processed EPI data in a single step to minimize incidental spatial blurring. Normalized EPI data were resampled (2 mm^3^) using 5^th^-order b-splines. Analyses employed data that were spatially smoothed (4-mm) using *3DblurInMask*. To minimize signal mixing, smoothing was confined to the gray-matter compartment, defined using a variant of the Harvard-Oxford cortical and subcortical atlases that was expanded to include the bed nucleus of the stria terminalis (BST) and periaqueductal gray (PAG)^73–77^. Extended amygdala (EA) hypothesis testing focused on anatomically defined ROIs (see below). To maximize anatomical resolution, no additional spatial filters were applied, consistent with prior work by our team and recent recommendations^78,79^.

##### EA ROIs

Ce and BST activation was quantified using well-established, anatomically defined, bilateral ROIs and spatially unsmoothed fMRI data^78,79^ (see **Fig. 2** in the main report). The derivation of the Ce ROI is detailed in Tillman et al.^78^. The probabilistic BST ROI was developed by Theiss and colleagues and thresholded at 25%^77^. It mostly encompasses the supracommissural BST, given the difficulty of reliably discriminating the borders of regions below the anterior commissure in T1-weighted images^80^. Bilateral ROIs were decimated to the 2-mm resolution of the fMRI data. EA ROI analyses used standardized regression coefficients extracted and averaged for each parametric modulator (e.g., Probability), region, and subject. Unlike conventional whole-brain voxelwise analyses—which screen thousands of voxels for statistical significance and yield optimistically biased associations—anatomically defined ROIs “fix” the measurements of interest *a priori*, providing statistically unbiased effect-size estimates^81^.

#### Skin Conductance Data Processing Pipeline

Skin conductance data were processed using in-house Matlab (version 9.9.0.1467703) code adapted from *PsPM*^82,83^. Scanner slice-acquisition artifacts were corrected using a 13-Hz notch-filter. Outliers and segments with artifactually low variance were automatically flagged and corrected using piecewise cubic Hermite polynomial interpolation. Scans with excessive artifacts (>30%) were censored. Artifact-corrected data were low-pass filtered (25^th^-order zero-phase IIR filter; cutoff frequency: 0.5-Hz) and down-sampled (12.5 Hz). To ensure data validity, scans that did not show numerically positive skin conductance responses to aversive outcomes (e.g., shock) presentation (averaged across trials) were censored.

#### Data Exclusions for fMRI

Three subjects had insufficient usable fMRI data (<4 M-DART scans; e.g. due to scanner errors) and were excluded from analyses, yielding a final fMRI sample of 101 individuals.

#### Data Exclusions for SCR

Two subjects had insufficient usable SCR data and were excluded from analyses, yielding a final SCR sample of 102 individuals.

### Human Subjects Statistical Analyses

#### Modeling Probability, Risk, and Ambiguity in the M-DART fMRI Task

This section defines how *Probability*, *Risk*, and *Ambiguity* were modeled in the M-DART task to investigate the effects of each threat facet on self-reported distress, SCR, and fMRI BOLD. Details on the experimental design can be found in the *M-DART fMRI task* section, and methods that are specific to the outcome modality can be found below. All modeling was performed separately for unambiguous and ambiguous trials. On unambiguous trials, *Probability* was modeled as the probability of receiving an aversive outcome (i.e. how full the bar was; p_Threat_), and *Risk* was modeled as the variance in outcome (i.e. the inverse quadratic function of threat-probability; Var[p_Threat_]). In this formulation, *Risk* is maximal at when *Probability* is 50%, and lowest when *Probability* of threat is at 0% or 100% (i.e. certain-safety and certain-threat), making *Risk* orthogonal to *Probability* (see **Fig. 1c**). On ambiguous trials, *Ambiguity* was modeled as the amount of the bar that was occluded. The effects of Ambiguity were estimated alongside "rational probability" of threat, to ensure *Ambiguity* results could not be better attributed to *Probability* (see **Fig. 1d**). Rational probability of threat was a measure of a rational agent’s expectation, calculated by taking the known probability and adding half of the ambiguous range, effectively modeling the midpoint of the possible threat probabilities (i.e. known probability + Ambiguity/2; p_Threat_ + .5*A_Threat_; or, the size of the red bar, plus half of the white bar in **Fig. 1a**). This approach accounts for the effect of *Ambiguity* above and beyond the effect of probability^84^. Together, these models allowed us to independently estimate the effects of *Probability*, *Risk*, and *Ambiguity*.

#### Self-reported Distress Analyses

*Effects of Probability, Risk, and Ambiguity on distress ratings.* To test whether distress ratings changed with increasing levels of Probability, Risk, and Ambiguity, we fit a linear mixed-effects model (`mixedlm()` in `statsmodels`; version 0.14.4)^85^ to trial-level ratings. The model jointly estimated the effects of Probability and Risk on unambiguous trials and Ambiguity on ambiguous trials. We also included Rational Probability (i.e. visible threat Probability + Ambiguity/2) as a covariate on ambiguous trials to isolate the effect of Ambiguity and ensure it could not be attributed to changes in threat-probability; we do not interpret or report the Rational Probability effect. We rescaled each predictor within its relevant trials by min–max normalizing to a 0–1 range using the full-sample observed range, mean-centering, and doubling, so that every predictor spanned a common range of 2.

These rescaled predictors were then entered into a single mixed effects model. The model included Probability, Risk, Ambiguity, Rational Probability, and trial type as fixed effects, with unambiguous trials as the reference category, and a random intercept for each subject. We report z-statistics for the effect of Probability, Risk, and Ambiguity in the main text; estimates, standard errors, 95% confidence intervals and exact P values for all reported tests are provided in **Supplementary Table 1**. For visualization, we calculated each subject’s mean distress rating at each regressor level and then averaged across subjects to obtain grand means and standard errors of the mean for Probability (**Fig. 1e**), Risk (**Fig. 1f**), and Ambiguity (**Fig. 1g**).

*Within-subject sensitivity to Probability, Risk, and Ambiguity.* To directly compare the relative effect of Probability, Risk, and Ambiguity on self-report within subjects, we derived subject-specific regression coefficients by fitting the same model described above (Probability, Risk, Ambiguity, Rational Probability, and trial type) separately for each subject, rather than as a single model across the whole sample. As above, predictors were rescaled by min–max normalizing, mean-centering, and doubling, but here using each subject’s own observed range rather than the full-sample range, so that beta estimates reflect each subject’s individual scale. Distress ratings were analyzed in their native units (0–6). From each subject’s model we retained the coefficient for Probability, Risk, and Ambiguity. Rational Probability was again included only as a covariate and not interpreted. To test whether the effect of Probability differed from that of Risk and Ambiguity, we entered these subject-level beta estimates into a linear mixed-effects model with predictor identity as a categorical fixed effect and a random intercept for subject, setting Probability as the reference level. This yielded two planned comparisons — Risk vs. Probability and Ambiguity vs. Probability. We report z-statistics for these comparisons (full statistics in **Supplementary Table 1**). For visualization, we also averaged subject-level beta estimates for each predictor across subjects to obtain grand means and standard errors of the mean for the effect of each predictor on distress ratings (**Fig. 1h**).

### SCR Analyses

#### SCR First-Level Trial Modeling

For each subject, first-level modeling was performed using standard general linear models (GLMs). Each cue was separately modeled using variable-duration rectangular (‘boxcar’) functions convolved with a canonical electrodermal response function (ERF). Epochs corresponding to the presentation of aversive and benign reinforcers and rating prompts were simultaneously modeled using a similar approach. To maximize signal separation, reinforcer presentation was modeled using a reinforcer-specific ERF derived from an independent sample^83^ and its temporal derivative. To mitigate potential drift artifacts, first-level models included 0-4^th^ order polynomials. This produced trial-specific estimates of SCR for each cue.

#### SCR Second- and Third-Level Group Modeling

Because this data provides a trial-level estimate, similar to the distress ratings above, we used the same analytic approach to compute *within-subject sensitivity to Probability, Risk, and Ambiguity*. Trial-level SCR estimates were within-subject z-transformed prior to analysis to account for between-subject variability in electrodermal reactivity. In brief, within-subject models were used to estimate the effects of Probability, Risk, and Ambiguity. Then, the subject-specific parameter estimates were tested in a linear mixed-effects model, to obtain direct comparisons of Risk vs. Probability and Ambiguity vs. Probability. We report z-statistics for these comparisons (full statistics in **Supplementary Table 1**). For visualization, we averaged subject-level beta estimates for each SCR-predictor across subjects to obtain grand-mean and standard errors of the mean (**Fig. 1i**).

#### fMRI Within-Subject Analyses - First-level Modeling

We estimated first-level general linear models (GLMs) using a custom Python pipeline built on the nltools library version 0.5.1^86^. To construct the design matrices, events were initially defined at a high temporal resolution of 1000 Hz, downsampled to the fMRI repetition time (TR), and then convolved with a canonical hemodynamic response function (HRF). A temporal derivative was added to the main effect of the outcome regressor to better model the rapid and complex neural response during the outcome period (which included an aversive stimulus, visual, and auditory cues), accounting for potential timing variations in the BOLD response.

To mitigate low-frequency noise and scanner drift, a combination of polynomial basis functions (up to the third order) and a discrete cosine transform (DCT) basis set (128-s cutoff) were included as nuisance regressors in the design matrices. This ensures that the modeled effects are not confounded by slow, systematic signal changes over the course of each fMRI scan. To account for physiological and motion-related noise, we included several additional nuisance regressors. As in our previous work^64,87^, these included volume-to-volume displacement, six motion parameters and their first differences, six motion-squared parameters and their first differences, brain edge, cerebrospinal fluid (CSF) signal, CSF edge, white matter, and high-frequency noise. Volumes with excessive volume-to-volume displacement (0.5075 mm) and those during and immediately following reinforcer delivery were censored.

Multiple GLMs were used to investigate different aspects of cue processing. Across all GLMs, the outcome and rating periods were modeled identically. We included HRF-convolved rectangular (‘boxcar’) regressors for the outcome-period constant, a mean-centered parametric regressor for the experienced outcome coded as +1 for shock and −1 for no shock, and the rating period spanning the duration of each subject’s actual response time.

The following sections describe the three distinct GLMs, which differed in how the cue period was modeled to address our research questions.

#### Parametric Modulation of Probability, Risk, and Ambiguity

To investigate neural responses to Probability, Risk, and Ambiguity, we constructed a GLM with parametric regressors for the cue period. The cue period, defined as the interval from cue onset to outcome onset, was modeled separately for unambiguous and ambiguous trials.

The cue period for unambiguous trials was modeled using three regressors. The first, a rectangular (‘boxcar’) regressor, captured the main effect of the cue. To account for the linear relationship between neural activity and trial parameters, we included two parametric regressors: one for Probability of threat, and another for the magnitude of Risk. As above, Risk was defined as threat variance, is an inversely quadratic function of probability, with minimal values at a probability of 0 and 1, and a maximal value at a probability of 0.5^88^.

The cue period of ambiguous trials was modeled with a rectangular (‘boxcar’) regressor for the main effect of the cue. This was accompanied by two parametric regressors: one for the Rational Probability of threat under ambiguity, and another for the Ambiguity level. The Rational Probability of threat was a measure of a rational agent’s expectation. It was calculated by taking the known probability and adding half of the ambiguous range, effectively modeling the midpoint of the possible threat probabilities. Prior to entering the GLM, all four parametric regressors (Probability, Risk, Rational Probability, and Ambiguity) were rescaled within each subject by min–max normalizing to a 0–1 range, mean-centering, and doubling, so that each regressor spanned a common range of 2.

#### Discrete-Level Analysis of Probability and Ambiguity Cues

To estimate and visualize the effects of each discrete level of Probability/Risk/Ambiguity, and ensure the pattern was consistent with a linear function, we performed a second GLM. In contrast to the first GLM, which used continuous parametric regressors, this model was designed for a more granular analysis by treating each unique Probability and Ambiguity level as a separate event type. The purpose of this approach was to determine whether neural activity was driven by specific discrete levels of Probability, Risk and Ambiguity, rather than a continuous linear function of these variables.

On unambiguous trials, the cue period was modeled using a set of regressors that captured both the main effect of cue and discrete probability levels (we did not separately model Risk, which can be inferred from Probability). First, a rectangular (‘boxcar’) regressor was used to capture the cue effect. Then, separate boxcar regressors were used to capture each level of threat probability (e.g., 1 if *p_Threat_* = 0.20, 0 otherwise).

Ambiguous trials were modeled similarly, with rectangular (‘boxcar’) regressors for the main effect of cue and for each level of Ambiguity and Rational Probability, defined as visible threat probability + Ambiguity/2 (e.g., 1 if Ambiguity = 50, 0 otherwise).

#### Direct Comparison of Cues as a Function of Probabilities

To investigate specific differences between specific cues, we implemented additional first-level GLMs that compared cues with specific values (e.g. 0%-vs-100%, 0%-vs-50%, or 40-60% vs. 10-30%, etc.). This allowed hypothesis-driven comparisons between predefined groups of trials (e.g. certain-vs-uncertain threat, or changes at differing levels of threat probability). The specific comparisons included: Certain Threat vs. Certain Safe: 0%-vs-100%; Uncertain Threat vs. Certain Threat: 10%-90%-vs-100%; 50% vs. Certain Safe: 50%-vs-0%; Changes as the probability of threat increases: 10-30%-vs-0%, 40-60%-vs-10-30%, 70-90%-vs-40-60%, 100%-vs-70-90%, 10%-vs-0%, and Ambiguous-vs-Unambiguous trials.

In each model, the cue period was modeled with three regressors: a rectangular (‘boxcar’) regressor to capture the main effect of the trials included in the comparison, a parametric regressor that directly contrasted the trials of interest coded as +1 and -1, (e.g. 100% as coded as +1, and 0% coded as -1 for Certain Threat vs. Certain Safe), and a third boxcar regressor that accounted for all trials not part of the comparison.

#### fMRI Group-level Analyses - Second-level Modeling

We estimated group-level effects for each contrast voxelwise using a one-sample t-test against zero, implemented as ordinary least squares regression with an intercept-only design matrix via the `regress()` method of the `Brain_Data` class in `nltools`(version 0.5.1)^86^. This produced whole-brain t- and p-value maps for each contrast.

We corrected voxelwise p-values for multiple comparisons using the Benjamini–Hochberg false discovery rate (FDR) procedure^89^, implemented with the `fdrcorrection()` function from `statsmodels`^85^, at q < 0.005. For the Probability, Risk, and Ambiguity parametric effects, we applied a single FDR correction across their pooled p-values, treating them as one family because they were estimated within the same first-level model and reported together. We corrected all other contrasts independently.

### fMRI Brain-signature Analyses

As a convergent test, we examined whether responses to Probability, Risk, and Ambiguity differentially reflected two independently validated, previously published whole-brain neural signatures: a generalized negative-affect signature and an anticipatory-anxiety signature. The generalized negative-affect signature^26^ was derived as the common component of negative affect shared across four aversive modalities: mechanical pain, thermal pain, aversive sounds, and aversive images. The anticipatory-anxiety signature^27^ was associated with anxious anticipation specific to threat uncertainty. Each signature is defined by a whole-brain weight map that assigns every voxel a coefficient, derived from a multivariate predictive model trained on independent data. The sign and magnitude of each coefficient indicate how strongly, and in which direction, that voxel’s activity contributes to the psychological construct the signature was built to capture.

We computed signature prediction scores for six first-level contrast maps, organized into two sets of three: a parametric set (Probability, Risk, Ambiguity) and a categorical set (Certain Threat vs. Certain Safe, Uncertain Threat vs. Certain Threat, Ambiguous vs. Unambiguous). For each subject, contrast, and signature, the pattern-prediction score was calculated as the dot product between the subject-level first-level beta map and the corresponding signature weight map. Higher scores indicate a closer match between the contrast’s activity pattern and the signature, reflecting stronger expression of the corresponding psychological construct (e.g., negative affect or anxious anticipation); lower scores indicate a weaker match. The generalized negative affect signature results for the parametric set are presented in the main text (**Fig. 1k**); the remaining three — the anticipatory-anxiety signature for the parametric set, and both signatures for the categorical set — are presented in **Supplementary Fig. 1** (panels **f**, **d**, and **e**, respectively).

For each signature and contrast set, we tested whether pattern-prediction scores differed across contrasts using a linear mixed-effects model with contrast as a categorical fixed effect and a random intercept for subject. This produced four models in total, one per signature × contrast-set combination, while accounting for the repeated-measures structure of scores within the subject. These models yielded z-statistics for each predictor relative to its reference level, set to Probability for the parametric set and Certain Threat vs. Certain Safe for the categorical set. Full statistics are provided in **Supplementary Table 1**.

### fMRI ROI Analyses

We performed ROI analyses using ROIs for BST and Ce (defined above). ROIs were used to mask spatially unsmoothed first-level statistical maps. We avoided spatially smoothed maps because smoothing can blur signals from neighboring structures into these small subcortical regions, biasing the resulting ROI estimates^50^.

We applied this masking procedure to first-level maps from first-level GLMs (defined above), to characterize how Ce and BST respond to each regressor of interest (e.g. Probability, Risk, Ambiguity, etc.). For each subject, map, and ROI, we computed the mean parameter estimate by averaging voxelwise beta values within the binarized ROI mask, yielding a single scalar per subject × map × region. We tested for regional and regressor-related differences in mean beta values using a linear mixed-effects model with regressor, region, and their interaction as fixed effects, and a random intercept for subject (**Fig. 2a**, **Fig. 3b**). Where relevant, we ran post-hoc comparisons as linear combinations of the model’s fixed-effect coefficients, applying the Benjamini-Hochberg procedure to control the false discovery rate (FDR; q=0.05) when more than one contrast was tested within a given model. We report z-statistics for each comparison of interest (full statistics in **Supplementary Table 1**).

For probability-bin transitions (e.g., 40–60%-vs-10–30%), we computed a per-subject BST−Ce difference score for each transition. We tested for an overall effect of transition using a one-way ANOVA and reported the F-statistic for this omnibus test. We then evaluated planned, Holm-corrected pairwise comparisons, derived as linear combinations of the fixed-effect coefficients from a linear mixed-effects model with a random intercept for subject, comparing the lowest-probability transition (0% vs. 10–30%) against each of the other three transitions (**Fig. 3k**). For these comparisons, we report z-statistics (full statistics, including the omnibus F-test, in **Supplementary Table 1**).

### fMRI Timecourse Regression Analysis

To characterize the temporal evolution of threat Probability-, Risk-, and Ambiguity-related signal following cue onset, and to ensure that these effects were not being driven by the outcome, we used a single-trial regression approach. This approach was applied to two measures of brain activity: (1) whole-brain pattern-expression of the negative affect signature^26^ (**Fig. 1l**), and (2) mean activity within anatomical regions of interest (Ce, BST; **Supplementary Fig. 5**). For both measures, we linearly detrended and z-transformed each voxel’s timecourse within scan before spatially collapsing it into a single signal value per TR. For the whole-brain analysis, this value was computed as a dot product of voxel signals against the signature’s weights. For the two ROI analyses (Ce and BST), we computed it as a simple average across voxels. For both kinds of analyses, this was done for each trial (i.e. from -1 s before cue onset until 8 s following cue onset). To maximize our ability to discern timecourse differences when the TR was not locked to the stimulus onset, we upsampled this signal to 10 Hz via linear interpolation. This produced cue-aligned windows of estimated BOLD from 1 second before cue-onset to 8s after cue onset for each combination of trial and subject.

At each timepoint within this window, we fit a within-subject (‘first-level’) linear regression model using Probability, Risk, and Ambiguity as predictors of interest. Probability and Risk served as predictors only on unambiguous trials, whereas Rational Probability and Ambiguity served as predictors only on ambiguous trials (see *Modeling Probability, Risk, and Ambiguity for the M-DART fMRI Task* for regressor definitions). We also included Trial Type, coded as +1 for unambiguous and −1 for ambiguous trials, and Outcome, coded as +1 for shock and −1 for no-shock, as additional predictors. Outcome was included as a control variable rather than a regressor of interest because Probability of threat and actual shock occurrence are inherently correlated (higher-probability trials are more likely to end in shock), modeling outcome alongside Probability ensures that cue-period estimates of Probability are not confounded by trial Outcome. This analysis yielded a timecourse of regression coefficients for each predictor per scan (e.g. a beta for the effect of probability for each 100ms bin in each scan). For each signature/ROI, we averaged regression coefficients over scans to produce a single timecourse per predictor for each subject.

To test whether each predictor’s beta timecourse differed significantly from zero, we ran one-sample t-tests on the subject-level beta estimates against zero at each timepoint within the 0–8 s window. Within each signature/ROI, we pooled the resulting p-values across all predictors and timepoints into a single family and corrected for multiple comparisons using the Benjamini-Hochberg false discovery rate (FDR; q=0.05) procedure. For visualization, we plotted the mean and standard error of the mean of the subject-level beta timecourses across subjects at each timepoint, separately for each predictor and each signature/ROI, and marked timepoints surviving FDR correction as horizontal lines beneath each timecourse, with color indicating the corresponding predictor’s line.

### fMRI Modeling of Regional BOLD Responses to Threat Probability

To test whether Ce and BST differ in their sensitivity profile across the threat-probability range, we modeled each region’s BOLD response as a function of threat probability using a sigmoid function, characterized by its inflection point (*x_0_*, the probability at which the response rises) and slope (*k*; **Fig. 3h-j**). Because individual subject fits are often unstable, we fit the logistic function across subjects. To provide a basis for statistical comparison, we used a subject-level bootstrap to generate empirical distributions of these parameters for each region. For Ce and BST, we extracted subject-level mean parameter estimates (β) for each threat-probability cue (0, 10, 20, …, 100%). Within each region, β estimates were normalized to the group-mean response at the probability extremes to yield a normalized response y,

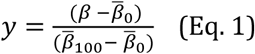

where β̅_0_ and β̅_100_denote the group-mean β at 0% and 100% threat probability, respectively, such that 0% and 100% threat probability corresponded to normalized responses of 0 and 1. Within each region, we fit a logistic sigmoid to the pooled normalized responses across subjects,

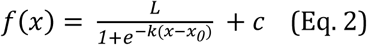

where *L* is the upper asymptote, *x* is threat probability, *x_0_*is the inflection point, *k* is the slope, and *c* is an offset that anchors the fitted curve to the median normalized response at the lowest threat probability (0%). Because normalized responses spanned the interval [0, 1] by construction, *L* was fixed at 1. *x_0_* and *k* were estimated by nonlinear least-squares (`scipy.optimize.curve_fit`). For each region, we performed 1,000 bootstrap iterations. In each bootstrapping iteration we drew beta estimates from a random selection of 50 subjects with replacement and fit the sigmoid function (Eq. 2) to the pooled normalized responses of the resampled subjects; all 1,000 iterations converged in both regions. This produced an empirical distribution of 1,000 (*x_0_*, *k*) pairs per region (**Fig. 3h,i**). Bootstrap distributions of *x_0_* and *k* were compared between Ce and BST using two-sided Mann-Whitney U tests (**Supplementary Table 1**). Results were visualized as the mean curve across bootstrap iterations, with its 99.9% confidence interval (**Fig. 3j**); the distributions of fitted parameters across iterations are shown in **Fig. 3h,i**.

## NHP Methods

All procedures were performed in accordance with federal guidelines and were approved by the University of Wisconsin-Madison Institutional Animal Care and Use Committee. The monkeys were housed at the Wisconsin National Primate Research Center and the Harlow Center for Biological Psychology. Animals were pair-housed in standard primate cages with a twelve-hour light/dark cycle, fed chow twice daily with *ad libitum* water access and daily enrichment. Experimental design and techniques were aimed at minimizing animal use and suffering. All drugs and treatments were given in consultation with veterinary staff.

### NHP Subjects

Fourteen adult female rhesus macaques (*Macaca mulatta*) were selected for this experiment. Seven underwent bilateral ibotenate infusions targeting the bed nucleus of the stria terminalis (BST) using an MRI-guided surgical procedure^90^. Each of the lesioned animals was paired with a cage-mate control. Two of these pairs were dropped over the course of the study and were used in other experiments. This resulted in the following sample sizes for each test: *HIP-NEC-only Paradigm* n=7 pairs; *Novel Conspecific Paradigm*, n=7 lesioned and n=6 control animals; *HIP-Persistence Paradigm* n=6 pairs; and *HIP-Persistence Paradigm* second test n=5 pairs.

### NHP Intraoperative MRI-guided infusion surgery

The placement of trajectory guide bases under sterile conditions followed published methods^91–93^ modified to target the dorsal BST and are only briefly described here. Animals were anesthetized with ketamine (up to 20 mg/kg, IM), intubated and prepared for surgery, and then placed in an MRI compatible-stereotaxic head frame. Surgical subjects received isoflurane anesthesia for the duration of the procedure along with atropine (0.01–0.4 mg/kg, IM) and buprenorphine (0.01–0.03 mg/kg, IM, repeated every 6–12 h). Plasmalyte (up to 10 mg/kg/h i.v.) was administered to maintain fluids. Cefazolin (20–25 mg/kg, IM or i.v.) was administered 1 day before surgery and immediately prior to surgery, and then every 6 hours while under anesthesia. Vital signs (heart rate, respiration, oxygen saturation, and end-tidal CO_2_) were continuously monitored. Body temperature was monitored during the surgical and MRI procedures and was maintained in the MRI by wrapping the animals for warmth while incorporating a hot water heating device within the wrap.

### NHP MRI Acquisition

The surgical subject remained in the head frame and was transported under anesthesia from the surgical suite to the MRI suite. Sterile conditions were maintained throughout transport and during the iMRI procedure. A custom built 3" circular surface coil (MR Instruments, Minneapolis, MN) was positioned above the animal’s head with the trajectory guide base located near the center of the coil. The procedure began by acquiring a structural T1-weighted "roadmap” image for intraoperative targeting of the BST using a 3D Inversion Recovery Gradient Echo (IR GRE) MRI sequence with a 12° flip angle, 512 x 256 x 512 matrix (0.352 x 0.352 x 0.40 mm voxels) in a coronal slab, TR = 9.1 ms, TE = 3.9 ms, inversion time = 450 ms, 97 Hz/pixel receiver bandwidth, with a 6.5 minute total acquisition time.

The surgery targeted dorsal BST region above the anterior commissure and medial to the internal capsule, and were designed to include the lateral dorsal BST (which corresponds to the oval nucleus in the rodent and matches the peak BST activation in our human studies). To lesion as much of the dorsal BST as possible while minimizing damage to surrounding regions, a single 2.0 µl infusion was placed in each hemisphere. An effort was made to maximize lesion accuracy by aligning the catheter guide along a posterolateral to anteroventral trajectory that pointed to the intersection of the internal capsule and the anterior commissure at the most anteroventral aspect of the dorsal caudate. Adjustments were made to the injector depth based on images obtained from the intraoperative MRI scans to bring the tip of the catheter as close to anterior commissure as possible without penetrating the ventricle.

The cannula was loaded with an infusate of 1.0 mg ibotenic acid (Abcam, Waltham, MA; catalog # ab120041) per 100 µl of PBS mixed with 0.66 mM of the MR-visible contrast agent gadolinium (Multi-Hance; Bracco Diagnostics, Monroe Township, NJ, USA), which was delivered at a rate of 0.2 µL/min. For a qualitative visualization of the volumetric infusate delivery region, immediately following each infusion the previously described 3D roadmap was reacquired to capture contrast agent uptake at the injection site. Post-infusion T1-weighted scans were used to create the gadolinium cloud overlap image (see method for *Lesion Estimation*). After the infusions were complete, the animal was transported back to the surgical suite and the trajectory guide bases were removed. To reduce intracranial pressure and prevent brain swelling, mannitol (up to 2.0 g/kg, i.v.) was given as needed. The incision was closed in layers before the animal was allowed to recover from anesthesia. Animals were given buprenorphine twice on the day following the surgery (0.01–0.03 mg/kg, IM). Cefazolin (20–25 mg/kg, IM or i.v.) or cephalexin (20–25 mg/kg, PO) was given twice daily for 5 days after surgery to prevent infection. The animals were allowed to recover for an average of 100.57 days before post-surgery testing commenced.

### NHP Behavioral Testing: Human Intruder Paradigm-Persistence (HIP-Persistence Paradigm)

Animals were placed in a testing cage (L: 76.5 cm × W: 71.5 cm × H: 79 cm) with a Plexiglass front for unobstructed viewing. A human ‘intruder’ unfamiliar to the monkey entered the room and stood 2.5 meters from the testing cage. The intruder stood for 30 minutes with their profile presented to the animal, avoiding direct eye contact and making all efforts to remain motionless. After 30 minutes of the no-eye-contact (NEC) condition the intruder left the room and the animals’ behavior was observed for an additional 30-minutes while alone in the testing cage.

### NHP Behavioral Testing: Human Intruder Paradigm-NEC Only (HIP-NEC-only Paradigm)

Animals were placed in a testing cage (L: 76.5 cm × W: 71.5 cm × H: 79 cm) with a Plexiglass front for unobstructed viewing. A human ‘intruder’ unfamiliar to the monkey entered the room and stood 2.5 meters from the testing cage. The intruder stood for 30 minutes with their profile presented to the animal, avoiding direct eye contact and making all efforts to remain motionless.

### NHP Behavioral Testing: Novel Conspecific Paradigm

Testing in the novel conspecific paradigm occurred on a single day, with no pre-training. An older, larger, more dominant animal, referred to as the ‘stimulus animal,’ was placed in a double-length testing cage (L: 153 cm × W: 71.5 cm × H: 79 cm). The stimulus animal was allowed to habituate to the cage for 5 minutes. A metal dividing panel was then inserted to separate the testing cage into two equal sized compartments, one containing the stimulus animal. The test animal was then placed in the other compartment and allowed to habituate for ∼2 minutes before testing began. The divider was removed and the animals were allowed to freely interact for 30 minutes.

### NHP Behavioral Testing: Behavioral Coding

Behavior was scored by an experienced observer blind to the groups using a closed-circuit audiovisual system. Freezing was defined as a tense body posture for more than 3 seconds in the absence of other movements except slow movements of the head or eye blinks. In addition to dyadic interactive behaviors, nonsocial behaviors, such as freezing and locomotion, were also scored, as well as ‘coo’ and ‘bark’ vocalizations (data not shown).

### NHP MRI Data Processing

Raw DICOM files were converted to NIfTI-1 format using dcm2niix (v1.0.20260416)^94^. For each subject, the pre-procedure BRAVO sequence roadmap scan was used for co-registration to the NMT reference template.

Intensity inhomogeneity correction was applied to the reference T1 using the N4 bias field correction algorithm^65^ as implemented in ANTsPy (v0.6.3), with a shrink factor of 4, B-spline control-point spacing of 200 mm, and [50, 50, 50, 50] iterations at each of four resolution levels (convergence tolerance 1e-07). Brain extraction was performed using NHP-BrainExtraction (muSkullStrip, http://github.com/HumanBrainED/NHP-BrainExtraction), a 2-D slice-wise U-Net trained on multi-site non-human primate MRI and implemented in PyTorch (v2.12.0). The model checkpoint ‘Site-All-T-epoch_36_update_with_Site_6_plus_7-epoch_09’ was used. The predicted binary brain mask was multiplied by the N4-corrected volume to produce the skull-stripped image, which was then cropped to the tight bounding box of non-zero voxels with a 5-voxel isotropic margin. Intensities exceeding the 99th percentile of brain voxels were clipped to that value to suppress scanner saturation artefacts prior to registration.

Each subject’s skull-stripped, cropped, and clipped reference T1 was registered to the NMT v2.0 symmetric full-head skull-stripped template (NMT_v2.0_sym_fh_SS.nii.gz; 400 × 400 × 300 voxels, 0.25 mm isotropic)^95^ using ANTsPy. Registration employed the SyNRA transform sequence (centre-of-mass initialisation, followed by rigid, affine, and symmetric diffeomorphic normalisation stages). The similarity metric was Mattes mutual information (32 histogram bins) at all stages. SyN-specific parameters were: gradient step 0.25, flow smoothing σ = 1.5 mm, total smoothing σ = 0, with (100, 70, 50, 40) iterations across four resolution levels. A binary mask derived from the SARM atlas^96^ (all non-zero regions, dilated by 8 mm) was applied at every registration stage to restrict the objective function to subcortical brain regions, and ensure overlap in the injected region that was robust to variation in skull-stripping that resulted from the craniotomy.

All remaining post-infusion BRAVO scans for each subject were brought into template space in two steps: each scan was first rigidly registered to the subject’s reference T1 in native space (Mattes MI, 32 histogram bins) to correct for inter-scan head motion, and the resulting transform was then composed with the subject-level SyNRA transforms and applied via linear interpolation, yielding every scan resampled to the 0.25 mm isotropic NMT grid.

### NHP Lesion Estimation

Lesion’s extent was estimated based on MRI scans, using Python (v3.12.12) using nibabel (v5.4.2), NumPy (v2.3.5), and SciPy (v1.15.3). In short, the difference between post-infusion scans and pre-infusion scans were used to detect the location of Gd after each injection and used to create a binary mask reflecting the estimated injected region for each subject.

For each subject, differences between the pre- and post-infusion differences in T1 intensity were used to estimate lesion extent. First voxels outside the brain mask were set to zero, and T1 images were first scaled by the 99th percentile of their positive brain voxels and clipped to [0, 1]. This step was necessary to prevent large spurious differences at the skull-strip boundary. Thresholding and seed selection were restricted to a subcortical sub-region of the search mask to avoid spurious cortical and ventricular signals. Specifically, we calculated pre-post difference images in areas near the target BST region (within 8mm of the BST in the SARM atlas^96^, above the anterior commissure; 287,792 voxels, 4497 mm³). Within the target-region mask, the mean (μ) and standard deviation (σ) of the hyperintensity map were computed, and voxels exceeding a z-value of 3.0 were binarised. Connected components were labelled using ‘scipy.ndimage.label’. To isolate the infusion site, the seed anchor was defined as the z-restricted mask region with a dorsal ceiling set 5 mm (20 voxels) above the most superior voxel of the BST atlas label. This ceiling prevents superficial white-matter artefacts dorsal to the BST from overriding the injection peak during seed selection. The connected component containing the peak hyperintensity voxel within this seed region was retained; if that voxel was sub-threshold, the component whose centroid lay nearest to the seed peak was selected. A morphological opening (3 × 3 × 3 structuring element, 1 iteration) was applied to sever thin bridges connecting the injection component to distant structures; for compact injection sites smaller than 50 mm³ the opening was skipped when the eroded core shrank to less than 25% of the original volume, since small spherical blobs have no structural bridges but cannot survive erosion. This approach was complemented with inclusion of hypointensity detection within 2mm of the hyperintensity mask to ensure that lesions estimate was not impacted by susceptibility artifacts at the injection site. Components with volumes outside 5–2000 mm³ were discarded. This resulted in an estimate of the injection location for each infusion in each subject. All estimated lesions were manually inspected on the post-surgery T1 images to ensure accuracy.

Per-infusion maps were then summed across all 7 animals, yielding a voxel-wise integer image with values ranging from 0 (no subjects) to 7 (all subjects). Overlap images were saved in NMT template space and visualised as overlays on the NMT skull-stripped template using a discrete seven-level sequential colormap (**Fig. 4a; Supplementary Fig. 7f**).

### NHP Summary and validation of Lesion Location Estimates

Injections were generally located lateral to the internal capsule above the anterior commissure, in the region of the BST. However, most lesions were not specific to the BST region. In fact, the majority of subjects had damage slightly superior to the BST, corresponding to the ventral anterior caudate region that borders the BST. A subset of subjects had limited overlap with the BST in at least 1 injection (i.e. **Fig. 4a**). Estimates of lesion extent were compared to post-mortem GFAP/NeuN staining on a subset of subjects (e.g. **Fig. 4c**), which supports these MRI-based analyses. Post-mortem validation confirmed the interpretation that these lesions damaged a subset of dorsal BST, included portions of ventral caudate, and in some cases included limited portions of the BST. The estimates of BST overlap are indicated by markers in all figures and shown in **Fig. 4f**.

### NHP Tissue collection and sectioning

A subset of experimental animals (n = 4) were euthanized by transcardial perfusion with heparinized PBS at room temperature, followed by fixation with 4% paraformaldehyde (PFA) in PBS at 4°C. Brains were removed, and fixation was continued in the perfusion solution overnight at 4°C. Brains were placed in a brain block and cut into slabs (approximately 14 mm thick) that were subsequently processed through increasing concentrations of sucrose: 10%, 20%, and 30%. The slab containing the BST was frozen and cut at 40 microns on a cryostat (CryoStar NX50, ThermoFisher Scientific). Tissue sections were stored in cryoprotectant (40 mM potassium phosphate/11mM sodium phosphate buffer, pH 7.2, containing 30% ethylene glycol and 30% sucrose) at -20°C until use.

### NHP Immunohistochemistry and histology

To evaluate the location and extent of the ibotenic acid-induced lesion, chromogenic labeling of neuronal nuclei (NeuN) was performed to observe regions of neuronal loss, while labeling of glial fibrillary acidic protein (GFAP) was used to observe regions of glial cell activation. Sections adjacent to those stained for NeuN and GFAP were processed for acetylcholinesterase (AChE), a cholinergic marker that was used to identify anatomical boundaries in the region of the BST using a previously published procedure^97^.

Prior to staining, tissue sections were washed with PBS at room temperature to remove cryoprotectant. To block endogenous peroxidase activity, tissue sections were incubated in 0.9% hydrogen peroxide diluted in PBS containing 0.02% Triton X-100 for 30 minutes. All subsequent incubations were carried out at room temperature in PBS containing 0.3% Triton X-100. Three 5-min washes with PBS were performed in between each incubation step.

*NeuN Detection.* Sections were blocked in 5% normal horse serum (Vector Laboratories, Burlingame, CA) for 1 hour. This was followed by an overnight incubation with a 1:1,000 dilution of mouse monoclonal anti-NeuN antibody (Millipore Sigma, Burlington, MA; catalog # MAB377), and a 1-hour incubation with a 1:200 dilution of biotinylated horse anti-mouse secondary antibody (Vector Laboratories).

*GFAP Detection.* Sections were blocked in 2% normal goat serum for 1 hour (Vector Laboratories). This was followed by overnight incubation with a 1:10,000 dilution of rabbit polyclonal anti-glial fibrillary acidic protein (GFAP) antibody (Agilent DAKO, Santa Clara, CA; catalog # z033429-2), and a 1-hour incubation with a 1:200 dilution of biotinylated goat anti-rabbit secondary antibody (Vector Laboratories).

*Chromogenic Development and Imaging.* Chromogenic signal was developed using the Vectastain ABC Kit with diaminobenzidine (DAB) peroxidase substrate (Vector Laboratories). Sections were mounted on Superfrost Plus microscope slides (Brain Research Laboratories, Waban, MA), which were air-dried, dehydrated in graded ethanol (95% and then 100%), and cleared in xylene for 5 min before cover slipping with Permount mounting medium (Fisher Scientific, Waltham, MA). Images were captured using a DM6000B light microscope (Leica Microsystems, Buffalo Grove, IL, USA).

### NHP Statistical Analyses

All statistical analyses for NHPs were performed using linear mixed effects models, with paired-animals as the random variable to account for shared experiences between cage-mates. Behavioral data were computed as log-transformed durations in 5-min intervals. Mixed effects models were fitted using behavior as the dependent variable, fixed-effect predictors for Group and Interval and pair as a random effect (`mixedlm()` in `statsmodels`; version 0.14.4)^85^. We tested for a significant effect of the Group (BST-lesion vs. Control) parameter estimate, after controlling for pair and interval. Group differences in ALN − NEC difference scores were tested with linear models including pair as a covariate. Full statistics are provided in **Supplementary Table 1**.

## Preregistration

This study was not pre-registered.

## Resource Sharing

Raw data and select materials are publicly available at the National Institute of Mental Health Data Archive (https://nda.nih.gov/edit_collection.html?id=4400). Neuroimaging maps are available at NeuroVault (https://neurovault.org/collections/UCNAATUK/). Task materials are available on Zenodo (https://zenodo.org/records/22866010). De-identified processed data and analytic code is available on GitHub (https://github.com/foxlab-ucdavis/M-DART_Paper).

## Acknowledgments

Authors acknowledge assistance and critical feedback from V. Elam, E. Fekete, L. Friedman, R. Kovner, B. Nacewicz, M. Olsen, L. Pessoa, M. Riedel, M. Tabor, L. Williams, members of the Affective and Translational Neuroscience laboratory, staff of the Maryland Neuroimaging Center, Office of Academic Computing Services at the University of Maryland, and the University of Wisconsin Harlow Center for Biological Psychology and Lane Neuroimaging Laboratory. This work was partially supported by the California National Primate Center, Wisconsin National Primate Research Center, University of Wisconsin HealthEmotions Research Institute, National Institutes of Health (MH107563, OD011106, AA030042, AA031261, MH131264, MH121409, MH121735, MH128336, MH129851, OD011107, MH126426, MH046729, MH081884), University of California, University of Wisconsin, and University of Maryland. Authors declare no conflicts of interest.

## Author Contributions

A.S.F., A.J.S., K.A.D., and J.F.S. designed the overall study. A.S.F. envisioned the present project. A.S.F., J.F.S., and A.J.S. developed and optimized the human neuroimaging paradigm. K.A.D. managed human data collection and study administration. K.A.D., P.R.D., Z.S.S., D.B.W., J.H., J.V.C, E.L., J.K.E, G.B.C., and J.F.S. collected human data. M.O., J.F.S. and A.S.F. developed data processing and analytic software for human neuroimaging analyses. J.F.S. processed human neuroimaging data. A.S.F., M.O., C.M.D., J.F.S., and A.J.S. developed the analytic strategy and analyzed human data. M.O., C.M.D., A.S.F., E.D.B., K.N.K., J.F.S., and A.J.S. interpreted human data. N.H.K, J.A.O. and A.S.F. designed the nonhuman primate experiments. J.A.O. and N.H.K oversaw the nonhuman primate surgeries and data collection. P.H.R., C.W.C. and A.S.F. performed lesion quantification. J.C. and A.S.F. performed data processing and analysis of nonhuman primate data. M.O. A.S.F. and A.J.S. interpreted nonhuman primate data. M.O, C.M.D., A.S.F., K.N.K., J.F.S., and A.J.S. wrote the paper. M.O. and A.S.F. created figures. M.O. and K.A.D. created tables. A.S.F., E.D.B., N.H.K., and A.J.S. funded and supervised all aspects of the study. All authors contributed to reviewing and revising the paper and approved the final version.

