## Supplementary Material for "Extended amygdala function is governed by threat probability, not uncertainty"

### Supplementary Note, Figures and Tables

#### Supplementary Note

The primary analyses reported in the main text show that threat Probability is the dominant driver of whole-brain fMRI activation, with Risk and Ambiguity showing comparatively weaker effects (**Fig. 1j**). Nevertheless, both dimensions of threat uncertainty produced reliable neural correlates that replicate and extend prior neuroimaging findings. Here, because most prior work has focused on rewards with only a handful of studies utilizing aversive stimulation, we discuss the neural correlates of Risk and Ambiguity in the MDART task in greater detail.

##### Effects of Risk

Neural responses to graded risk were modeled as a parametric modulation of outcome uncertainty, using a reverse-U-shaped function that peaked at the 50% threat-probability cue and approached zero at the 0% and 100% threat-probability cues. Risk-related activity was centered primarily in bilateral insular cortex, with peaks in the anterior insula and adjacent extension into orbitofrontal cortex (OFC). Additional activity was observed in paracingulate cortex, medial frontal cortex and lateral frontal regions, including inferior frontal gyrus (IFG) and middle frontal gyrus (MFG; $q < 0.005$, FDR-corrected; **Supplementary Fig. 4**; **Supplementary Table 5**).

This pattern is consistent with prior work showing that risk processing recruits a distributed network spanning insular, medial frontal, and lateral frontal regions[1,2]. For instance, Preuschoff et al.[3] linked anterior insula activity to anticipated risk and risk-prediction errors. Mohr et al. 's[1] meta-analysis further showed that right anterior insula and dorsomedial prefrontal cortex (dmPFC) were consistently recruited across paradigms involving either potential gains or losses and across tasks involving either decision-making or risk anticipation. In contrast, the left anterior insula was particularly evident when potential losses were involved and during risk anticipation. This pattern is especially relevant to our task, in which cues signaled the probability of an upcoming aversive outcome without requiring a choice. Bilateral anterior insula and medial frontal recruitment may therefore reflect processes involved in calculating or anticipating risk when potential outcomes carry aversive significance. It is reasonable to suspect that this is relevant to the individual's intention to increase certainty about what will happen and dedicate additional mental effort when appropriate.

Paracingulate and lateral frontal activity may reflect broader processes engaged during probabilistic cue processing. Prior studies have linked anterior cingulate and lateral prefrontal activity to outcome uncertainty and anticipatory arousal measured via skin conductance response[4] and have shown that lateral prefrontal activity varies with dynamically changing probabilistic uncertainty[5]. These findings suggest that neural systems are characterized primarily in economic choice and reward/loss paradigms are also engaged during the anticipation of uncertain aversive outcomes and are not specific to the modality or valence of the outcome.

##### Effects of Ambiguity

Neural responses to graded Ambiguity were modeled as a parametric function of cue-mask coverage, with greater masking corresponding to less precise information about threat probability. Ambiguity-related activity was most prominent in frontal regions, including bilateral middle and superior frontal gyri, IFG, frontal pole, and OFC, with additional involvement of insular cortex and inferior parietal cortex ($q < 0.005$, FDR-corrected; **Supplementary Fig. 4**; **Supplementary Table 6**).

Our findings are consistent with prior work implicating frontal and parietal regions in ambiguity processing across both decision-making and passive anticipation paradigms. Huettel et al.[6] reported ambiguity-related activity in posterior IFG and posterior parietal cortex during monetary choices under ambiguity. Bach et al.[7] subsequently showed that similar regions, including posterior IFG extending into MFG and posterior parietal cortex, responded to ambiguity during passive anticipation of painful electric shocks, even in the absence of a decision. A coordinate-based meta-analysis by Wu et al.[2] further confirmed this network, identifying left MFG/IFG and right inferior parietal lobule (IPL) as ambiguity-specific regions in a direct ambiguity versus risk contrast. Our results similarly revealed ambiguity-related activity in IFG, MFG, and inferior parietal cortex, suggesting that this frontal–parietal response is not specific to economic choice or appetitive outcomes, but also supports the processing of incompletely specified aversive probabilities.

The convergence with Bach et al.[7], the only prior study employing a comparable passive Pavlovian design, is particularly notable and strengthens the case that IFG and posterior parietal cortex track ambiguity as a representational property of threat cues regardless of whether a choice is required. The additional involvement of superior frontal gyrus and frontal pole, not consistently reported in prior ambiguity studies, may reflect the greater statistical sensitivity of our larger sample (N = 101) or other yet to be identified nuances of these tasks.

OFC and insular involvement are also consistent with prior ambiguity research. Hsu et al.[8] found that increasing ambiguity was associated with greater OFC activity during choices involving incompletely specified outcome probabilities, whereas Wu et al.'s[2] meta-analysis identified anterior insula as a consistent correlate of ambiguity processing. However, both OFC and anterior insula were also recruited by the Risk in the present study, suggesting that these regions may contribute to uncertainty processing more broadly rather than responding selectively to Ambiguity. Consistent with this interpretation, Wu et al.[2] identified the right anterior insula as the only region showing robust convergence across Risk and Ambiguity, whereas our conjunction analysis revealed shared anterior insula activity in the left hemisphere. Both findings implicate anterior insula as a common neural correlate of Risk and Ambiguity, suggesting that it may support uncertainty processing across both known and incompletely specified threat probabilities – regardless of the nature of the uncertainty, valence of the stimulus, or relevance to decision-making.

#### Supplementary Figures

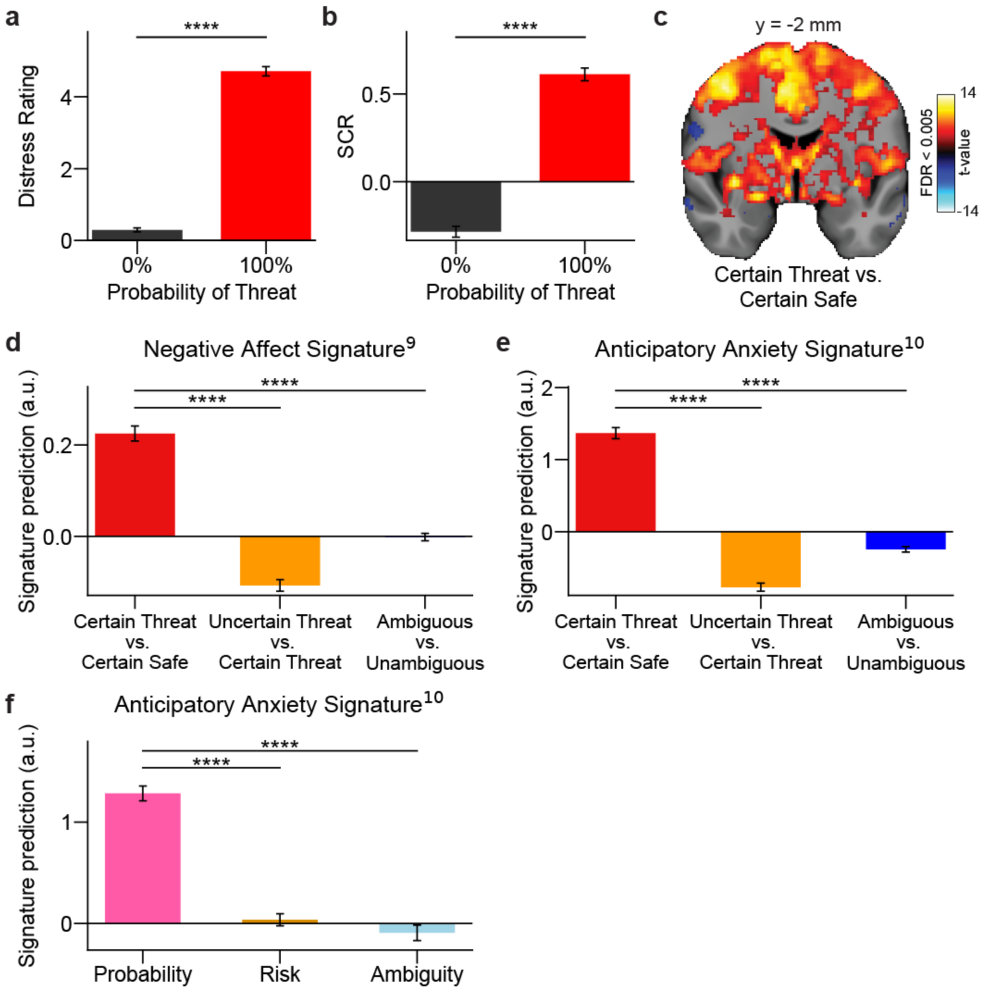


**Supplementary Fig. 1 |Certain-Threat cues elicit robust subjective, physiological, and neural responses, and these effects are not driven by uncertainty or ambiguity.**

**a**, Distress ratings for 0% and 100% Probability of Threat trials (N = 104). Distress was significantly higher for 100% than for 0% trials (linear mixed-effects model; z = 130.35, $p < 0.0001$).

**b**, Skin conductance response (SCR) for 0% and 100% Probability of Threat trials (N = 102). SCR was significantly higher for 100% than for 0% trials (linear mixed-effects model; z = 24.96, $p < 0.0001$).

**c**, Whole-brain activation for the certain threat vs. certain safe contrast ($q < 0.005$, FDR-corrected; N = 101). Coronal slice at MNI y = -2mm. Color scale, t-values.

**d,e,** Expression of the negative affect[9] (**d**) and anticipatory anxiety[10] (**e**) signatures across contrasts of interest (N = 101). In both, signature expression was greater for certain threat vs. certain safe than for uncertain threat vs. certain threat and ambiguous vs. unambiguous (linear mixed-effects model; **d**: uncertain threat vs. certain threat, z = −18.30; ambiguous vs. unambiguous, z = −12.47; **e**: uncertain threat vs. certain threat, z = −25.23; ambiguous vs. unambiguous, z = −19.03; all $p < 0.0001$).

**f**, Expression of the anticipatory anxiety signature[10] as a function of Probability, Risk, and Ambiguity (N = 101). Signature expression was significantly greater for Probability than for Risk and Ambiguity (linear mixed-effects model; Risk: z = −13.06; Ambiguity: z = −14.40; both $p < 0.0001$;  Corresponding to Figure 1k).

Error bars, SEM. All statistical tests were two-sided. ****$p < 0.0001$. a.u., arbitrary units.

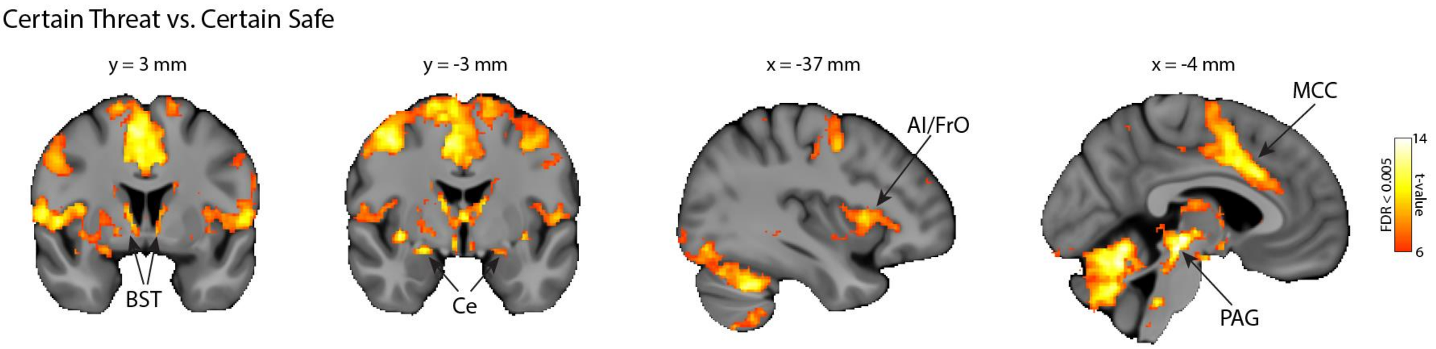


**Supplementary Fig. 2 | Whole-brain effects of Certain Threat vs. Certain Safe.**

Whole-brain group-level t-statistic maps for the Certain Threat versus Certain Safe contrast, overlaid on the mean skull-stripped T1 template and thresholded at FDR < 0.005 (t > 6; two-sided group-level one-sample t-test, N = 101), only positive effects are shown. Coronal slices (y = 3 mm, y = −3 mm) show clusters in the bed nucleus of the stria terminalis (BST) and central nucleus of the amygdala (Ce). Sagittal slices (x = −37 mm, x = −4 mm) show clusters in the anterior insula/frontal operculum (AI/FrO), midcingulate cortex (MCC), and periaqueductal gray (PAG).

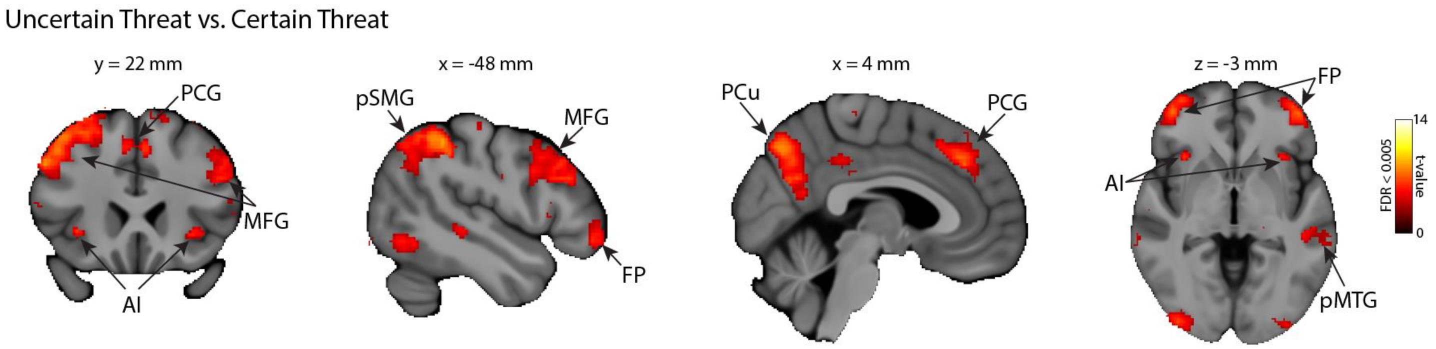


**Supplementary Fig. 3 | Whole-brain effects of Uncertain Threat vs. Certain Threat.**

Whole-brain group-level t-statistic maps for the Uncertain Threat versus Certain Threat contrast, overlaid on the mean skull-stripped T1 template and thresholded at FDR < 0.005 (two-sided group-level one-sample t-test, N = 101), only positive effects are shown. The coronal slice (y = 22 mm) shows clusters in the paracingulate gyrus (PCG), middle frontal gyrus (MFG), and anterior insula (AI). Sagittal slices (x = −48 mm, x = 4 mm) show clusters in the posterior supramarginal gyrus (pSMG), MFG, frontal pole (FP), precuneus (PCu), and PCG. The axial slice (z = −3 mm) shows bilateral clusters in the FP and AI, along with the posterior middle temporal gyrus (pMTG).

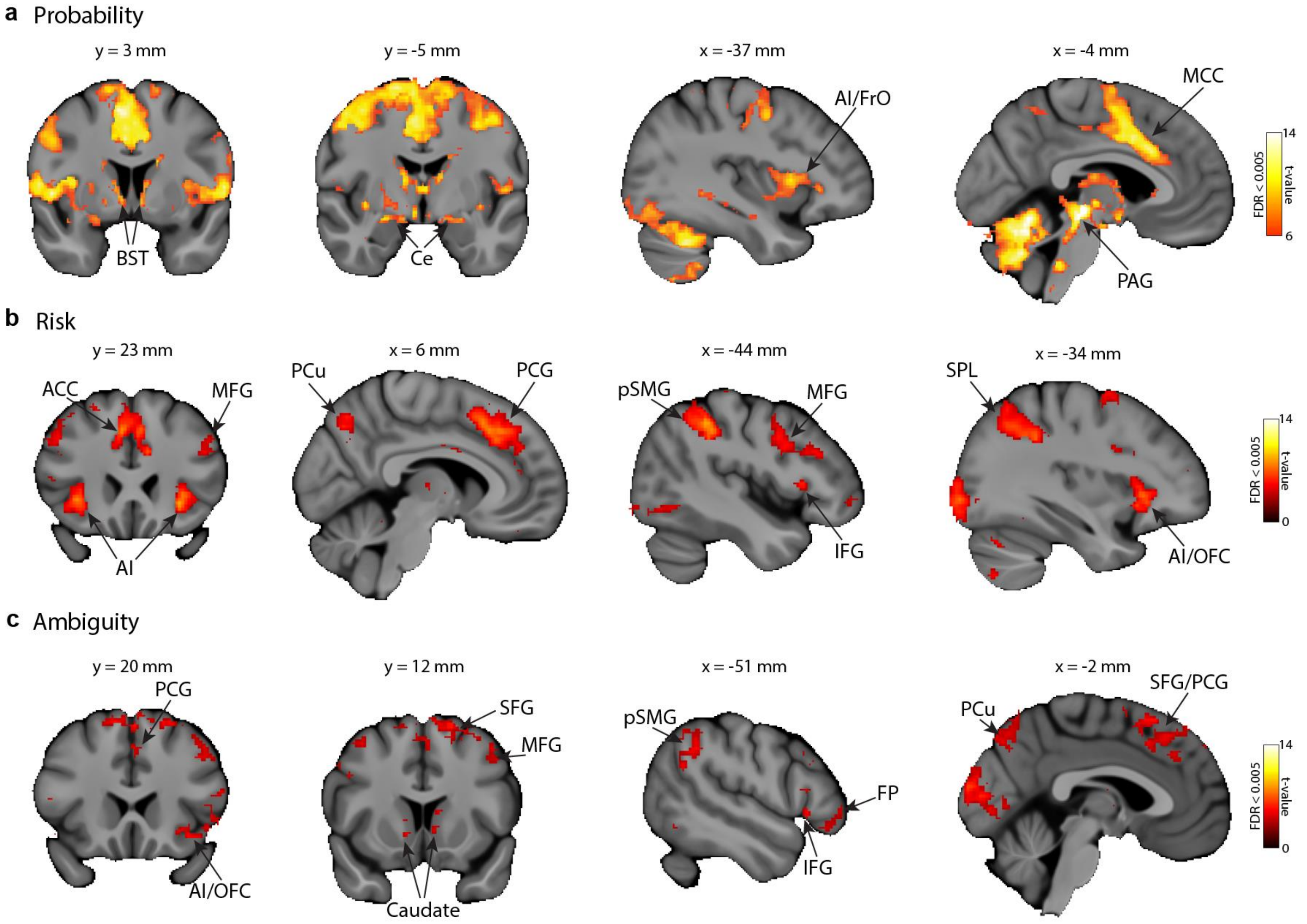


**Supplementary Fig. 4 | Whole-brain effects of Probability, Risk, and Ambiguity.**

Whole-brain group-level t-statistic maps overlaid on the mean skull-stripped T1 template, thresholded at FDR < 0.005 (two-sided group-level one-sample t-test, N = 101), only positive effects are shown.

**a**, Probability contrast, further thresholded at t > 6. Coronal slices (y = 3 mm, y = −5 mm) show clusters in the bed nucleus of the stria terminalis (BST) and central nucleus of the amygdala (Ce); sagittal slices (x = −37 mm, x = −4 mm) show clusters in the anterior insula/frontal operculum (AI/FrO), midcingulate cortex (MCC), and periaqueductal gray (PAG).

**b**, Risk contrast. Coronal slices (y = 23 mm, x = −44 mm, x = −34 mm) and a sagittal slice (x = 6 mm) show clusters in the anterior cingulate cortex (ACC), middle frontal gyrus (MFG), anterior insula (AI), precuneus (PCu), paracingulate gyrus (PCG), posterior supramarginal gyrus (pSMG), inferior frontal gyrus (IFG), superior parietal lobule (SPL), and anterior insula/orbitofrontal cortex (AI/OFC).

**c**, Ambiguity contrast. Coronal slices (y = 20 mm, y = 12 mm) and sagittal slices (x = −51 mm, x = −2 mm) show clusters in the paracingulate gyrus (PCG), anterior insula/orbitofrontal cortex (AI/OFC), superior frontal gyrus (SFG), middle frontal gyrus (MFG), posterior supramarginal gyrus (pSMG), inferior frontal gyrus (IFG), frontal pole (FP), precuneus (PCu), and caudate nucleus.

Color scale reflects t-value; note the higher lower-bound in **a** (6–14) versus **b,c** (0–14), reflecting the additional t > 6 threshold applied to the Probability map.

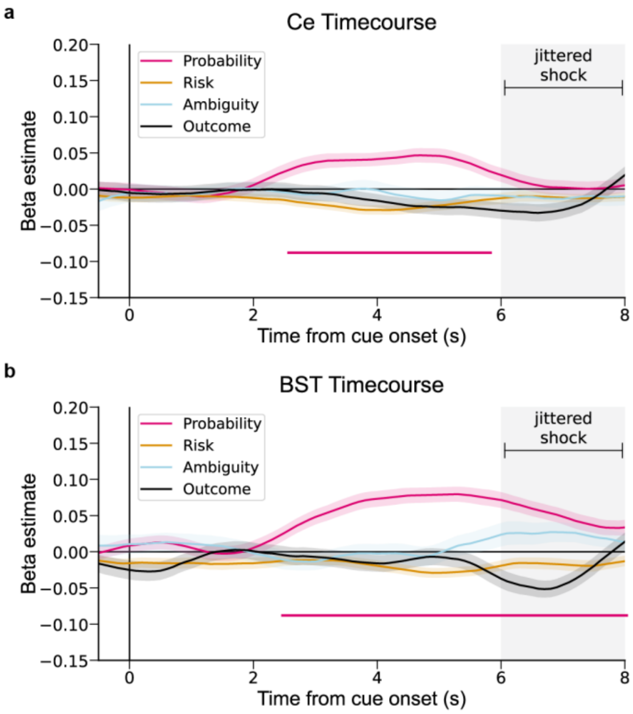


**Supplementary Fig. 5 | Probability signal dissociates from outcome response in Ce and BST.**

**a,b,** Timecourse of beta estimates in Ce (**a**) and BST (**b**). At each time point, mean ROI activity was modeled as a function of Probability, Risk, Ambiguity, and Outcome (shock vs. no shock), averaged across N = 101 subjects. Probability-driven effects during cue anticipation were dissociable from responses to aversive outcomes. Horizontal lines mark timepoints at which the beta estimate was significantly greater than zero (two-sided one-sample t-tests, FDR-corrected across predictors and timepoints within each ROI); line colors match the corresponding traces. Time is aligned to cue onset (0 s) and sampled every 100 ms (81 timepoints per line, 0–8 s window); shaded interval (6–8 s) denotes the jittered outcome period. Shaded areas around lines, SEM.

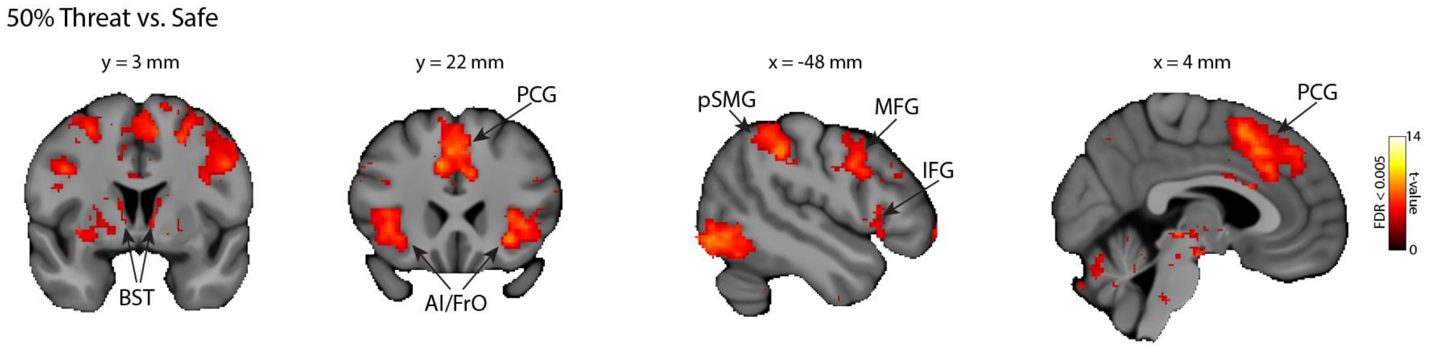


**Supplementary Fig. 6 | Whole-brain effects of 50% Threat vs. Safe.**

Whole-brain group-level t-statistic maps for the 50% Threat versus Safe contrast, overlaid on the mean skull-stripped T1 template and thresholded at FDR < 0.005 (two-sided group-level one-sample t-test, N = 101), only positive effects are shown. Coronal slices (y = 3 mm, y = 22 mm) show clusters in the bed nucleus of the stria terminalis (BST), paracingulate gyrus (PCG), and anterior insula/frontal operculum (AI/FrO). Sagittal slices (x = −48 mm, x = 4 mm) show clusters in the posterior supramarginal gyrus (pSMG), middle frontal gyrus (MFG), inferior frontal gyrus (IFG), and PCG.

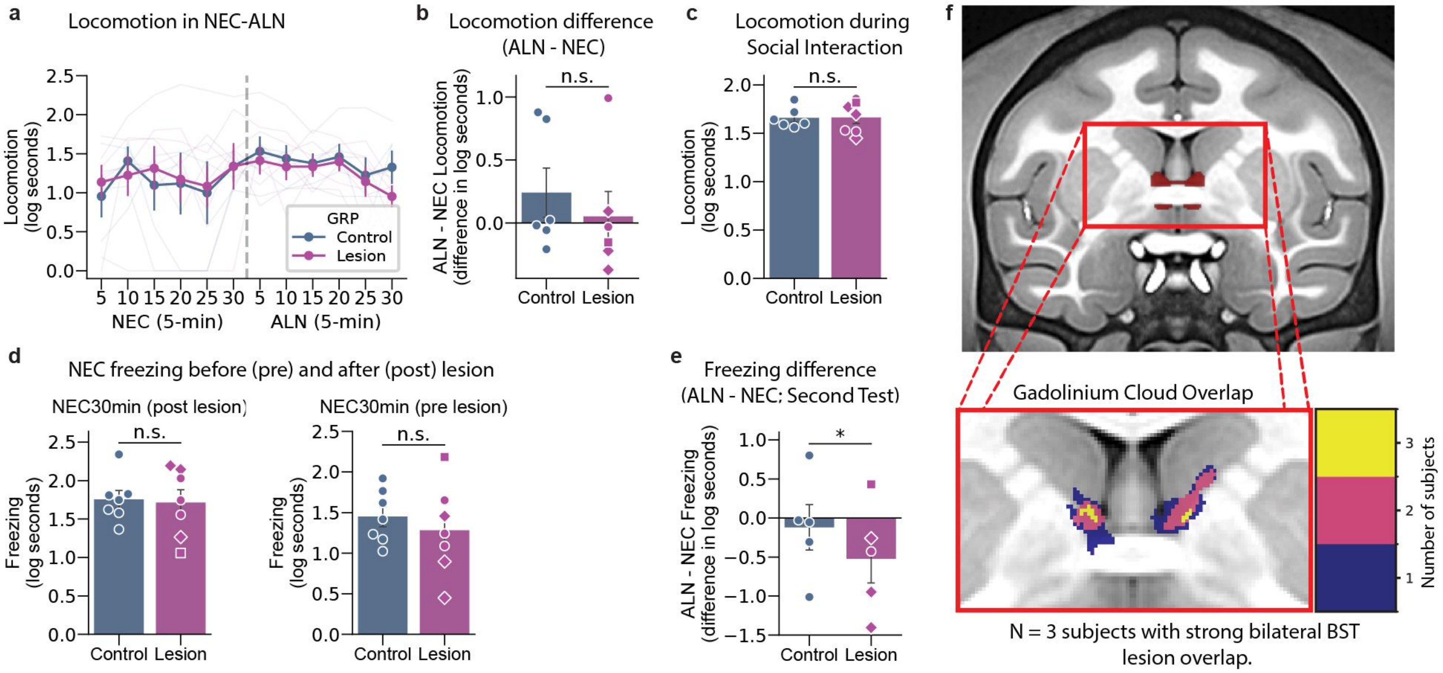


**Supplementary Fig. 7 | BST lesions do not impair general locomotion, and freezing effects replicate across independent assessments.**

**a**, Mean locomotion across 5-min bins during NEC and ALN portions of the HIP-Persistence paradigm in BST-lesioned (magenta, N = 6) and control (blue, N = 6) animals; thin lines show individual animals. Unlike freezing, locomotion did not differ between groups depending on condition (linear mixed-effects model; group × condition interaction: $z = 1.37$, $p = 0.171$), indicating that BST lesions did not impair general motor ability.

**b**, ALN−NEC difference in locomotion during the HIP-Persistence paradigm in BST-lesioned (magenta, N = 6) and control (blue, N = 6) groups; individual animals shown with markers matching **Fig. 4f** (diamonds: bilateral BST overlap; circles: unilateral overlap; squares: no significant overlap). Difference scores did not differ significantly between groups across the full sample (linear model; $t(5) = -0.85$, $p = 0.435$), nor when restricted to animals with confirmed bilateral BST overlap (linear model; $t(2) = -1.13$, $p = 0.377$).

**c**, Mean locomotion during the 30-min Social Interaction paradigm in BST-lesioned (magenta, N = 7) and control (blue, N = 6) animals; individual animals shown with markers matching **Fig. 4f.** Locomotion did not differ between groups across the full sample (linear mixed-effects model; group effect: $z = 0.28$, $p = 0.783$), nor when restricted to animals with confirmed bilateral BST overlap ($z = -0.28$, $p = 0.783$; N = 3), consistent with the absence of a motor confound observed in **a** and **b**.

**d**, Mean freezing during an independent assessment in the 30-min NEC-only context before lesion (left) and after lesion (right), in BST-lesioned (magenta, N = 7) and control (blue, N = 7) animals; individual animals shown with markers matching **Fig. 4f.** Freezing did not differ significantly between groups across the full sample either before lesion (linear mixed-effects model; group effect: $z = -1.36$, $p = 0.174$) or after lesion ($z = -0.37$, $p = 0.709$). When restricted to animals with confirmed bilateral BST overlap (N = 3), freezing differed significantly between groups before lesion ($z = -2.45$, $p = 0.014$) but not after lesion ($z = 0.73$, $p = 0.468$).

**e**, ALN−NEC difference in freezing during a second exposure to the HIP-Persistence paradigm, in BST-lesioned (magenta, N = 5) and control (blue, N = 5) groups; individual animals shown with markers matching **Fig. 4f**. Difference scores differed significantly between groups across the full sample (linear model; $t(4) = -3.03$, $p = 0.039$), but the effect was no longer significant when restricted to animals with confirmed bilateral BST overlap (linear model; $t(2) = -2.53$, $p = 0.127$).

**f**, BST ROI shown in NHP template space (top), zoomed to show the voxelwise overlap of gadolinium-visualized infusion clouds in the three BST-Lesion Group subjects with bilateral lesions, colored by the number of subjects with coverage at each voxel (bottom). Error bars (a-e), SEM. All statistical tests were two-sided. $^{*}p < 0.05$.

### Supplementary Tables

**Supplementary Table 1 | Statistical results for Figures 1–4 and Supplementary Figures 1 and 7**

Estimates, standard errors, 95% confidence intervals, test statistics and exact P values for every test reported for these figures. All tests are two-sided. Mixed-effects models were fitted by REML (statsmodels 0.14.4); z statistics are Wald tests of fixed-effect coefficients or of linear contrasts among them. Adjusted P values are given where a correction for multiple comparisons was applied.

| Panel | Measure | Effect / comparison | N | Estimate | SE | 95% CI | Test statistic | P | Adjusted P |
|---|---|---|---|---|---|---|---|---|---|
| **Figure 1 and Supplementary Figure 1** | | | | | | | | | |
| **A. Self-reported distress: trial-level model** *Linear mixed-effects model of trial-level distress ratings (0–6) with Probability and Risk (unambiguous trials), Ambiguity and Rational Probability (ambiguous trials) and trial type as fixed effects and a random intercept per subject; 10,776 ratings. Estimate, change in rating per unit of the rescaled predictor (a).* | | | | | | | | | |
| Fig. 1e | Distress | Probability | 104 | 2.160 | 0.015 | [2.131, 2.189] | z = 144.09 | $1.3 \times 10^{-4511}$ | — |
| Fig. 1f | Distress | Risk | 104 | 0.310 | 0.014 | [0.282, 0.338] | z = 21.73 | $1.1 \times 10^{-104}$ | — |
| Fig. 1g | Distress | Ambiguity | 104 | 0.374 | 0.025 | [0.325, 0.424] | z = 14.83 | $9.6 \times 10^{-50}$ | — |
| **B. Within-subject sensitivity: distress and SCR** *Subject-level regression coefficients (b) entered into a linear mixed-effects model with regressor as a fixed effect (Probability as reference) and a random intercept per subject. “Probability (vs 0)” tests the mean Probability coefficient against zero; other rows test the difference from Probability. Units: distress, rating points; SCR, within-subject z.* | | | | | | | | | |
| Fig. 1h | Distress | Probability (vs 0) | 104 | 2.158 | 0.048 | [2.064, 2.253] | z = 44.69 | $3.9 \times 10^{-436}$ | — |
| Fig. 1h | Distress | Risk vs Probability | 104 | −1.846 | 0.068 | [−1.980, −1.712] | z = −27.03 | $6.9 \times 10^{-161}$ | — |
| Fig. 1h | Distress | Ambiguity vs Probability | 104 | −1.780 | 0.068 | [−1.914, −1.646] | z = −26.06 | $1.1 \times 10^{-149}$ | — |
| Fig. 1i | SCR | Probability (vs 0) | 102 | 0.412 | 0.021 | [0.372, 0.453] | z = 19.82 | $2.2 \times 10^{-87}$ | — |
| Fig. 1i | SCR | Risk vs Probability | 102 | −0.556 | 0.029 | [−0.614, −0.498] | z = −18.89 | $1.4 \times 10^{-79}$ | — |
| Fig. 1i | SCR | Ambiguity vs Probability | 102 | −0.432 | 0.029 | [−0.490, −0.374] | z = −14.68 | $8.9 \times 10^{-49}$ | — |
| **C. Whole-brain neural signatures: parametric effects** *Subject-level pattern-expression scores (a.u.) for each parametric regressor, modelled with regressor as a fixed effect (Probability as reference) and a random intercept per subject.* | | | | | | | | | |
| Fig. 1k | Negative affect | Risk vs Probability | 101 | −0.203 | 0.023 | [−0.247, −0.158] | z = −8.96 | $3.4 \times 10^{-19}$ | — |
| Fig. 1k | Negative affect | Ambiguity vs Probability | 101 | −0.203 | 0.023 | [−0.247, −0.158] | z = −8.95 | $3.4 \times 10^{-19}$ | — |
| SF 1f | Anticipatory anxiety | Risk vs Probability | 101 | −1.247 | 0.095 | [−1.435, −1.060] | z = −13.06 | $5.5 \times 10^{-39}$ | — |
| SF 1f | Anticipatory anxiety | Ambiguity vs Probability | 101 | −1.375 | 0.095 | [−1.563, −1.188] | z = −14.40 | $5.0 \times 10^{-47}$ | — |
| **D. Certain threat vs certain safety** *Linear mixed-effects model of trial-level responses on certain trials (100% vs 0% threat probability) with a random intercept per subject; 2,901 ratings (a) and 3,609 SCR trials (b).* | | | | | | | | | |
| SF 1a | Distress | Certain threat (100%) vs certain safety (0%) | 104 | 4.396 | 0.034 | [4.330, 4.462] | z = 130.35 | $1.3 \times 10^{-3692}$ | — |
| SF 1b | SCR | Certain threat (100%) vs certain safety (0%) | 102 | 0.880 | 0.035 | [0.811, 0.949] | z = 24.96 | $1.6 \times 10^{-137}$ | — |
| **E. Whole-brain neural signatures: categorical contrasts** *Subject-level pattern-expression scores (a.u.) for three first-level contrasts, modelled with contrast as a fixed effect (certain threat vs certain safety as reference) and a random intercept per subject.* | | | | | | | | | |
| SF 1d | Negative affect | Uncertain threat vs certain threat | 101 | −0.332 | 0.018 | [−0.368, −0.297] | z = −18.30 | $8.2 \times 10^{-75}$ | — |
| SF 1d | Negative affect | Ambiguous vs unambiguous trials | 101 | −0.226 | 0.018 | [−0.262, −0.191] | z = −12.47 | $1.1 \times 10^{-35}$ | — |
| SF 1e | Anticipatory anxiety | Uncertain threat vs certain threat | 101 | −2.137 | 0.085 | [−2.303, −1.971] | z = −25.23 | $2.0 \times 10^{-140}$ | — |
| SF 1e | Anticipatory anxiety | Ambiguous vs unambiguous trials | 101 | −1.612 | 0.085 | [−1.778, −1.446] | z = −19.03 | $8.9 \times 10^{-81}$ | — |

| Panel | Measure | Effect / comparison | N | Estimate | SE | 95% CI | Test statistic | P | Adjusted P |
|---|---|---|---|---|---|---|---|---|---|
| **Figure 2** | | | | | | | | | |
| **F. Ce and BST: Probability, Risk and Ambiguity** *Mean ROI parameter estimates (a.u.) from the parametric first-level model, analyzed with a linear mixed-effects model (regressor × region as fixed effects, random intercept per subject) and linear contrasts of its coefficients. Adjusted P, Benjamini–Hochberg FDR across 10 planned contrasts (c).* | | | | | | | | | |
| Fig. 2a | Ce | Probability (vs 0) | 101 | 0.137 | 0.020 | [0.099, 0.176] | z = 7.00 | $2.6 \times 10^{-12}$ | $2.6 \times 10^{-11}$ |
| Fig. 2a | BST | Probability (vs 0) | 101 | 0.122 | 0.020 | [0.084, 0.160] | z = 6.22 | $5.0 \times 10^{-10}$ | $2.5 \times 10^{-9}$ |
| Fig. 2a | Ce | Risk (vs 0) | 101 | −0.022 | 0.020 | [−0.061, 0.016] | z = −1.13 | 0.258 | 0.287 |
| Fig. 2a | BST | Risk (vs 0) | 101 | 0.032 | 0.020 | [−0.006, 0.071] | z = 1.64 | 0.101 | 0.169 |
| Fig. 2a | Ce | Ambiguity (vs 0) | 101 | −0.025 | 0.020 | [−0.064, 0.013] | z = −1.30 | 0.194 | 0.270 |
| Fig. 2a | BST | Ambiguity (vs 0) | 101 | 0.008 | 0.020 | [−0.031, 0.046] | z = 0.40 | 0.686 | 0.686 |
| Fig. 2a | Ce | Risk vs Probability | 101 | −0.160 | 0.027 | [−0.212, −0.107] | z = −5.91 | $3.4 \times 10^{-9}$ | $8.5 \times 10^{-9}$ |
| Fig. 2a | BST | Risk vs Probability | 101 | −0.090 | 0.027 | [−0.143, −0.037] | z = −3.33 | $8.7 \times 10^{-4}$ | — |
| Fig. 2a | Ce | Ambiguity vs Probability | 101 | −0.163 | 0.027 | [−0.216, −0.110] | z = −6.03 | $1.6 \times 10^{-9}$ | $5.4 \times 10^{-9}$ |
| Fig. 2a | BST | Ambiguity vs Probability | 101 | −0.114 | 0.027 | [−0.167, −0.061] | z = −4.23 | $2.4 \times 10^{-5}$ | — |
| Fig. 2a | Ce vs BST | Region × (Risk vs Probability) | 101 | −0.070 | 0.038 | [−0.144, 0.005] | z = −1.82 | 0.068 | — |
| Fig. 2a | Ce vs BST | Region × (Ambiguity vs Probability) | 101 | −0.049 | 0.038 | [−0.124, 0.026] | z = −1.28 | 0.202 | — |
| **Figure 3** | | | | | | | | | |
| **G. Ce vs BST response to 50% vs 0% threat** *Mean ROI parameter estimates (a.u.) for the 50% vs 0% contrast, analyzed with a linear mixed-effects model with region as a fixed effect (BST as reference) and a random intercept per subject. Estimate, Ce − BST.* | | | | | | | | | |
| Fig. 3b | ROI β | Ce vs BST | 101 | −0.088 | 0.030 | [−0.147, −0.030] | z = −2.95 | 0.003 | — |
| **H. Sigmoid fits: Ce vs BST** *Two-sided Mann–Whitney U tests comparing bootstrap distributions of the sigmoid midpoint (x0, % threat probability) and slope (k) between BST and Ce (1,000 half-sample bootstrap fits per region). Estimate, difference in bootstrap medians (BST − Ce) (d).* | | | | | | | | | |
| Fig. 3h,i | Sigmoid parameter | Midpoint (x0): BST vs Ce | 1,000 / 1,000 fits | −46.840 | — | — | U = 193,918 | $3.4 \times 10^{-124}$ | — |
| Fig. 3h,i | Sigmoid parameter | Slope (k): BST vs Ce | 1,000 / 1,000 fits | 0.004 | — | — | U = 543,695 | $7.2 \times 10^{-4}$ | — |
| **I. BST − Ce change across adjacent probability bins** *Per-subject BST − Ce difference scores (a.u.) for four adjacent probability-bin transitions. Omnibus: one-way ANOVA. Planned comparisons: lowest-probability transition (0% → 10–30%) vs each other transition, as linear contrasts of a linear mixed-effects model (transition as fixed effect, random intercept per subject); estimate, lowest transition minus the comparison transition; adjusted P, Holm across the 3 comparisons (e).* | | | | | | | | | |
| Fig. 3k | BST − Ce | Effect of transition (omnibus) | 101 | — | — | — | $F(3,400) = 2.61$ | 0.051 | — |
| Fig. 3k | BST − Ce | 0% → 10–30% vs 10–30% → 40–60% | 101 | 0.017 | 0.030 | [−0.041, 0.075] | z = 0.56 | 0.574 | 0.574 |
| Fig. 3k | BST − Ce | 0% → 10–30% vs 40–60% → 70–90% | 101 | 0.077 | 0.030 | [0.019, 0.135] | z = 2.59 | 0.010 | 0.029 |
| Fig. 3k | BST − Ce | 0% → 10–30% vs 70–90% → 100% | 101 | 0.046 | 0.030 | [−0.012, 0.104] | z = 1.57 | 0.117 | 0.234 |

| Panel | Measure | Effect / comparison | N | Estimate | SE | 95% CI | Test statistic | P | Adjusted P |
|---|---|---|---|---|---|---|---|---|---|
| **Figure 4 and Supplementary Figure 7** | | | | | | | | | |
| **J. HIP-Persistence freezing: lesion effects by context**<br>*Linear mixed-effects model of freezing (log seconds per 5-min bin) with group (lesion vs control), context (NEC vs ALN-following-NEC) and their interaction, plus time bin, as fixed effects and a random intercept per cage-mate pair. Estimates, lesion − control (log s); interaction, lesion effect in NEC minus lesion effect in ALN (f).* | | | | | | | | | |
| Fig. 4e | Freezing | Group × context interaction | 6 L / 6 C | 0.443 | 0.206 | [0.039, 0.847] | z = 2.15 | 0.031 | — |
| Fig. 4e | Freezing | Lesion vs control, ALN | 6 L / 6 C | −0.458 | 0.146 | [−0.743, −0.172] | z = −3.14 | 0.002 | — |
| Fig. 4e | Freezing | Lesion vs control, NEC | 6 L / 6 C | −0.015 | 0.146 | [−0.300, 0.271] | z = −0.10 | 0.920 | — |
| Fig. 4e | Freezing | Lesion vs control, ALN, bilateral subset | 3 L / 6 C | −1.060 | 0.172 | [−1.397, −0.723] | z = −6.16 | $7.2 \times 10^{-10}$ | — |
| **K. HIP-Persistence freezing: ALN − NEC difference**<br>*Ordinary least-squares regression of each animal's mean ALN − NEC difference in freezing (log s) on group, with cage-mate pair as a fixed effect. Estimate, lesion − control (f).* | | | | | | | | | |
| Fig. 4g | Freezing | ALN − NEC difference: lesion vs control | 6 L / 6 C | −0.443 | 0.194 | [−0.941, 0.055] | t(5) = −2.29 | 0.071 | — |
| Fig. 4g | Freezing | ALN − NEC difference: lesion vs control, bilateral subset | 3 L / 6 C | −0.711 | 0.124 | [−1.244, −0.178] | t(2) = −5.74 | 0.029 | — |
| **L. Social interaction freezing**<br>*Linear mixed-effects model of freezing (log s per 5-min bin) during the 30-min social interaction with group and time bin as fixed effects and a random intercept per cage-mate pair. Estimate, lesion − control (f).* | | | | | | | | | |
| Fig. 4i | Freezing | Lesion vs control | 7 L / 6 C | −0.394 | 0.095 | [−0.581, −0.208] | z = −4.14 | $3.5 \times 10^{-5}$ | — |
| Fig. 4i | Freezing | Lesion vs control, bilateral subset | 3 L / 6 C | −0.527 | 0.121 | [−0.765, −0.289] | z = −4.35 | $1.4 \times 10^{-5}$ | — |
| **M. Locomotion**<br>*Same models as for freezing, applied to locomotion (log s): HIP-Persistence group × context interaction (SF 7a; model as in Fig. 4e), HIP-Persistence ALN − NEC difference (SF 7b; model as in Fig. 4g) and social interaction (SF 7c; model as in Fig. 4i). Estimates, lesion − control (f).* | | | | | | | | | |
| SF 7a | Locomotion | Group × context interaction | 6 L / 6 C | 0.189 | 0.138 | [−0.082, 0.460] | z = 1.37 | 0.171 | — |
| SF 7b | Locomotion | ALN − NEC difference: lesion vs control | 6 L / 6 C | −0.189 | 0.223 | [−0.762, 0.384] | t(5) = −0.85 | 0.435 | — |
| SF 7b | Locomotion | ALN − NEC difference: lesion vs control, bilateral subset | 3 L / 6 C | −0.460 | 0.409 | [−2.219, 1.298] | t(2) = −1.13 | 0.377 | — |
| SF 7c | Locomotion | Lesion vs control | 7 L / 6 C | 0.014 | 0.051 | [−0.086, 0.114] | z = 0.28 | 0.783 | — |
| SF 7c | Locomotion | Lesion vs control, bilateral subset | 3 L / 6 C | −0.019 | 0.068 | [−0.153, 0.115] | z = −0.28 | 0.783 | — |
| **N. NEC-only freezing before and after lesion**<br>*Independent 30-min NEC-only assessment before and after surgery; model as in Fig. 4i. Estimate, lesion − control (f).* | | | | | | | | | |
| SF 7d | Freezing | Before lesion: lesion vs control | 7 L / 7 C | −0.169 | 0.124 | [−0.413, 0.075] | z = −1.36 | 0.174 | — |
| SF 7d | Freezing | Before lesion: lesion vs control, bilateral subset | 3 L / 7 C | −0.465 | 0.190 | [−0.836, −0.093] | z = −2.45 | 0.014 | — |
| SF 7d | Freezing | After lesion: lesion vs control | 7 L / 7 C | −0.043 | 0.115 | [−0.269, 0.183] | z = −0.37 | 0.709 | — |
| SF 7d | Freezing | After lesion: lesion vs control, bilateral subset | 3 L / 7 C | 0.102 | 0.141 | [−0.174, 0.379] | z = 0.73 | 0.468 | — |
| **O. Second HIP-Persistence exposure: ALN − NEC difference**<br>*Model as in Fig. 4g, for a second exposure to the HIP-Persistence paradigm. Estimate, lesion − control (f).* | | | | | | | | | |
| SF 7e | Freezing | ALN − NEC difference: lesion vs control | 5 L / 5 C | −0.401 | 0.133 | [−0.769, −0.033] | t(4) = −3.03 | 0.039 | — |
| SF 7e | Freezing | ALN − NEC difference: lesion vs control, bilateral subset | 3 L / 5 C | −0.505 | 0.200 | [−1.365, 0.355] | t(2) = −2.53 | 0.127 | — |

P values ≥ 0.001 are given to three decimals; smaller P values are exact values computed from the test statistic. N, number of human participants (Fig. 3h,i: number of bootstrap fits per region; Fig. 4 and SF 7: see f). SCR, skin conductance response; SF, Supplementary Figure; ROI, region of interest; Ce, central nucleus of the amygdala; BST, bed nucleus of the stria terminalis; a.u., arbitrary units. Fig. 1l and Supplementary Fig. 5 timecourse tests are shown graphically (FDR-corrected across predictors and timepoints; Methods) and are not tabulated.

(a) Before model fitting, each parametric predictor was min–max scaled to 0–1 within its trial type, mean-centered and doubled, so that all predictors span a common range of 2 (Methods). Rational Probability was included as a covariate on ambiguous trials and is not interpreted.

(b) Per-subject ordinary least-squares coefficients for Probability, Risk and Ambiguity, with predictors rescaled within each subject (Methods).

(c) The FDR family comprised 10 planned contrasts: the eight Fig. 2a contrasts with an adjusted P above, plus two Ce vs BST comparisons not reported in the main text (Risk, Ce vs BST: estimate −0.054 [−0.107, −0.001], z = −2.01, P = 0.044, adjusted P = 0.088; Ambiguity, Ce vs BST: estimate −0.033 [−0.086, 0.019], z = −1.24, P = 0.216, adjusted P = 0.270). BST Risk vs Probability and BST Ambiguity vs Probability, and the two Region × facet interactions, are model coefficients and were not part of the FDR family.

(d) Each bootstrap iteration resampled 50 of the 101 participants with replacement; all 1,000 fits converged in both regions. Bootstrap medians: x0, BST 16.1 and Ce 62.9; k, BST 0.077 and Ce 0.073.

(e) Omnibus: one-way ANOVA (ordinary least squares) on 404 difference scores (four per participant); partial $\eta^2 = 0.019$.

(f) Non-human primates: N, number of lesioned (L) and control (C) animals; controls were the lesioned animals' cage-mates. Bilateral subset, analyses restricted to the 3 lesioned animals with bilateral BST overlap (Fig. 4f), compared with all controls. HIP, human intruder paradigm; NEC, no-eye-contact; ALN, alone period following NEC. t tests have residual degrees of freedom in parentheses.

#### Supplementary Table 2 | Brain regions showing greater activation for Certain Threat vs. Certain Safe

Whole-brain analysis contrasting cue-evoked activation during Certain Threat versus Certain Safe trials (Certain Threat > Certain Safe), thresholded at FDR-corrected q < 0.005 (positive contrast only). $mm^3$, volume of the contiguous suprathreshold cluster; Label, anatomical region containing the local activation peak, identified using the probabilistic Harvard-Oxford cortical and subcortical atlases; L, left hemisphere; R, right hemisphere; t, peak voxel t-statistic; x, y, z, peak voxel coordinates in MNI space (mm). Clusters are ordered by descending volume, and peaks within each cluster by descending t-statistic.

| $mm^3$ | Label | t | x | y | z |
|---|---|---|---|---|---|
| **677,912** | R Brain-Stem | 14.76 | 6 | -28 | -10 |
| | L Brain-Stem | 14.45 | -6 | -28 | -12 |
| | R Postcentral Gyrus | 14.12 | 38 | -24 | 60 |
| | L Cingulate Gyrus, anterior division | 13.46 | -8 | 12 | 36 |
| | R Precentral Gyrus | 13.05 | 36 | -8 | 64 |
| | R Juxtapositional Lobule Cortex (formerly Supplementary Motor Cortex) | 12.88 | 8 | 2 | 56 |
| | R Cingulate Gyrus, anterior division | 12.72 | 10 | 10 | 38 |
| | R Paracingulate Gyrus | 12.39 | 6 | 8 | 46 |
| | R Central Opercular Cortex | 12.23 | 50 | 4 | 4 |
| | R Occipital Fusiform Gyrus | 12.01 | 24 | -88 | -12 |
| | R Superior Frontal Gyrus | 11.85 | 16 | -4 | 70 |
| | R Lateral Occipital Cortex, inferior division | 11.74 | 34 | -82 | -14 |
| | R Left Thalamus | 11.61 | 0 | -8 | 8 |
| | R Amygdala | 11.38 | 22 | 0 | -12 |
| | L Juxtapositional Lobule Cortex (formerly Supplementary Motor Cortex) | 11.36 | -2 | 6 | 48 |
| | L Precentral Gyrus | 11.26 | -46 | -10 | 50 |
| | R Cingulate Gyrus, posterior division | 11.24 | 10 | -18 | 44 |
| | R Occipital Pole | 10.97 | 14 | -92 | -12 |
| | R Frontal Operculum Cortex | 10.81 | 40 | 12 | 4 |
| | L Thalamus | 10.72 | -4 | -16 | -2 |
| | R Superior Parietal Lobule | 10.54 | 22 | -48 | 66 |
| | L Occipital Fusiform Gyrus | 10.45 | -26 | -76 | -18 |
| | R Lingual Gyrus | 10.33 | 10 | -84 | -12 |
| | R Thalamus | 10.33 | 2 | -20 | 10 |
| | L Frontal Operculum Cortex | 10.25 | -38 | 12 | 6 |
| | L Superior Frontal Gyrus | 10.22 | -14 | -6 | 68 |
| | L Caudate | 10.21 | -12 | -2 | 14 |
| | R Caudate | 10.19 | 10 | 0 | 12 |
| | R Insular Cortex | 10.00 | 36 | 4 | 10 |
| | L Lateral Occipital Cortex, inferior division | 9.85 | -38 | -78 | -12 |
| | R Frontal Orbital Cortex | 9.70 | 38 | 30 | 0 |
| | R Parietal Operculum Cortex | 9.45 | 58 | -22 | 18 |
| | L Middle Frontal Gyrus | 9.14 | -36 | -4 | 58 |
| | R Putamen | 8.77 | 24 | 10 | -4 |
| | L Supramarginal Gyrus, anterior division | 8.71 | -66 | -28 | 20 |
| | L Occipital Pole | 8.56 | -26 | -98 | -2 |
| | R Supramarginal Gyrus, anterior division | 8.37 | 62 | -26 | 28 |
| | R Pallidum | 8.34 | 16 | 4 | 0 |
| | L Central Opercular Cortex | 8.32 | -44 | 2 | 8 |

| | | | | | |
|---|---|---|---|---|---|
| | L Insular Cortex | 8.22 | -34 | 2 | 10 |
| | L Amygdala | 8.16 | -22 | -2 | -14 |
| | L Precuneus Cortex | 8.06 | -12 | -46 | 50 |
| | L Superior Parietal Lobule | 7.97 | -20 | -50 | 64 |
| | R Precuneus Cortex | 7.77 | 12 | -52 | 58 |
| | L Amygdala | 7.75 | -24 | -6 | -12 |
| | L Parietal Operculum Cortex | 7.68 | -56 | -38 | 28 |
| | L Lateral Occipital Cortex, superior division | 7.56 | -30 | -72 | 22 |
| | R Frontal Pole | 7.43 | 34 | 44 | 26 |
| | L Frontal Pole | 7.28 | -32 | 48 | 30 |
| | L Intracalcarine Cortex | 7.25 | -22 | -64 | 4 |
| | R Supramarginal Gyrus, posterior division | 7.24 | 66 | -36 | 28 |
| | L Frontal Orbital Cortex | 7.24 | -28 | 20 | -12 |
| | R Lateral Occipital Cortex, superior division | 7.11 | 30 | -86 | 14 |
| | R Middle Temporal Gyrus, temporooccipital part | 7.08 | 58 | -60 | 2 |
| | L Pallidum | 6.91 | -20 | -10 | -4 |
| | L Putamen | 6.65 | -26 | 4 | -4 |
| | R Temporal Occipital Fusiform Cortex | 6.48 | 46 | -58 | -18 |
| | L Lingual Gyrus | 6.27 | -4 | -88 | -12 |
| | L Inferior Temporal Gyrus, temporooccipital part | 6.17 | -50 | -48 | -26 |
| | R Intracalcarine Cortex | 6.15 | 24 | -62 | 6 |
| | L Hippocampus | 5.92 | -26 | -18 | -12 |
| | R Inferior Temporal Gyrus, temporooccipital part | 5.84 | 46 | -54 | -10 |
| | L Middle Temporal Gyrus, temporooccipital part | 5.79 | -50 | -56 | 4 |
| | L Postcentral Gyrus | 5.40 | -50 | -22 | 36 |
| | L Cuneal Cortex | 5.18 | -18 | -74 | 28 |
| | R Hippocampus | 5.10 | 26 | -16 | -12 |
| | R Superior Temporal Gyrus, posterior division | 5.09 | 52 | -18 | -6 |
| | L Temporal Fusiform Cortex, posterior division | 4.92 | -40 | -36 | -22 |
| | L Temporal Occipital Fusiform Cortex | 4.60 | -40 | -48 | -18 |
| | R Inferior Frontal Gyrus, pars opercularis | 4.43 | 42 | 14 | 24 |
| | R Inferior Temporal Gyrus, posterior division | 3.89 | 48 | -30 | -22 |
| | R Temporal Pole | 3.87 | 38 | 8 | -22 |
| | R Cuneal Cortex | 3.77 | 6 | -80 | 40 |
| | L Temporal Pole | 3.31 | -58 | 10 | -10 |
| **240** | R Frontal Pole | 4.46 | 24 | 68 | -6 |
| **136** | L Frontal Medial Cortex | 3.95 | -4 | 38 | -28 |
| **136** | R Temporal Fusiform Cortex, posterior division | 4.73 | 38 | -18 | -24 |
| **104** | R Temporal Pole | 4.46 | 38 | 8 | -44 |
| **72** | R Intracalcarine Cortex | 3.59 | 10 | -72 | 12 |
| **64** | R Superior Frontal Gyrus | 4.13 | 6 | 50 | 30 |
| **56** | R Inferior Temporal Gyrus, posterior division | 3.54 | 58 | -20 | -34 |
| **48** | L Temporal Pole | 3.61 | -36 | 2 | -44 |
| | L Temporal Fusiform Cortex, anterior division | 3.57 | -36 | 0 | -40 |
| **48** | L Inferior Frontal Gyrus, pars opercularis | 3.83 | -44 | 16 | 18 |
| **48** | L Cuneal Cortex | 3.68 | -14 | -74 | 20 |
| **48** | R Lateral Occipital Cortex, superior division | 3.33 | 22 | -68 | 50 |
| **40** | R Inferior Temporal Gyrus, anterior division | 3.54 | 46 | 2 | -42 |
| **32** | R Temporal Pole | 3.40 | 30 | 10 | -42 |
| **32** | R Inferior Temporal Gyrus, posterior division | 3.67 | 50 | -22 | -28 |

| | | | | | |
|---|---|---|---|---|---|
| **32** | L Supramarginal Gyrus, posterior division | 4.22 | -54 | -44 | 18 |
| **24** | L Temporal Pole | 4.01 | -30 | 4 | -48 |
| **24** | L Temporal Pole | 4.32 | -32 | 4 | -40 |
| **24** | L Inferior Temporal Gyrus, posterior division | 3.60 | -44 | -16 | -38 |
| **24** | R Inferior Temporal Gyrus, posterior division | 3.83 | 46 | -20 | -32 |
| **24** | R Parahippocampal Gyrus, anterior division | 3.61 | 24 | -18 | -32 |
| **24** | R Frontal Pole | 3.41 | 12 | 50 | -22 |
| **24** | L Frontal Orbital Cortex | 4.12 | -38 | 32 | -20 |
| **16** | L Temporal Fusiform Cortex, anterior division | 3.31 | -30 | -4 | -42 |
| **16** | R Brain-Stem | 3.29 | 8 | -24 | -34 |
| **16** | R Frontal Medial Cortex | 3.59 | 6 | 44 | -28 |
| **16** | R Frontal Pole | 3.58 | 42 | 36 | -18 |
| **16** | L Frontal Pole | 3.47 | -22 | 70 | -4 |
| **16** | R Cingulate Gyrus, posterior division | 3.47 | 0 | -26 | 26 |
| **8** | R Brain-Stem | 3.30 | 8 | -32 | -52 |
| **8** | L Inferior Temporal Gyrus, anterior division | 3.19 | -44 | -4 | -44 |
| **8** | R Inferior Temporal Gyrus, posterior division | 3.46 | 48 | -16 | -38 |
| **8** | R Brain-Stem | 3.21 | 10 | -16 | -38 |
| **8** | R Brain-Stem | 3.30 | 22 | -34 | -32 |
| **8** | R Inferior Temporal Gyrus, posterior division | 3.21 | 50 | -12 | -32 |
| **8** | L Inferior Temporal Gyrus, posterior division | 3.27 | -46 | -20 | -26 |
| **8** | R Frontal Pole | 3.31 | 20 | 44 | -22 |
| **8** | R Inferior Temporal Gyrus, posterior division | 3.42 | 52 | -34 | -20 |
| **8** | R Frontal Pole | 3.28 | 30 | 34 | -20 |
| **8** | R Frontal Pole | 3.61 | 22 | 54 | -20 |
| **8** | L Frontal Orbital Cortex | 3.41 | -20 | 26 | -18 |
| **8** | L Frontal Orbital Cortex | 3.23 | -42 | 30 | -18 |
| **8** | R Frontal Pole | 3.29 | 30 | 52 | -18 |
| **8** | L Frontal Pole | 3.33 | -28 | 40 | -16 |
| **8** | L Frontal Pole | 3.29 | -26 | 52 | -16 |
| **8** | L Frontal Pole | 3.43 | -16 | 58 | -12 |
| **8** | R Frontal Pole | 3.20 | 24 | 58 | -10 |
| **8** | L Frontal Pole | 3.42 | -22 | 58 | -10 |
| **8** | L Intracalcarine Cortex | 3.21 | -6 | -74 | 12 |
| **8** | L Cuneal Cortex | 3.29 | -6 | -78 | 22 |
| **8** | R Cuneal Cortex | 3.65 | 8 | -76 | 24 |
| **8** | L Superior Frontal Gyrus | 3.21 | -4 | 32 | 52 |

#### Supplementary Table 3 | Brain regions showing greater activation for Uncertain Threat vs. Certain Threat

Whole-brain analysis contrasting cue-evoked activation during Uncertain Threat (Unambiguous + Ambiguous) versus Certain Threat (0% and 100% threat) trials (Uncertain Threat > Certain Threat), thresholded at FDR-corrected $q < 0.005$ (positive contrast only). $mm^3$, volume of the contiguous suprathreshold cluster; Label, anatomical region containing the local activation peak, identified using the probabilistic Harvard-Oxford cortical and subcortical atlases; L, left hemisphere; R, right hemisphere; t, peak voxel t-statistic; x, y, z, peak voxel coordinates in MNI space (mm). Clusters are ordered by descending volume, and peaks within each cluster by descending t-statistic.

| $mm^3$ | Label | t | x | y | z |
|---|---|---|---|---|---|
| **65,232** | R Supramarginal Gyrus, posterior division | 11.58 | 48 | -38 | 48 |
| | R Lateral Occipital Cortex, superior division | 10.79 | 42 | -60 | 50 |
| | R Angular Gyrus | 9.65 | 42 | -54 | 54 |
| | R Precuneus Cortex | 8.76 | 6 | -68 | 46 |
| | L Supramarginal Gyrus, posterior division | 8.73 | -46 | -42 | 48 |
| | L Lateral Occipital Cortex, superior division | 8.13 | -26 | -68 | 48 |
| | R Lateral Occipital Cortex, inferior division | 6.53 | 38 | -90 | -2 |
| | L Angular Gyrus | 6.39 | -48 | -56 | 52 |
| | R Occipital Pole | 5.89 | 36 | -90 | 10 |
| | L Precuneus Cortex | 5.51 | -4 | -72 | 46 |
| **19,856** | R Middle Frontal Gyrus | 8.61 | 30 | 20 | 54 |
| | R Frontal Pole | 7.03 | 46 | 38 | 22 |
| | R Superior Frontal Gyrus | 6.55 | 22 | 28 | 58 |
| **9,264** | R Inferior Temporal Gyrus, temporooccipital part | 8.00 | 60 | -52 | -12 |
| | R Middle Temporal Gyrus, temporooccipital part | 7.24 | 58 | -42 | -10 |
| | R Middle Temporal Gyrus, posterior division | 6.23 | 60 | -22 | -20 |
| | R Inferior Temporal Gyrus, posterior division | 5.69 | 58 | -26 | -20 |
| | R Lateral Occipital Cortex, inferior division | 5.38 | 58 | -66 | -10 |
| **7,768** | R Frontal Pole | 7.58 | 42 | 50 | -10 |
| **6,896** | L Middle Frontal Gyrus | 6.60 | -46 | 30 | 28 |
| **6,624** | L Paracingulate Gyrus | 7.82 | -8 | 18 | 46 |
| | R Paracingulate Gyrus | 7.38 | 6 | 36 | 40 |
| | L Superior Frontal Gyrus | 5.46 | -4 | 12 | 58 |
| | R Superior Frontal Gyrus | 3.83 | 4 | 38 | 54 |
| **5,208** | L Frontal Pole | 7.50 | -42 | 52 | -4 |
| **4,536** | L Middle Temporal Gyrus, posterior division | 5.52 | -60 | -28 | -18 |
| | L Lateral Occipital Cortex, inferior division | 5.46 | -46 | -64 | -6 |
| | L Inferior Temporal Gyrus, temporooccipital part | 5.23 | -54 | -50 | -14 |
| | L Superior Temporal Gyrus, posterior division | 4.55 | -56 | -34 | 4 |
| | L Inferior Temporal Gyrus, posterior division | 3.55 | -56 | -32 | -20 |
| **2,288** | L Middle Frontal Gyrus | 5.74 | -32 | 18 | 60 |
| **1,440** | L Lateral Occipital Cortex, inferior division | 5.35 | -28 | -88 | 2 |
| | L Occipital Pole | 4.81 | -36 | -92 | -4 |
| **992** | L Postcentral Gyrus | 4.68 | -56 | -10 | 26 |
| **840** | R Postcentral Gyrus | 4.75 | 64 | -6 | 26 |
| | R Precentral Gyrus | 4.58 | 58 | -4 | 24 |
| **592** | R Cingulate Gyrus, posterior division | 4.61 | 4 | -34 | 38 |
| **520** | L Inferior Frontal Gyrus, pars opercularis | 5.05 | -54 | 20 | 8 |

| | | | | | |
|---|---|---|---|---|---|
| **408** | L Insular Cortex | 5.64 | -32 | 22 | -4 |
| **328** | R Precentral Gyrus | 4.59 | 48 | 8 | 26 |
| **248** | R Insular Cortex | 6.14 | 34 | 22 | -2 |
| **248** | L Precentral Gyrus | 4.94 | -4 | -32 | 64 |
| **240** | R Middle Temporal Gyrus, posterior division | 3.62 | 60 | -10 | -30 |
| **240** | L Postcentral Gyrus | 5.71 | -50 | -22 | 58 |
| **224** | R Frontal Pole | 4.87 | 22 | 42 | -14 |
| **200** | R Parahippocampal Gyrus, posterior division | 5.05 | 34 | -34 | -14 |
| | R Temporal Fusiform Cortex, posterior division | 3.77 | 34 | -32 | -22 |
| **168** | R Hippocampus | 4.59 | 20 | -16 | -22 |
| **136** | R Inferior Frontal Gyrus, pars opercularis | 4.38 | 54 | 20 | 12 |
| **128** | R Paracingulate Gyrus | 3.87 | 10 | 46 | 20 |
| **64** | L Frontal Pole | 4.01 | -16 | 46 | -18 |
| **56** | R Cingulate Gyrus, posterior division | 4.36 | 6 | -44 | 16 |
| **48** | L Lateral Occipital Cortex, inferior division | 3.57 | -48 | -84 | -8 |
| **40** | R Insular Cortex | 3.68 | 36 | -6 | 10 |
| **32** | R Temporal Pole | 3.99 | 32 | 20 | -34 |
| **32** | R Lateral Occipital Cortex, inferior division | 3.43 | 32 | -88 | -18 |
| **32** | R Temporal Occipital Fusiform Cortex | 4.16 | 26 | -40 | -14 |
| **24** | R Thalamus | 3.84 | 12 | -26 | 12 |
| **24** | R Frontal Pole | 3.68 | 22 | 36 | 42 |
| **24** | R Precentral Gyrus | 4.02 | 4 | -28 | 66 |
| **24** | R Precentral Gyrus | 4.22 | 10 | -28 | 70 |
| **16** | R Hippocampus | 3.68 | 22 | -38 | 2 |
| **16** | R Frontal Pole | 3.42 | 6 | 66 | 4 |
| **16** | L Postcentral Gyrus | 3.47 | -6 | -42 | 66 |
| **8** | R Temporal Pole | 3.37 | 30 | 16 | -32 |
| **8** | L Frontal Medial Cortex | 3.30 | -8 | 32 | -24 |
| **8** | R Parahippocampal Gyrus, posterior division | 3.39 | 28 | -26 | -20 |
| **8** | L Hippocampus | 3.26 | -20 | -14 | -20 |
| **8** | R Frontal Orbital Cortex | 3.72 | 12 | 18 | -20 |
| **8** | R Hippocampus | 3.48 | 30 | -36 | -6 |
| **8** | L Thalamus | 3.30 | -18 | -24 | 0 |
| **8** | L Hippocampus | 3.30 | -12 | -40 | 4 |
| **8** | R Hippocampus | 3.50 | 16 | -38 | 4 |
| **8** | L Insular Cortex | 3.52 | -36 | -10 | 10 |
| **8** | L Cingulate Gyrus, anterior division | 3.24 | -10 | 36 | 14 |
| **8** | L Frontal Pole | 3.24 | -30 | 52 | 14 |
| **8** | R Cingulate Gyrus, anterior division | 3.31 | 6 | -8 | 30 |
| **8** | L Precentral Gyrus | 3.52 | -12 | -28 | 68 |

**Supplementary Table 4 | Brain regions where activation covaried with parametrically modulated threat Probability**

Whole-brain analysis of the parametric modulation of cue-evoked activation by trial-wise threat Probability, thresholded at FDR-corrected q < 0.005 (positive contrast only). $mm^3$, volume of the contiguous suprathreshold cluster; Label, anatomical region containing the local activation peak, identified using the probabilistic Harvard-Oxford cortical and subcortical atlases; L, left hemisphere; R, right hemisphere; t, peak voxel t-statistic; x, y, z, peak voxel coordinates in MNI space (mm). Clusters are ordered by descending volume, and peaks within each cluster by descending t-statistic.

| $mm^3$ | Label | t | x | y | z |
|---|---|---|---|---|---|
| **664,336** | R Postcentral Gyrus | 15.12 | 38 | -24 | 60 |
| | R Brain-Stem | 15.06 | 8 | -28 | -10 |
| | L Brain-Stem | 14.13 | -6 | -28 | -12 |
| | R Juxtapositional Lobule Cortex (formerly Supplementary Motor Cortex) | 13.46 | 8 | 4 | 56 |
| | R Cingulate Gyrus, anterior division | 13.32 | 4 | 0 | 46 |
| | R Periaqueductal Gray | 13.24 | 4 | -30 | -6 |
| | R Precentral Gyrus | 13.17 | 40 | -14 | 50 |
| | L Cingulate Gyrus, anterior division | 13.10 | -6 | 8 | 38 |
| | R Central Opercular Cortex | 13.06 | 50 | 4 | 4 |
| | R Occipital Fusiform Gyrus | 12.85 | 24 | -88 | -12 |
| | R Left Thalamus | 12.83 | 0 | -10 | 10 |
| | R Superior Frontal Gyrus | 12.52 | 16 | -4 | 70 |
| | L Precentral Gyrus | 12.40 | -58 | 4 | 8 |
| | R Paracingulate Gyrus | 12.29 | 6 | 8 | 46 |
| | R Lateral Occipital Cortex, inferior division | 12.09 | 38 | -80 | -12 |
| | R Occipital Pole | 11.98 | 16 | -92 | -12 |
| | L Juxtapositional Lobule Cortex (formerly Supplementary Motor Cortex) | 11.90 | -2 | 4 | 48 |
| | R Amygdala | 11.71 | 22 | -2 | -12 |
| | L Central Opercular Cortex | 11.68 | -54 | 2 | 4 |
| | R Frontal Operculum Cortex | 11.21 | 40 | 12 | 4 |
| | R Superior Parietal Lobule | 11.20 | 30 | -42 | 66 |
| | L Thalamus | 11.13 | -4 | -16 | -2 |
| | R Insular Cortex | 11.11 | 42 | 12 | 0 |
| | L Cingulate Gyrus, posterior division | 11.02 | -12 | -24 | 40 |
| | L Superior Frontal Gyrus | 10.74 | -8 | -6 | 72 |
| | R Lingual Gyrus | 10.70 | 12 | -84 | -14 |
| | R Parietal Operculum Cortex | 10.47 | 52 | -22 | 18 |
| | L Frontal Operculum Cortex | 10.32 | -44 | 12 | -2 |
| | R Thalamus | 10.28 | 10 | -2 | 12 |
| | L Occipital Fusiform Gyrus | 10.25 | -26 | -74 | -18 |
| | L Caudate | 9.76 | -12 | -2 | 14 |
| | R Frontal Orbital Cortex | 9.47 | 38 | 30 | 0 |
| | R Caudate | 9.32 | 8 | 6 | 0 |
| | R Supramarginal Gyrus, anterior division | 9.27 | 60 | -26 | 28 |
| | R Middle Temporal Gyrus, temporooccipital part | 9.18 | 54 | -60 | 0 |
| | L Occipital Pole | 9.10 | -26 | -98 | -2 |
| | L Bed Nucleus of the Stria Terminalis | 8.90 | -6 | 4 | -2 |
| | R Precuneus Cortex | 8.89 | 12 | -52 | 58 |

| | | | | | |
|---|---|---|---|---|---|
| | L Lateral Occipital Cortex, inferior division | 8.85 | -38 | -78 | -12 |
| | L Supramarginal Gyrus, anterior division | 8.70 | -62 | -28 | 20 |
| | L Superior Parietal Lobule | 8.49 | -20 | -52 | 62 |
| | L Insular Cortex | 8.47 | -34 | 2 | 10 |
| | R Pallidum | 8.40 | 18 | -2 | -4 |
| | L Precuneus Cortex | 8.37 | -8 | -54 | 58 |
| | R Putamen | 8.27 | 24 | 10 | -4 |
| | L Postcentral Gyrus | 8.26 | -66 | -22 | 26 |
| | L Amygdala | 8.25 | -24 | -4 | -12 |
| | L Temporal Occipital Fusiform Cortex | 8.04 | -32 | -62 | -14 |
| | L Parietal Operculum Cortex | 7.84 | -54 | -28 | 20 |
| | R Lateral Occipital Cortex, superior division | 7.57 | 30 | -72 | 30 |
| | R Temporal Occipital Fusiform Cortex | 7.48 | 46 | -58 | -18 |
| | L Pallidum | 7.44 | -20 | -10 | -4 |
| | R Inferior Temporal Gyrus, temporooccipital part | 7.22 | 46 | -40 | -22 |
| | L Lateral Occipital Cortex, superior division | 7.20 | -30 | -74 | 22 |
| | L Frontal Orbital Cortex | 7.08 | -28 | 18 | -14 |
| | L Frontal Pole | 7.00 | -32 | 46 | 30 |
| | R Frontal Pole | 6.94 | 34 | 44 | 26 |
| | L Intracalcarine Cortex | 6.86 | -18 | -66 | 8 |
| | R Supramarginal Gyrus, posterior division | 6.83 | 68 | -38 | 24 |
| | L Hippocampus | 6.57 | -26 | -18 | -12 |
| | L Inferior Temporal Gyrus, temporooccipital part | 6.30 | -50 | -50 | -26 |
| | L Putamen | 6.20 | -26 | 6 | -6 |
| | R Intracalcarine Cortex | 5.87 | 14 | -84 | 4 |
| | L Lingual Gyrus | 5.76 | -20 | -60 | 2 |
| | L Cuneal Cortex | 5.72 | -18 | -74 | 28 |
| | L Temporal Fusiform Cortex, posterior division | 5.31 | -40 | -38 | -22 |
| | R Hippocampus | 5.30 | 26 | -16 | -12 |
| | L Middle Temporal Gyrus, temporooccipital part | 5.18 | -50 | -56 | 4 |
| | L Planum Polare | 4.97 | -44 | -20 | -4 |
| | L Inferior Temporal Gyrus, posterior division | 4.95 | -46 | -42 | -20 |
| | R Superior Temporal Gyrus, posterior division | 4.80 | 52 | -18 | -6 |
| | R Inferior Frontal Gyrus, pars opercularis | 4.33 | 44 | 14 | 22 |
| | R Inferior Temporal Gyrus, posterior division | 3.80 | 48 | -30 | -22 |
| **176** | R Frontal Pole | 4.81 | 24 | 68 | -6 |
| **120** | R Temporal Fusiform Cortex, posterior division | 5.06 | 38 | -18 | -24 |
| **96** | L Frontal Orbital Cortex | 4.26 | -40 | 30 | -18 |
| **72** | R Cuneal Cortex | 4.79 | 8 | -76 | 24 |
| **56** | R Inferior Temporal Gyrus, posterior division | 4.04 | 56 | -20 | -34 |
| **48** | R Superior Frontal Gyrus | 3.71 | 6 | 54 | 34 |
| **40** | R Temporal Fusiform Cortex, posterior division | 4.20 | 42 | -12 | -40 |
| **32** | L Temporal Pole | 4.36 | -36 | 4 | -44 |
| **32** | L Temporal Pole | 3.83 | -32 | 4 | -38 |
| **32** | L Frontal Orbital Cortex | 3.73 | -28 | 32 | -22 |
| **24** | L Brain-Stem | 3.50 | -2 | -34 | -56 |
| **24** | L Supramarginal Gyrus, posterior division | 3.68 | -54 | -44 | 18 |
| **16** | R Temporal Pole | 3.52 | 38 | 8 | -44 |
| **16** | R Temporal Pole | 3.66 | 48 | 8 | -42 |
| **16** | L Temporal Fusiform Cortex, anterior division | 3.84 | -32 | -6 | -32 |

| | | | | | |
|---|---|---|---|---|---|
| **16** | R Temporal Pole | 3.82 | 40 | 18 | -26 |
| **16** | L Thalamus | 3.58 | -20 | -34 | -4 |
| **16** | L Inferior Frontal Gyrus, pars opercularis | 3.90 | -44 | 16 | 20 |
| **8** | L Brain-Stem | 3.46 | -6 | -36 | -60 |
| **8** | L Temporal Fusiform Cortex, anterior division | 3.57 | -34 | 0 | -40 |
| **8** | R Inferior Temporal Gyrus, posterior division | 3.65 | 46 | -20 | -34 |
| **8** | R Parahippocampal Gyrus, anterior division | 3.54 | 24 | -18 | -32 |
| **8** | L Inferior Temporal Gyrus, posterior division | 3.56 | -46 | -20 | -28 |
| **8** | R Frontal Orbital Cortex | 3.83 | 18 | 30 | -18 |
| **8** | R Frontal Orbital Cortex | 3.69 | 18 | 22 | -16 |
| **8** | L Frontal Pole | 3.54 | -22 | 70 | -4 |
| **8** | R Cuneal Cortex | 3.46 | 8 | -82 | 24 |

**Supplementary Table 5 | Brain regions where activation covaried with parametrically modulated Risk**

Whole-brain analysis of the parametric modulation of cue-evoked activation by trial-wise Risk, thresholded at FDR-corrected q < 0.005 (positive contrast only). $mm^3$, volume of the contiguous suprathreshold cluster; Label, anatomical region containing the local activation peak, identified using the probabilistic Harvard-Oxford cortical and subcortical atlases; L, left hemisphere; R, right hemisphere; t, peak voxel t-statistic; x, y, z, peak voxel coordinates in MNI space (mm). Clusters are ordered by descending volume, and peaks within each cluster by descending t-statistic.

| $mm^3$ | Label | t | x | y | z |
|---|---|---|---|---|---|
| **28,024** | R Supramarginal Gyrus, posterior division | 9.05 | 48 | -38 | 48 |
| | R Lateral Occipital Cortex, superior division | 8.51 | 34 | -70 | 36 |
| | R Angular Gyrus | 7.31 | 38 | -54 | 48 |
| | R Precuneus Cortex | 5.66 | 6 | -68 | 44 |
| **16,376** | L Supramarginal Gyrus, posterior division | 8.74 | -46 | -42 | 46 |
| | L Supramarginal Gyrus, anterior division | 8.12 | -46 | -38 | 42 |
| | L Lateral Occipital Cortex, superior division | 8.10 | -28 | -68 | 30 |
| | L Precuneus Cortex | 6.49 | -4 | -72 | 48 |
| **13,672** | R Occipital Pole | 10.18 | 34 | -94 | 4 |
| | R Inferior Temporal Gyrus, temporooccipital part | 6.95 | 50 | -60 | -10 |
| | R Occipital Fusiform Gyrus | 5.54 | 36 | -70 | -14 |
| | R Lateral Occipital Cortex, inferior division | 5.16 | 50 | -68 | -14 |
| | R Middle Temporal Gyrus, posterior division | 4.22 | 62 | -34 | -12 |
| **11,232** | L Paracingulate Gyrus | 9.52 | -6 | 16 | 48 |
| | R Paracingulate Gyrus | 7.96 | 8 | 20 | 46 |
| | L Cingulate Gyrus, anterior division | 6.75 | -10 | 24 | 28 |
| | L Superior Frontal Gyrus | 4.74 | -16 | 14 | 62 |
| **10,424** | R Middle Frontal Gyrus | 6.04 | 32 | 16 | 54 |
| | R Precentral Gyrus | 5.66 | 48 | 8 | 26 |
| | R Frontal Pole | 5.55 | 48 | 36 | 26 |
| | R Superior Frontal Gyrus | 4.85 | 22 | 22 | 54 |
| **8,112** | L Occipital Pole | 7.95 | -32 | -92 | -2 |
| | L Lateral Occipital Cortex, inferior division | 5.19 | -42 | -70 | -8 |
| | L Inferior Temporal Gyrus, temporooccipital part | 4.76 | -48 | -58 | -8 |
| **4,552** | L Middle Frontal Gyrus | 5.83 | -42 | 30 | 26 |
| | L Precentral Gyrus | 5.26 | -40 | 2 | 34 |
| | L Inferior Frontal Gyrus, pars opercularis | 3.70 | -40 | 16 | 26 |
| **3,480** | L Insular Cortex | 8.96 | -30 | 22 | -4 |
| | L Frontal Orbital Cortex | 8.11 | -32 | 26 | 0 |
| | L Frontal Operculum Cortex | 5.74 | -40 | 16 | 6 |
| **3,112** | L Middle Frontal Gyrus | 6.31 | -26 | -4 | 52 |
| | L Superior Frontal Gyrus | 6.15 | -26 | 4 | 56 |
| **2,640** | R Insular Cortex | 7.96 | 34 | 24 | -2 |
| | R Frontal Orbital Cortex | 4.50 | 42 | 28 | -4 |
| | R Inferior Frontal Gyrus, pars triangularis | 4.12 | 52 | 22 | 12 |
| | R Frontal Operculum Cortex | 3.60 | 42 | 22 | 6 |
| **2,408** | R Frontal Pole | 5.73 | 42 | 48 | -18 |
| **568** | R Precuneus Cortex | 4.67 | 14 | -64 | 26 |
| **344** | R Putamen | 4.31 | 30 | 6 | -6 |
| **344** | L Frontal Pole | 3.92 | -42 | 48 | -4 |
| **208** | R Thalamus | 4.67 | 16 | -36 | 4 |
| | R Hippocampus | 4.35 | 24 | -34 | -6 |

| | | | | | |
|---|---|---|---|---|---|
| **208** | L Caudate | 4.38 | -8 | 10 | 2 |
| **176** | R Paracingulate Gyrus | 4.09 | 8 | 44 | 16 |
| **152** | R Thalamus | 3.95 | 10 | -6 | 8 |
| **128** | R Thalamus | 4.79 | 4 | -16 | 4 |
| **88** | R Cingulate Gyrus, anterior division | 3.92 | 4 | 2 | 28 |
| **80** | L Frontal Pole | 3.97 | -38 | 58 | 8 |
| **72** | R Caudate | 4.42 | 10 | 10 | 10 |
| **56** | L Thalamus | 4.45 | -10 | -24 | 12 |
| **56** | L Cingulate Gyrus, anterior division | 3.72 | -6 | 40 | 16 |
| **56** | R Precuneus Cortex | 4.01 | 4 | -66 | 34 |
| **48** | R Inferior Frontal Gyrus, pars triangularis | 3.82 | 50 | 20 | 4 |
| **40** | L Frontal Pole | 3.58 | -30 | 52 | 16 |
| **32** | R Lateral Occipital Cortex, inferior division | 3.57 | 42 | -84 | -18 |
| **32** | L Thalamus | 3.66 | -6 | -12 | 6 |
| **32** | R Frontal Pole | 3.80 | 32 | 52 | 6 |
| **24** | R Frontal Pole | 3.73 | 16 | 50 | -20 |
| **24** | R Frontal Pole | 3.67 | 30 | 48 | -12 |
| **24** | L Inferior Frontal Gyrus, pars opercularis | 3.54 | -48 | 16 | 2 |
| **24** | R Precuneus Cortex | 3.61 | 14 | -64 | 38 |
| **16** | R Frontal Pole | 3.55 | 10 | 40 | -22 |
| **16** | L Frontal Pole | 3.65 | -16 | 44 | -20 |
| **16** | L Lateral Occipital Cortex, inferior division | 3.49 | -50 | -80 | -6 |
| **16** | L Pallidum | 3.78 | -14 | 4 | -6 |
| **16** | L Hippocampus | 3.56 | -16 | -40 | 0 |
| **16** | R Caudate | 3.84 | 10 | 12 | 2 |
| **16** | R Putamen | 4.17 | 32 | -4 | 8 |
| **16** | L Central Opercular Cortex | 3.87 | -36 | -18 | 20 |
| **16** | R Paracingulate Gyrus | 3.62 | 6 | 40 | 20 |
| **16** | L Cingulate Gyrus, anterior division | 4.02 | -4 | 6 | 28 |
| **16** | R Cingulate Gyrus, anterior division | 3.73 | 6 | -6 | 32 |
| **16** | R Superior Frontal Gyrus | 3.46 | 22 | 14 | 64 |
| **8** | R Frontal Medial Cortex | 3.49 | 6 | 42 | -22 |
| **8** | R Middle Temporal Gyrus, posterior division | 3.42 | 62 | -16 | -16 |
| **8** | L Temporal Occipital Fusiform Cortex | 3.55 | -34 | -52 | -14 |
| **8** | R Thalamus | 3.75 | 12 | -26 | 12 |
| **8** | L Caudate | 3.57 | -16 | 4 | 18 |

**Supplementary Table 6 | Brain regions where activation covaried with parametrically modulated Ambiguity**

Whole-brain analysis of the parametric modulation of cue-evoked activation by trial-wise Ambiguity, thresholded at FDR-corrected $q < 0.005$ (positive contrast only). $mm^3$, volume of the contiguous suprathreshold cluster; Label, anatomical region containing the local activation peak, identified using the probabilistic Harvard-Oxford cortical and subcortical atlases; L, left hemisphere; R, right hemisphere; t, peak voxel t-statistic; x, y, z, peak voxel coordinates in MNI space (mm). Clusters are ordered by descending volume, and peaks within each cluster by descending t-statistic.

| $mm^3$ | Label | t | x | y | z |
|---|---|---|---|---|---|
| **19,000** | L Occipital Pole | 7.14 | -14 | -92 | 24 |
| | L Lateral Occipital Cortex, superior division | 6.49 | -28 | -72 | 36 |
| | R Occipital Pole | 6.48 | 6 | -92 | 18 |
| | R Supracalcarine Cortex | 6.06 | 0 | -88 | 10 |
| | R Cuneal Cortex | 5.76 | 4 | -88 | 18 |
| | R Lateral Occipital Cortex, superior division | 5.31 | 36 | -78 | 34 |
| | R Intracalcarine Cortex | 4.95 | 2 | -72 | 8 |
| | L Intracalcarine Cortex | 4.73 | -2 | -82 | 2 |
| | L Lingual Gyrus | 4.51 | -4 | -76 | -4 |
| | R Lingual Gyrus | 3.99 | 10 | -76 | 0 |
| **8,216** | L Superior Frontal Gyrus | 5.51 | -18 | 12 | 66 |
| | R Superior Frontal Gyrus | 5.41 | 10 | 22 | 60 |
| | L Paracingulate Gyrus | 4.90 | -4 | 26 | 44 |
| | L Middle Frontal Gyrus | 4.90 | -36 | 4 | 58 |
| **4,536** | R Lateral Occipital Cortex, superior division | 4.82 | 38 | -60 | 44 |
| | R Angular Gyrus | 4.76 | 50 | -48 | 42 |
| | R Supramarginal Gyrus, posterior division | 4.41 | 56 | -46 | 36 |
| | R Superior Parietal Lobule | 3.82 | 34 | -46 | 42 |
| **4,056** | L Angular Gyrus | 5.55 | -48 | -54 | 52 |
| | L Supramarginal Gyrus, posterior division | 5.09 | -52 | -48 | 40 |
| | L Lateral Occipital Cortex, superior division | 4.33 | -42 | -60 | 54 |
| | L Superior Parietal Lobule | 3.57 | -34 | -52 | 44 |
| **3,288** | L Precuneus Cortex | 5.00 | -4 | -70 | 50 |
| | R Lateral Occipital Cortex, superior division | 4.67 | 10 | -74 | 58 |
| | L Lateral Occipital Cortex, superior division | 4.63 | -10 | -74 | 56 |
| | R Precuneus Cortex | 3.99 | 8 | -64 | 62 |
| **2,824** | L Inferior Frontal Gyrus, pars triangularis | 5.52 | -50 | 22 | 0 |
| | L Frontal Pole | 5.33 | -42 | 42 | -12 |
| | L Frontal Orbital Cortex | 5.01 | -36 | 20 | -12 |
| | L Inferior Frontal Gyrus, pars opercularis | 4.93 | -56 | 22 | 16 |
| | L Insular Cortex | 4.68 | -30 | 18 | -8 |
| **2,744** | L Middle Frontal Gyrus | 5.77 | -40 | 24 | 46 |
| **1,160** | R Lateral Occipital Cortex, inferior division | 4.90 | 48 | -70 | -8 |
| | R Middle Temporal Gyrus, temporooccipital part | 4.27 | 62 | -52 | -6 |
| | R Inferior Temporal Gyrus, temporooccipital part | 4.18 | 54 | -58 | -6 |
| | R Temporal Occipital Fusiform Cortex | 3.99 | 42 | -54 | -10 |
| **1,088** | R Frontal Orbital Cortex | 4.81 | 48 | 28 | -6 |
| | R Frontal Pole | 4.33 | 52 | 44 | -4 |
| | R Inferior Frontal Gyrus, pars triangularis | 4.07 | 56 | 24 | -2 |
| **888** | L Frontal Pole | 4.69 | -28 | 52 | 24 |

| | | | | | |
|---|---|---|---|---|---|
| **488** | L Lateral Occipital Cortex, inferior division | 4.48 | -48 | -68 | -14 |
| **480** | R Middle Frontal Gyrus | 5.27 | 38 | 10 | 54 |
| **392** | R Middle Temporal Gyrus, posterior division | 4.67 | 64 | -24 | -6 |
| **304** | R Precentral Gyrus | 4.53 | 46 | 10 | 34 |
| | R Middle Frontal Gyrus | 4.41 | 42 | 8 | 34 |
| **160** | L Lateral Occipital Cortex, inferior division | 4.80 | -42 | -84 | 4 |
| **152** | R Lingual Gyrus | 4.35 | 22 | -66 | -6 |
| | R Occipital Fusiform Gyrus | 3.98 | 26 | -64 | -6 |
| **152** | L Lateral Occipital Cortex, inferior division | 3.95 | -56 | -66 | -6 |
| **136** | L Caudate | 4.28 | -8 | 12 | 0 |
| **120** | L Middle Temporal Gyrus, posterior division | 4.18 | -64 | -32 | -6 |
| **120** | L Caudate | 4.73 | -10 | 14 | 10 |
| **112** | R Middle Temporal Gyrus, posterior division | 4.09 | 66 | -36 | -10 |
| **112** | L Lateral Occipital Cortex, superior division | 4.54 | -36 | -84 | 12 |
| **96** | R Lateral Occipital Cortex, inferior division | 3.93 | 42 | -74 | 10 |
| **88** | L Middle Temporal Gyrus, posterior division | 4.11 | -68 | -24 | -8 |
| **80** | R Inferior Temporal Gyrus, temporooccipital part | 3.84 | 48 | -56 | -18 |
| **80** | L Middle Temporal Gyrus, posterior division | 3.74 | -60 | -28 | -12 |
| **80** | L Thalamus | 4.15 | -6 | 0 | 6 |
| **80** | R Lateral Occipital Cortex, superior division | 4.23 | 30 | -62 | 62 |
| **72** | R Lateral Occipital Cortex, superior division | 4.11 | 42 | -70 | 38 |
| **64** | L Inferior Temporal Gyrus, posterior division | 4.26 | -60 | -26 | -22 |
| **64** | R Inferior Temporal Gyrus, temporooccipital part | 4.10 | 56 | -50 | -16 |
| **64** | L Inferior Temporal Gyrus, temporooccipital part | 3.62 | -48 | -54 | -12 |
| **64** | R Caudate | 4.68 | 10 | 12 | -2 |
| **64** | R Middle Frontal Gyrus | 3.67 | 52 | 14 | 40 |
| **56** | L Middle Temporal Gyrus, anterior division | 4.04 | -58 | -2 | -32 |
| **56** | L Frontal Pole | 4.11 | -28 | 56 | -2 |
| **56** | L Frontal Pole | 4.22 | -36 | 64 | -2 |
| **56** | L Middle Frontal Gyrus | 3.99 | -30 | 32 | 34 |
| **56** | L Precuneus Cortex | 4.25 | -8 | -64 | 40 |
| **56** | L Juxtapositional Lobule Cortex (formerly Supplementary Motor Cortex) | 4.01 | -2 | 4 | 62 |
| **48** | L Occipital Fusiform Gyrus | 3.64 | -38 | -66 | -20 |
| **48** | R Temporal Occipital Fusiform Cortex | 4.01 | 32 | -46 | -8 |
| **48** | R Inferior Frontal Gyrus, pars opercularis | 4.17 | 50 | 20 | 12 |
| **48** | L Middle Frontal Gyrus | 4.06 | -46 | 28 | 32 |
| **40** | R Frontal Orbital Cortex | 3.75 | 36 | 24 | -8 |
| **40** | R Intracalcarine Cortex | 3.97 | 8 | -78 | 12 |
| **40** | L Thalamus | 3.73 | -6 | -22 | 16 |
| **40** | L Caudate | 3.74 | -14 | -4 | 20 |
| **40** | R Cuneal Cortex | 3.79 | 2 | -84 | 34 |
| **32** | L Middle Temporal Gyrus, posterior division | 3.77 | -56 | -38 | -14 |
| **32** | R Lateral Occipital Cortex, superior division | 3.95 | 38 | -78 | 12 |
| **32** | L Inferior Frontal Gyrus, pars opercularis | 3.72 | -54 | 18 | 28 |
| **32** | L Precentral Gyrus | 4.54 | -32 | -18 | 44 |
| **32** | R Superior Frontal Gyrus | 3.74 | 24 | 16 | 54 |
| **32** | R Superior Frontal Gyrus | 3.94 | 22 | 26 | 56 |
| **24** | R Middle Temporal Gyrus, anterior division | 3.72 | 54 | 0 | -36 |
| **24** | L Temporal Fusiform Cortex, posterior division | 4.03 | -34 | -26 | -20 |
| **24** | R Parahippocampal Gyrus, posterior division | 3.85 | 22 | -36 | -18 |
| **24** | L Frontal Pole | 4.12 | -4 | 66 | -16 |
| **24** | L Occipital Fusiform Gyrus | 3.85 | -24 | -70 | -12 |

| | | | | | |
|---|---|---|---|---|---|
| **24** | L Inferior Temporal Gyrus, temporooccipital part | 3.75 | -54 | -56 | -14 |
| **24** | R Temporal Occipital Fusiform Cortex | 3.90 | 28 | -42 | -12 |
| **24** | L Temporal Occipital Fusiform Cortex | 4.06 | -30 | -60 | -6 |
| **24** | L Lateral Occipital Cortex, superior division | 3.44 | -18 | -72 | 58 |
| **16** | R Brain-Stem | 4.08 | 4 | -36 | -38 |
| **16** | L Inferior Temporal Gyrus, posterior division | 4.02 | -56 | -32 | -28 |
| **16** | L Inferior Temporal Gyrus, posterior division | 3.55 | -64 | -40 | -22 |
| **16** | R Inferior Temporal Gyrus, temporooccipital part | 3.64 | 58 | -58 | -16 |
| **16** | L Temporal Occipital Fusiform Cortex | 3.46 | -22 | -56 | -12 |
| **16** | L Middle Temporal Gyrus, posterior division | 3.53 | -64 | -18 | -8 |
| **16** | L Middle Temporal Gyrus, posterior division | 3.78 | -66 | -44 | -6 |
| **16** | L Middle Temporal Gyrus, posterior division | 3.56 | -58 | -42 | -2 |
| **16** | R Caudate | 3.76 | 10 | 18 | 2 |
| **16** | R Caudate | 3.74 | 12 | 12 | 6 |
| **16** | R Caudate | 3.58 | 12 | 8 | 12 |
| **16** | R Caudate | 3.49 | 12 | 4 | 20 |
| **16** | R Inferior Frontal Gyrus, pars triangularis | 3.49 | 58 | 26 | 18 |
| **16** | L Paracingulate Gyrus | 3.51 | -8 | 46 | 24 |
| **16** | R Cuneal Cortex | 3.62 | 2 | -80 | 28 |
| **16** | R Cingulate Gyrus, anterior division | 3.73 | 6 | 22 | 30 |
| **16** | L Middle Frontal Gyrus | 3.53 | -52 | 26 | 32 |
| **16** | L Paracingulate Gyrus | 3.57 | -10 | 30 | 30 |
| **16** | L Middle Frontal Gyrus | 3.79 | -34 | 8 | 34 |
| **16** | L Frontal Pole | 3.47 | -24 | 40 | 36 |
| **16** | L Frontal Pole | 3.67 | -14 | 50 | 34 |
| **16** | R Precuneus Cortex | 3.49 | 12 | -70 | 46 |
| **16** | L Superior Frontal Gyrus | 3.72 | -14 | 32 | 50 |
| **16** | R Lateral Occipital Cortex, superior division | 3.50 | 24 | -74 | 56 |
| **8** | R Brain-Stem | 3.55 | 14 | -30 | -44 |
| **8** | R Brain-Stem | 3.59 | 14 | -30 | -38 |
| **8** | R Middle Temporal Gyrus, anterior division | 3.57 | 58 | -6 | -32 |
| **8** | R Brain-Stem | 3.43 | 2 | -22 | -28 |
| **8** | R Inferior Temporal Gyrus, posterior division | 3.44 | 60 | -16 | -28 |
| **8** | L Temporal Fusiform Cortex, posterior division | 3.82 | -40 | -40 | -26 |
| **8** | R Inferior Temporal Gyrus, posterior division | 3.80 | 56 | -26 | -26 |
| **8** | L Temporal Fusiform Cortex, posterior division | 3.74 | -34 | -38 | -24 |
| **8** | R Brain-Stem | 3.91 | 16 | -22 | -24 |
| **8** | R Frontal Orbital Cortex | 3.45 | 30 | 22 | -24 |
| **8** | R Frontal Orbital Cortex | 3.92 | 12 | 22 | -22 |
| **8** | R Temporal Occipital Fusiform Cortex | 3.46 | 38 | -58 | -18 |
| **8** | L Frontal Orbital Cortex | 3.43 | -46 | 34 | -18 |
| **8** | L Frontal Pole | 3.46 | -44 | 42 | -18 |
| **8** | L Frontal Pole | 3.52 | -48 | 42 | -16 |
| **8** | R Temporal Occipital Fusiform Cortex | 3.41 | 30 | -52 | -14 |
| **8** | R Inferior Temporal Gyrus, temporooccipital part | 3.58 | 48 | -48 | -14 |
| **8** | L Middle Temporal Gyrus, posterior division | 3.42 | -58 | -34 | -14 |
| **8** | L Frontal Pole | 3.41 | -10 | 64 | -14 |
| **8** | R Subcallosal Cortex | 3.48 | 12 | 16 | -12 |
| **8** | L Middle Temporal Gyrus, temporooccipital part | 3.63 | -64 | -50 | -8 |
| **8** | L Parahippocampal Gyrus, posterior division | 3.53 | -14 | -38 | -6 |
| **8** | R Middle Temporal Gyrus, temporooccipital part | 3.49 | 52 | -52 | -2 |
| **8** | R Lingual Gyrus | 3.45 | 12 | -44 | -2 |
| **8** | L Middle Temporal Gyrus, posterior division | 3.44 | -66 | -36 | -2 |

| | | | | | |
|---|---|---|---|---|---|
| **8** | R Hippocampus | 3.75 | 20 | -40 | 2 |
| **8** | L Thalamus | 3.45 | -2 | -4 | 4 |
| **8** | L Thalamus | 3.64 | -16 | -36 | 6 |
| **8** | R Caudate | 3.72 | 12 | 8 | 6 |
| **8** | L Frontal Pole | 3.47 | -46 | 50 | 6 |
| **8** | L Planum Temporale | 3.55 | -42 | -36 | 8 |
| **8** | L Putamen | 3.48 | -24 | 2 | 8 |
| **8** | L Frontal Pole | 3.44 | -44 | 52 | 8 |
| **8** | L Thalamus | 3.42 | -8 | -16 | 12 |
| **8** | L Caudate | 3.48 | -10 | 4 | 16 |
| **8** | R Middle Frontal Gyrus | 3.44 | 52 | 28 | 24 |
| **8** | R Lateral Occipital Cortex, superior division | 3.41 | 52 | -60 | 26 |
| **8** | R Precentral Gyrus | 3.52 | 44 | 6 | 26 |
| **8** | R Lateral Occipital Cortex, superior division | 3.45 | 38 | -68 | 28 |
| **8** | L Paracingulate Gyrus | 3.45 | -8 | 38 | 30 |
| **8** | R Paracingulate Gyrus | 3.66 | 10 | 36 | 32 |
| **8** | L Frontal Pole | 3.52 | -14 | 60 | 34 |
| **8** | R Lateral Occipital Cortex, superior division | 3.50 | 48 | -68 | 36 |
| **8** | L Superior Parietal Lobule | 3.54 | -32 | -52 | 40 |
| **8** | R Cingulate Gyrus, posterior division | 3.65 | 4 | -20 | 40 |
| **8** | R Precuneus Cortex | 3.43 | 6 | -68 | 44 |
| **8** | R Lateral Occipital Cortex, superior division | 3.54 | 26 | -78 | 46 |
| **8** | R Middle Frontal Gyrus | 3.59 | 36 | 14 | 46 |
| **8** | L Middle Frontal Gyrus | 3.59 | -30 | 18 | 46 |
| **8** | L Precentral Gyrus | 3.55 | -48 | -4 | 48 |
| **8** | R Middle Frontal Gyrus | 3.79 | 40 | 16 | 48 |
| **8** | R Frontal Pole | 3.53 | 12 | 42 | 50 |
| **8** | L Lateral Occipital Cortex, superior division | 3.64 | -20 | -66 | 54 |
| **8** | L Precuneus Cortex | 3.45 | -4 | -58 | 66 |
| **8** | L Superior Frontal Gyrus | 3.62 | -8 | 2 | 68 |

#### Supplementary Table 7 | Brain regions showing greater activation for 50% threat vs. Safe

Whole-brain analysis contrasting cue-evoked activation during 50% (50% threat probability) versus Safe (0% threat probability) trials ( 50% Threat > Safe), thresholded at FDR-corrected q < 0.005 (positive contrast only). $mm^3$, volume of the contiguous suprathreshold cluster; Label, anatomical region containing the local activation peak, identified using the probabilistic Harvard-Oxford cortical and subcortical atlases; L, left hemisphere; R, right hemisphere; t, peak voxel t-statistic; x, y, z, peak voxel coordinates in MNI space (mm). Clusters are ordered by descending volume, and peaks within each cluster by descending t-statistic.

| $mm^3$ | Label | t | x | y | z |
|---|---|---|---|---|---|
| **110,288** | R Lateral Occipital Cortex, superior division | 11.83 | 32 | -66 | 30 |
| | R Lateral Occipital Cortex, inferior division | 10.56 | 50 | -70 | -12 |
| | R Occipital Pole | 10.48 | 36 | -90 | 8 |
| | R Superior Parietal Lobule | 10.25 | 32 | -46 | 44 |
| | R Supramarginal Gyrus, posterior division | 9.66 | 46 | -40 | 48 |
| | R Inferior Temporal Gyrus, temporooccipital part | 9.60 | 50 | -58 | -12 |
| | L Paracingulate Gyrus | 9.30 | -6 | 12 | 50 |
| | R Occipital Fusiform Gyrus | 8.95 | 36 | -78 | -14 |
| | L Middle Frontal Gyrus | 8.82 | -26 | -4 | 52 |
| | L Juxtapositional Lobule Cortex (formerly Supplementary Motor Cortex) | 8.70 | -4 | 8 | 54 |
| | R Supramarginal Gyrus, anterior division | 8.36 | 52 | -28 | 48 |
| | R Paracingulate Gyrus | 8.28 | 8 | 20 | 36 |
| | R Angular Gyrus | 8.01 | 36 | -54 | 48 |
| | R Superior Frontal Gyrus | 7.74 | 4 | 12 | 54 |
| | R Middle Frontal Gyrus | 7.25 | 28 | 8 | 56 |
| | R Postcentral Gyrus | 6.97 | 48 | -34 | 58 |
| | L Cingulate Gyrus, anterior division | 6.96 | -8 | 22 | 30 |
| | L Precentral Gyrus | 6.95 | -52 | 6 | 38 |
| | R Cingulate Gyrus, anterior division | 6.31 | 4 | 16 | 22 |
| | L Superior Frontal Gyrus | 6.15 | -14 | 12 | 62 |
| | R Precentral Gyrus | 6.07 | 32 | -26 | 54 |
| | R Precuneus Cortex | 5.99 | 10 | -72 | 48 |
| | R Temporal Occipital Fusiform Cortex | 5.67 | 28 | -54 | -14 |
| | R Middle Temporal Gyrus, temporooccipital part | 4.57 | 62 | -46 | -10 |
| | R Juxtapositional Lobule Cortex (formerly Supplementary Motor Cortex) | 4.40 | 10 | 4 | 44 |
| **65,480** | L Lateral Occipital Cortex, inferior division | 10.57 | -40 | -72 | -6 |
| | L Lateral Occipital Cortex, superior division | 10.19 | -22 | -66 | 48 |
| | L Occipital Pole | 9.28 | -32 | -92 | 10 |
| | L Superior Parietal Lobule | 9.15 | -26 | -56 | 48 |
| | L Supramarginal Gyrus, posterior division | 8.82 | -44 | -42 | 46 |
| | L Supramarginal Gyrus, anterior division | 8.22 | -46 | -38 | 42 |
| | L Occipital Fusiform Gyrus | 6.56 | -32 | -70 | -16 |
| | L Inferior Temporal Gyrus, temporooccipital part | 6.53 | -48 | -62 | -16 |
| | L Precuneus Cortex | 6.29 | -4 | -68 | 50 |
| | L Temporal Occipital Fusiform Cortex | 5.89 | -34 | -58 | -14 |
| | L Temporal Fusiform Cortex, posterior division | 4.84 | -32 | -44 | -20 |
| **5,760** | L Insular Cortex | 8.10 | -30 | 22 | -6 |
| | L Frontal Operculum Cortex | 7.08 | -40 | 22 | 0 |

| | | | | | |
|---|---|---|---|---|---|
| | L Inferior Frontal Gyrus, pars opercularis | 5.72 | -46 | 18 | 10 |
| **5,720** | R Precentral Gyrus | 7.39 | 46 | 8 | 34 |
| | R Inferior Frontal Gyrus, pars opercularis | 6.79 | 52 | 14 | 30 |
| | R Middle Frontal Gyrus | 5.42 | 48 | 34 | 30 |
| | R Frontal Pole | 3.97 | 50 | 38 | 22 |
| **4,568** | R Frontal Orbital Cortex | 7.94 | 36 | 26 | -6 |
| | R Insular Cortex | 7.77 | 36 | 24 | 0 |
| | R Frontal Operculum Cortex | 6.66 | 38 | 20 | 4 |
| | R Inferior Frontal Gyrus, pars triangularis | 5.74 | 46 | 24 | 6 |
| **3,048** | R Caudate | 5.87 | 8 | 6 | 0 |
| | R Putamen | 5.80 | 22 | 8 | 4 |
| | R Pallidum | 4.78 | 16 | 6 | -4 |
| **1,856** | L Thalamus | 5.55 | -10 | -12 | 8 |
| | L Brain-Stem | 4.94 | -6 | -30 | -20 |
| | R Brain-Stem | 4.04 | 0 | -20 | -24 |
| **1,368** | R Brain-Stem | 5.32 | 8 | -30 | -24 |
| | R Thalamus | 4.75 | 12 | -14 | 2 |
| **752** | L Caudate | 5.94 | -8 | 6 | 6 |
| **464** | L Frontal Pole | 4.74 | -30 | 54 | 22 |
| **456** | R Insular Cortex | 5.59 | 36 | -20 | 8 |
| | R Central Opercular Cortex | 4.29 | 46 | -18 | 20 |
| | R Parietal Operculum Cortex | 3.88 | 42 | -22 | 16 |
| **320** | L Middle Frontal Gyrus | 4.23 | -44 | 34 | 34 |
| **304** | R Thalamus | 5.17 | 12 | -24 | 14 |
| **208** | R Brain-Stem | 5.41 | 2 | -32 | -42 |
| **160** | L Putamen | 4.13 | -22 | 8 | -2 |
| **112** | R Middle Frontal Gyrus | 4.37 | 42 | 34 | 20 |
| **104** | L Frontal Pole | 5.44 | -48 | 52 | -2 |
| **96** | L Brain-Stem | 4.67 | -6 | -48 | -58 |
| **96** | L Thalamus | 4.29 | -12 | -16 | 16 |
| **72** | L Middle Frontal Gyrus | 4.33 | -46 | 20 | 28 |
| **72** | L Precentral Gyrus | 4.00 | -56 | -4 | 46 |
| **64** | L Superior Frontal Gyrus | 4.24 | -6 | 14 | 70 |
| **56** | R Temporal Fusiform Cortex, posterior division | 4.21 | 28 | -36 | -20 |
| **56** | R Lingual Gyrus | 4.42 | 6 | -86 | -10 |
| **48** | R Middle Temporal Gyrus, posterior division | 3.99 | 66 | -24 | -4 |
| **48** | L Heschls Gyrus (includes H1 and H2) | 4.08 | -42 | -28 | 14 |
| **40** | R Brain-Stem | 4.14 | 10 | -28 | -38 |
| **40** | L Precentral Gyrus | 3.82 | -52 | 6 | 2 |
| **40** | L Planum Temporale | 3.84 | -54 | -24 | 8 |
| **40** | L Cingulate Gyrus, anterior division | 3.89 | -8 | 38 | 18 |
| | L Paracingulate Gyrus | 3.88 | -4 | 40 | 20 |
| **32** | L Inferior Temporal Gyrus, anterior division | 4.37 | -46 | -4 | -42 |
| **32** | R Frontal Pole | 3.99 | 26 | 48 | -14 |
| **32** | L Pallidum | 3.84 | -16 | 2 | -6 |
| **32** | L Precentral Gyrus | 4.03 | -42 | -12 | 46 |
| **24** | L Brain-Stem | 3.85 | -6 | -42 | -52 |
| **24** | L Brain-Stem | 4.23 | -10 | -24 | -34 |
| **24** | R Frontal Pole | 4.38 | 44 | 52 | -16 |
| **24** | R Frontal Pole | 3.84 | 40 | 62 | -6 |
| **24** | R Hippocampus | 3.69 | 22 | -36 | 4 |
| **24** | L Insular Cortex | 4.15 | -38 | -18 | 14 |

| | | | | | |
|---|---|---|---|---|---|
| **24** | L Superior Frontal Gyrus | 4.00 | -8 | -6 | 68 |
| **16** | L Brain-Stem | 4.00 | -6 | -38 | -48 |
| **16** | L Brain-Stem | 3.75 | -4 | -34 | -24 |
| **16** | R Brain-Stem | 3.81 | 8 | -38 | -14 |
| **16** | R Hippocampus | 3.58 | 22 | -32 | -6 |
| **16** | R Putamen | 3.94 | 32 | -2 | 8 |
| **16** | R Thalamus | 3.86 | 18 | -32 | 10 |
| **16** | L Caudate | 3.86 | -14 | 12 | 10 |
| **16** | L Frontal Pole | 3.73 | -26 | 48 | 14 |
| **16** | L Central Opercular Cortex | 3.67 | -44 | -22 | 16 |
| **16** | R Caudate | 3.69 | 16 | 8 | 16 |
| **16** | R Frontal Pole | 3.68 | 30 | 48 | 20 |
| **16** | R Cingulate Gyrus, posterior division | 4.00 | 8 | -26 | 28 |
| **16** | L Cingulate Gyrus, posterior division | 4.20 | -6 | -24 | 30 |
| **16** | R Angular Gyrus | 3.69 | 58 | -48 | 34 |
| **16** | L Cingulate Gyrus, anterior division | 4.09 | -4 | -2 | 34 |
| **16** | R Superior Frontal Gyrus | 3.82 | 4 | 36 | 54 |
| **16** | R Superior Frontal Gyrus | 3.78 | 18 | -2 | 66 |
| **16** | L Postcentral Gyrus | 3.72 | -14 | -42 | 72 |
| **8** | R Brain-Stem | 3.70 | 8 | -48 | -58 |
| **8** | L Brain-Stem | 3.64 | -8 | -44 | -48 |
| **8** | R Brain-Stem | 3.72 | 2 | -28 | -32 |
| **8** | R Inferior Temporal Gyrus, posterior division | 3.63 | 48 | -20 | -32 |
| **8** | R Temporal Occipital Fusiform Cortex | 3.56 | 36 | -40 | -20 |
| **8** | L Brain-Stem | 3.70 | -10 | -22 | -20 |
| **8** | R Frontal Pole | 3.60 | 30 | 54 | -18 |
| **8** | R Brain-Stem | 3.55 | 12 | -26 | -16 |
| **8** | R Frontal Pole | 3.57 | 26 | 54 | -14 |
| **8** | R Middle Temporal Gyrus, posterior division | 3.63 | 50 | -30 | -8 |
| **8** | R Frontal Pole | 3.67 | 44 | 58 | -6 |
| **8** | R Insular Cortex | 3.93 | 36 | -12 | -4 |
| **8** | R Thalamus | 3.76 | 18 | -34 | 2 |
| **8** | L Planum Temporale | 3.59 | -58 | -18 | 2 |
| **8** | R Frontal Pole | 3.57 | 40 | 62 | 2 |
| **8** | L Thalamus | 3.71 | -6 | -2 | 4 |
| **8** | R Insular Cortex | 3.61 | 34 | 10 | 4 |
| **8** | R Insular Cortex | 3.64 | 32 | 12 | 10 |
| **8** | R Frontal Pole | 3.59 | 22 | 52 | 20 |
| **8** | R Middle Frontal Gyrus | 3.59 | 44 | 20 | 28 |
| **8** | L Middle Frontal Gyrus | 3.61 | -40 | 28 | 34 |
| **8** | R Precuneus Cortex | 4.02 | 16 | -66 | 38 |
| **8** | R Cingulate Gyrus, anterior division | 3.64 | 4 | 4 | 38 |
| **8** | L Cingulate Gyrus, anterior division | 3.75 | -6 | 4 | 38 |
| **8** | R Juxtapositional Lobule Cortex (formerly Supplementary Motor Cortex) | 3.63 | 8 | -4 | 52 |
| **8** | R Superior Frontal Gyrus | 3.82 | 14 | 28 | 52 |
| **8** | R Precentral Gyrus | 3.82 | 22 | -16 | 64 |
| **8** | R Precentral Gyrus | 3.73 | 14 | -20 | 76 |
